# Contextual inference underlies the learning of sensorimotor repertoires

**DOI:** 10.1101/2020.11.23.394320

**Authors:** James B. Heald, Máté Lengyel, Daniel M. Wolpert

## Abstract

Humans spend a lifetime learning, storing and refining a repertoire of motor memories. However, it is unknown what principle underlies the way our continuous stream of sensorimotor experience is segmented into separate memories and how we adapt and use this growing repertoire. Here we develop a principled theory of motor learning based on the key insight that memory creation, updating, and expression are all controlled by a single computation – contextual inference. Unlike dominant theories of single-context learning, our repertoire-learning model accounts for key features of motor learning that had no unified explanation and predicts novel phenomena, which we confirm experimentally. These results suggest that contextual inference is the key principle underlying how a diverse set of experiences is reflected in motor behavior.

---

Throughout our lives, we experience different contexts, in which the environment exhibits distinct dynamical properties, such as when manipulating different objects or walking on different surfaces. Although it has been recognized that the brain maintains multiple motor memories appropriate for these contexts ^1,2^, classical theories of motor learning have focused on how the brain adapts to a single type of environmental dynamics ^3–5^. However, with multiple memories come new computational challenges: the brain must decide when to create new memories, and how much to express and update them for each movement we make. The principles underlying these operations are poorly understood, and therefore it is unclear what aspects of motor learning are fundamentally determined by them. Here, we propose a unifying principle – contextual inference – that specifies how sensory cues and movement feedback affect memory creation, expression and learning. We show that contextual inference is the core feature that underlies a range of fundamental aspects of motor learning that previously could only be explained by proposing a number of distinct and often heuristic processes.

## COIN model: the three contributions of contextual inference

We developed the COIN (COntextual INference) model, a principled Bayesian model of learning a motor repertoire in which separate memories are stored for different contexts (see Suppl. Mat. & Fig. S1). Each memory stores information learned about the dynamical and sensory properties of the environment associated with the corresponding context. Crucially, neither contexts nor their transitions come labeled as such, and thus a major challenge for the learner is to continually infer which context they are in, based on a continuous stream of experience (Fig. 1a). In the COIN model, contextual inference fuses information from multiple sources: prior expectations about which context the learner is in, based on the history of contexts inferred so far (the overall occurrence probability of each context as well as the transition probabilities between them); and the probability that current state feedback (the sensory consequences of motor commands; Fig. 1a, purple) and sensory cues (sensory input that does not depend on action, such as the visual appearance of an object; Fig. 1a, green and yellow) are generated by each context. The result of contextual inference is a posterior distribution expressing the probability with which each known context, or a yet-unknown novel context, is active at that time. In turn, contextual inference determines memory creation, expression and updating. A new memory is created whenever the probability of a novel context becomes high (Fig. 1b). For determining the current motor command (Fig. 1e), rather than selecting a single memory to be expressed ^2,6^, the state associated with each memory (Fig. 1c) is expressed commensurate with the probability of the corresponding context under the posterior computed before experiencing state feedback (its “predicted probability”; Fig. 1d). Therefore, unlike in traditional models of motor learning, slow adaptation to a sudden change of environmental dynamics (Fig. 1e, red arrow) may not arise from a slow updating of the state of any individual memory (Fig. 1c, all are relatively constant during this period) but from the slow updating of the inferred context probabilities (Fig. 1d, red arrow), which determine the extent to which the inferred states are expressed in the motor output. Once a movement has been executed, each memory is updated according to the probability of the corresponding context under the posterior given the state feedback received (its “responsibilty”; Fig. 1f; numbered arrows 1 and 2 respectively show how high and low responsibility for the red context speeds up and slows down the updating of its state, Fig. 1c). Therefore, the COIN model, unlike traditional models, proposes that contextual inference is core to motor learning.

**Fig. 1.**
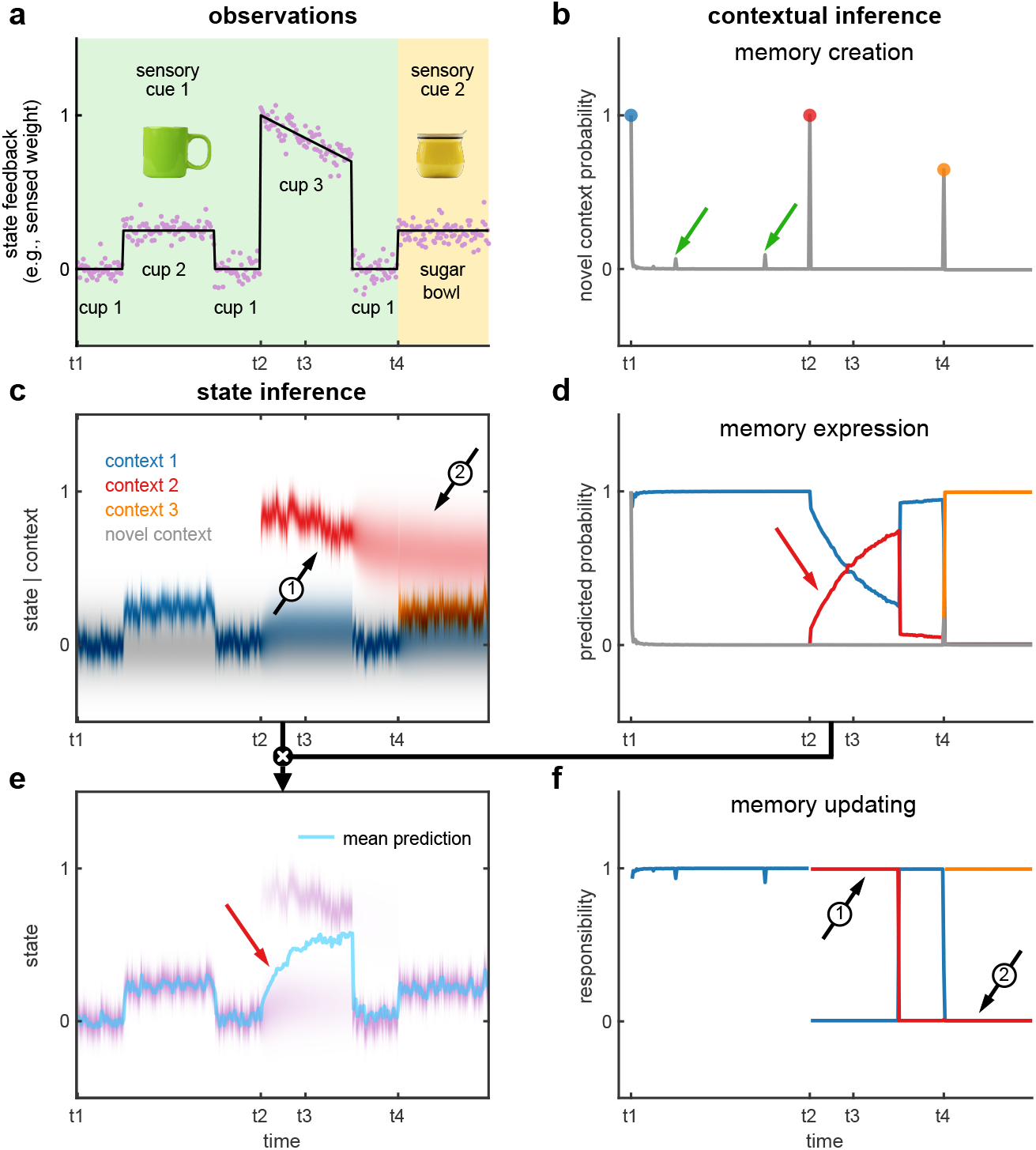
Contributions of contextual inference to motor learning in the COIN model. **a,** An example of experience (state feedback and sensory cues) when handling a series of visually-identical cups of different weights followed by a sugar bowl. The state feedback is noisy observations (purple dots) of the true weight (black line) shown on an arbitrary scale. Background color indicates different sensory cues (visual appearance of cups vs. sugar bowl). **(b-f)** The COIN model applied to the observations in **a**(see also Fig. S2). **b,** Probability that a novel context generated the observations. The colored circles shows memory creation events. When transitioning to and from cup 2 the novel context probability increases (green arrows) but is insufficient to generate a new memory. **c,** Predicted state distributions of the three contexts inferred by the model. Grey shows the prior distribution for a novel context. **d,** Predicted probability (before state feedback is observed) of each instantiated context as well as a novel context (grey). **e,** Predicted state distribution (purple), which is a mixture of the individual contexts’ predicted state distributions (**c**), weighted by their predicted probabilities (**d**). The motor output is the mean of the predicted state distribution (cyan line). **f,** Responsibility (context probability after observing state feedback) of each instantiated context as well as a novel context (grey). Parameters for the COIN model simulation are shown in Table S1 and validation of the COIN model inference in Figs. S3 and S4. See text for explanation of arrows in panels c-f.

## Memory creation and expression accounts for spontaneous and evoked recovery

As an ideal litmus test of the first two contributions of contextual inference to learning a motor repertoire, memory creation and expression (Fig. 1b and d), we revisited a widely-used spontaneous recovery paradigm. In this paradigm (Fig. 2b, top left), participants learn a perturbation *P*^+^ (Fig. 2a) applied by a robotic interface while reaching to a target, which is followed by brief exposure to the opposite perturbation *P*^−^, bringing performance back to baseline. Adaptation is measured during learning using occasional channel trials, *P*^c^, in which the robot constrains the movement so as to produce no error (Fig. 2a, see Methods for details). After the *P*^−^ phase, adaptation is assessed during a long and continuous series of channel trials. As in previous studies, our participants showed the intriguing feature of spontaneous recovery in this phase (Fig. 2c): a transient re-expression of *P*^+^ adaptation, rather than a simple decay towards baseline.

**Fig. 2.**
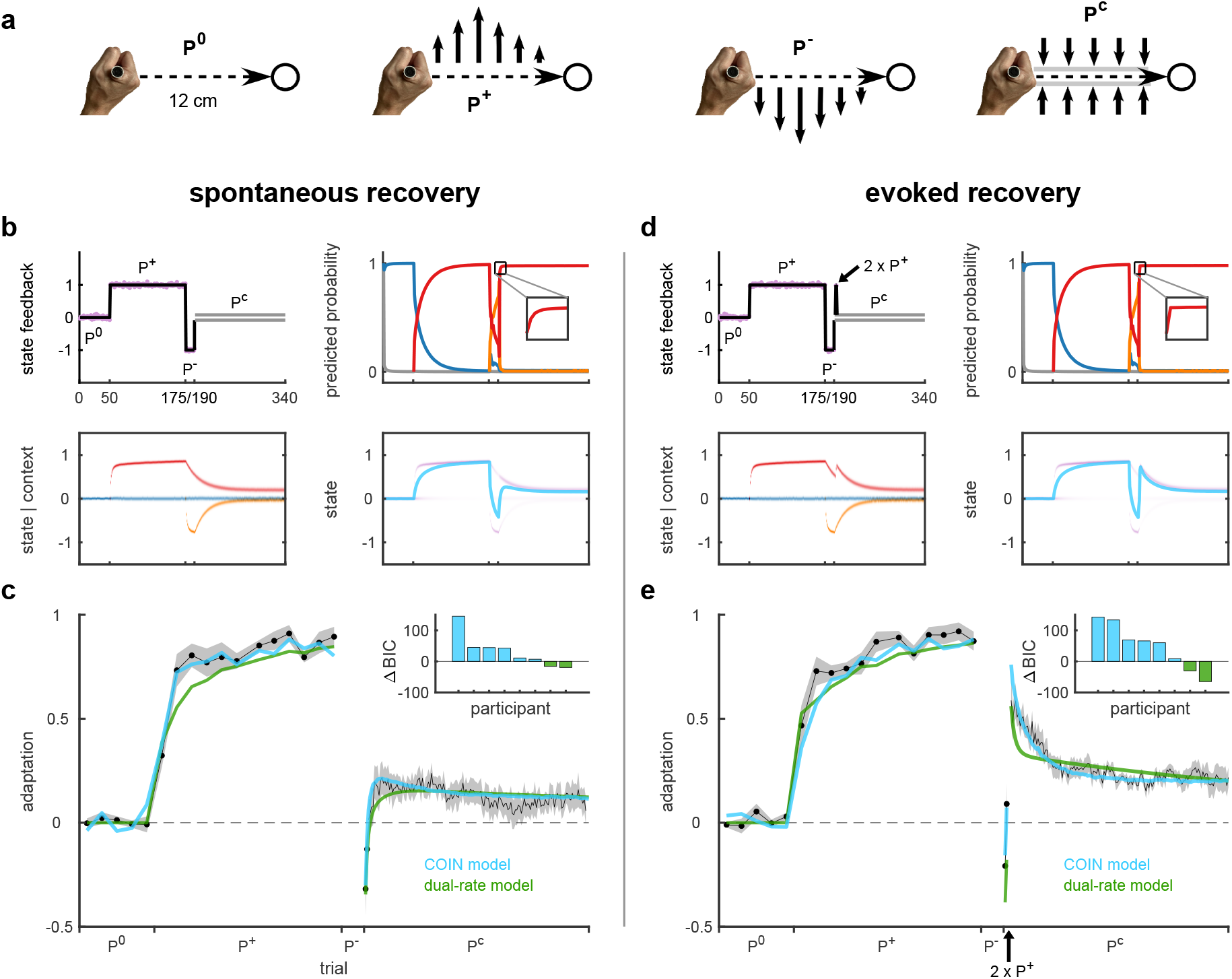
Memory creation and expression accounts for spontaneous and evoked recovery. **a,** Participants made reaching movements (dashed arrow) to a target (circle) while holding the handle of a vBOT robotic manipulandum ^7^ that could generate forces (black arrows) at the hand. The vBOT could either be passive (null field, *P*^0^) or generate a velocity-dependent force field that acted to the left (*P*^+^) or right (*P^−^*) of the current movement direction. Channel trials (*P*^c^) were used to assess adaptation by constraining the hand (with a stiff one-dimensional spring) to a straightline channel (grey lines) to the target and measuring the forces generated by the participant into the channel walls. **b,** Simulation of the spontaneous recovery paradigm with the COIN model. Top left: ground-truth perturbation (black line) and noisy state feedback observations (purple), as in Fig. 1a, with grey parallel lines showing channel phase. Bottom left: predicted state distributions of inferred contexts as in Fig. 1c. Top right: predicted probability of inferred contexts as in Fig. 1d. Inset shows magnification of beginning of channel phase (trials 194-203). Bottom right: predicted state distribution (purple) and mean predicted state (cyan), as in Fig. 1e. **c,** Mean adaptation (black, ± s.e.m. across participants) on the channel trials of the spontaneous recovery paradigm. The cyan and green lines show model fits (mean of individual participant fits) of the COIN and dual-rate models, respectively. Inset shows ΔBIC for individual participants, positive favors the COIN model. **d-e**, As in (**b-c**) for the evoked recovery paradigm in which the 3^rd^ and 4^th^ trials in the channel phase were replaced by *P*^+^ trials (black arrow). For COIN model parameter and model recovery (COIN vs. dual-rate) see Suppl. Mat. and Figs. S9 and S10.

Although there are no explicit sensory cues in this paradigm, according to our theory, contextual inference continues to play an important role even in the absence of such cues. We simulated the COIN model for the spontaneous recovery paradigm (Fig. 2b, using parameters in Table S1, see Methods). Starting with a memory appropriate for moving in the absence of a perturbation (*P*^0^, blue Fig. 2b, bottom left), new memories were created for the *P*^+^ (red) and *P*^−^ (orange) perturbations. Spontaneous recovery arises in the COIN model due to the dynamics of contextual inference. As *P*^+^ has been experienced for a longer time overall, it is quickly inferred to be active with a much higher probability during the *P*^c^ phase (Fig. 2b, top right). Therefore, while *P*^+^ is inferred to be active but its state has not yet decayed (Fig. 2b bottom left), the memory of *P*^+^ is transiently expressed in the subject’s motor command (Fig. 2b bottom right). Our mathematical analysis also confirmed that spontaneous recovery was a robust property of the COIN model (Suppl. Text and Fig. S5).

Importantly, the mechanism of spontaneous recovery is fundamentally different in the COIN model from that proposed by classical, single-context models of motor learning such as the dual-rate model ^3^. Critically, in those models, motor output is determined by a combination of individually updating memories whose expression does not change over time. Thus, the dynamics of adaptation is solely determined by the dynamics of memory updating. In contrast, in the COIN model, changes in motor output can occur without updating any individual memory, simply due to the re-expression of a memory if sensory-motor evidence indicates sufficiently strongly that a change in context has occurred. Therefore, we designed a novel “evoked recovery” paradigm in which two early trials in the channel trial phase of the spontaneous recovery paradigm were replaced with *P*^+^ (‘evoker’) trials (Fig. 2d, top left, akin to trigger trials in visuomotor learning ^2^). In this case, the COIN model predicts a strong and long-lasting recovery of *P*^+^-adapted behavior (Fig. 2d, bottom right), primarily due to the almost instantaneous inference that the *P*^+^ context is now active (Fig. 2d, top right, red) and the gradual decay of the *P*^+^ state over subsequent channel trials (Fig. 2d, bottom left, red). Our mathematical analysis suggested that this was also a robust prediction of the COIN model (Suppl. Text and Fig. S5). In contrast, the dual-rate model only predicts a transient recovery that rapidly decays due to the same underlying adaptation process with fast dynamics governing both recovery and decay (Fig. S7). In line with the predictions of the COIN model, participants showed a strong evoked recovery in response to the *P*^+^ trials (Fig. 2e). This recovery lasted for the duration of the experiment, defying models that predict a simple exponential decay to baseline ^2,8,9^ (Fig. S6 and Table S3). We fit both the COIN and dual-rate models to individual participants’ data in the spontaneous recovery and evoked recovery groups (Fig. 2c & e; Tables S1 and S2). The COIN model was able to fit the data accurately, but the dual-rate model (and its multi-rate extensions, Fig. S7) showed a qualitative mismatch in the time course of decay of evoked recovery (Fig. 2e). Formal model comparison with BIC ^10^ (insets in Fig. 2c & e) also provided strong support for the COIN model overall (Δ group-level BIC of 302.6 and 394.1 for the spontaneous and evoked recovery groups, respectively) and for the majority of participants individually (6 out of 8 for each experiment; individual fits shown in Fig. S8).

## Memory updating depends on contextual inference

According to the COIN model, the third contribution of contextual inference to learning a motor repertoire is controlling how each existing memory is updated following experience based on their respective inferred responsibilities (Fig. 1f), which depend on sensory cues and state feedback. In order to test this prediction, we conducted an experiment in which participants experienced a training phase in which two arbitrary sensory cues (the appearance of a visually defined target; Fig. 3a) were consistently paired with two perturbations (i.e. 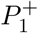 and 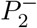 trials, subscript specifies sensory cue; Fig. 3b). In a subsequent test phase using triplet trials ^9,12^ (Fig. 3c), we measured how the memory associated with cue 1 was updated based on exposure to a single trial with a cue-perturbation combination that was either consistent 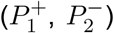 or inconsistent 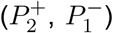 with those experienced during training. We fit the COIN model to the data of individual participants (Table S1). Under such conditions, in the COIN model, the responsibility of each context *c* for the perturbation experienced on the exposure trial is a product of two terms. The first term is the prior, i.e. the predicted probability with which the context was expected before experiencing the perturbation (as in Fig. 1d) – which is already conditioned on the cue visible from the outset of the trial, P(*c*|*q*). The second term expresses the likelihood of the state feedback in that context, P(*y*|*c*). This simple principle explains the intricate pattern of single-trial learning exhibited by our participants: relatively uniform updating across conditions before learning (Fig. 3d) and graded updating after learning, with maximal (minimal) updating in the 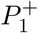-direction for consistent 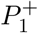 (inconsistent 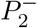) trials (Fig. 3e) – as well as how this pattern changed during the training phase (Fig. S12).

**Fig. 3.**
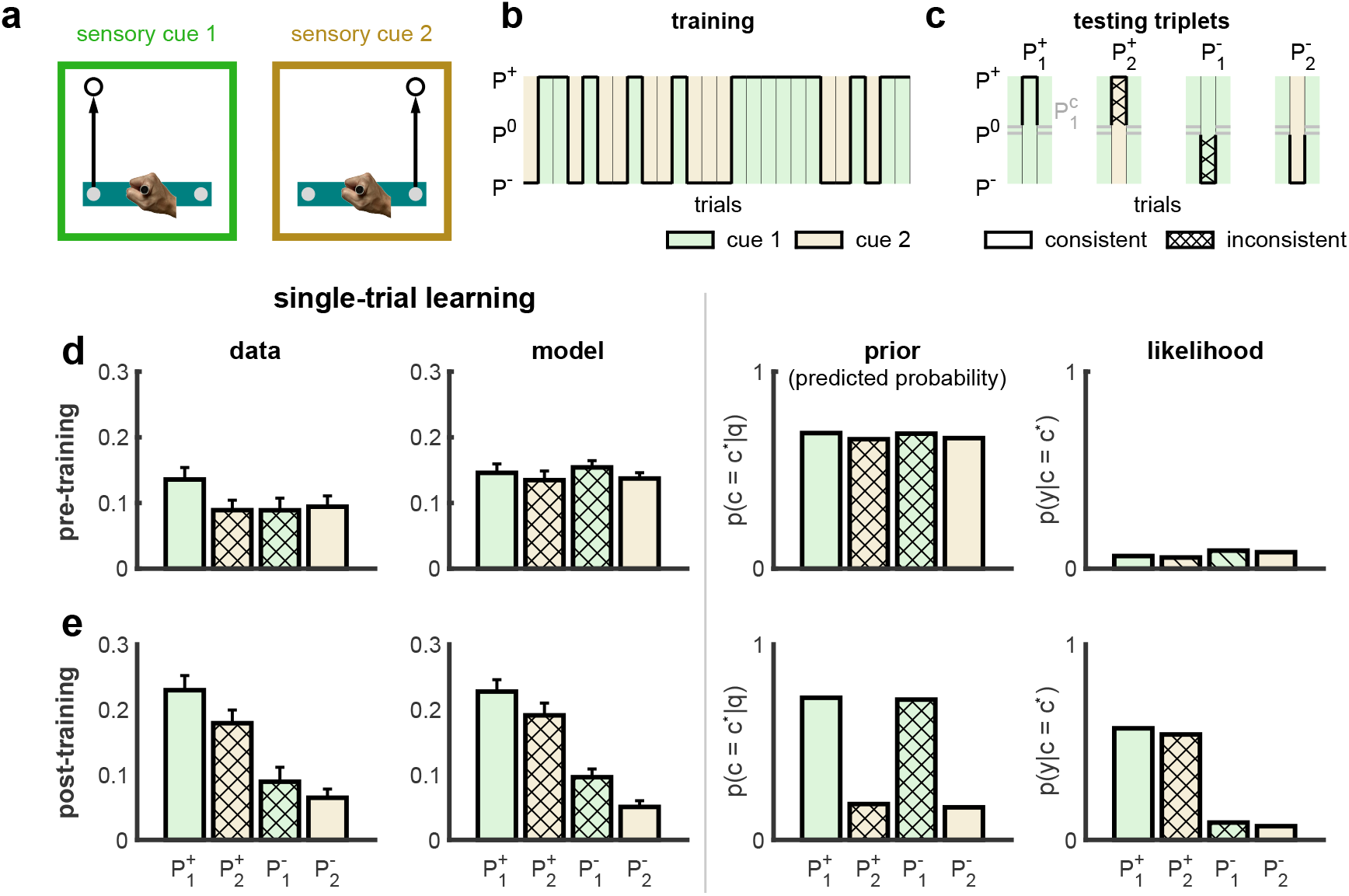
Memory updating depends on contextual inference. **a,** Experimental paradigm: sensory cues are defined by the location of a target (circle) to which participants had to move the corresponding control point (gray disks) on a virtual bar (horizontal rectangle) ^11^. **b,** Training: sensory cues (indicated by background color) are consistently matched to perturbations (black line) randomly selected on each trial. **c,** Triplets used to examine single-trial learning: two channel trials (both with cue 1, 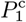) bracket an ‘exposure’ trial (one of the four combinations of sensory cue and perturbation). **d-e,** Single-trial learning, measured as the difference in adaptation expressed between the two channel trials of a triplet, for the four cue-perturbation combinations before (**d**) and after (**e**) the training phase. Experimental data shows mean ± s.e.m. across participants (left). Examining the differences in single-trial learning across the four cue-perturbations combinations (repeated-measures ANOVA) showed no significance before training (*F*_3,69_ = 1.87, *P* = 0.142) but a significant difference after training (*F*_3,69_ = 15.24, *P* = 1 × 10^*−*7^). The COIN model was fit to each participant and model predictions are shown as mean ± s.e.m. across participants (center-left). A positive value indicates a change in adaptation consistent with the perturbation experienced on the exposure trial (an increase following *P*^+^ and a decrease following *P^−^*). The model prior (center-right) and (normalized) likelihood (right) are also shown for the context that was predominantly expressed on the final channel trial of the triplet (*c*\*, which is the context associated with *P*^0^ before training, and *P*^+^ after training). The combination of the prior and likelihood produce the posterior (responsibilities). Using an analytic approximation, it is possible to show that the posterior is proportional to single-trial learning (see Suppl. Text and Fig. S11). Between the two inconsistent conditions post-training, the model accounts for the greater single-trial learning seen in 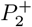 compared to 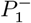 by developing an internal model that believes state feedback (determining the likelihood) is more strongly associated with contexts (force fields) than the sensory cues (determining the prior).

## Contextual inference underlies apparent changes in learning rates

Finally, we revisited classical results about apparent changes in learning rates under a variety of conditions, each of which required different explanations. What is common in all these cases is that the empirical finding of temporal (trial-to-trial) changes in adaptation has been interpreted as learning, i.e. changes to any existing memories (states), and thus differences between the magnitudes of these changes have been interpreted as differences in learning rates. The COIN model suggested that changes in adaptation can occur without learning, simply by changes in the way existing memories are expressed. Therefore, apparent changes in learning rates may be due to changes in memory expression rather than the memories themselves. To test this hypothesis, we used the parameters we had obtained by fitting each of the 40 participants in the experiments described above (Table S1 and Fig. S13), and thus the simulations provide parameter-free predictions. Fig. 4 shows three classical paradigms (column 1) with representative extant experimental data (column 2) and COIN model predictions (column 3), as well as the key internal components of the COIN model, state (column 4) and contextual inferences (column 5).

**Fig. 4.**
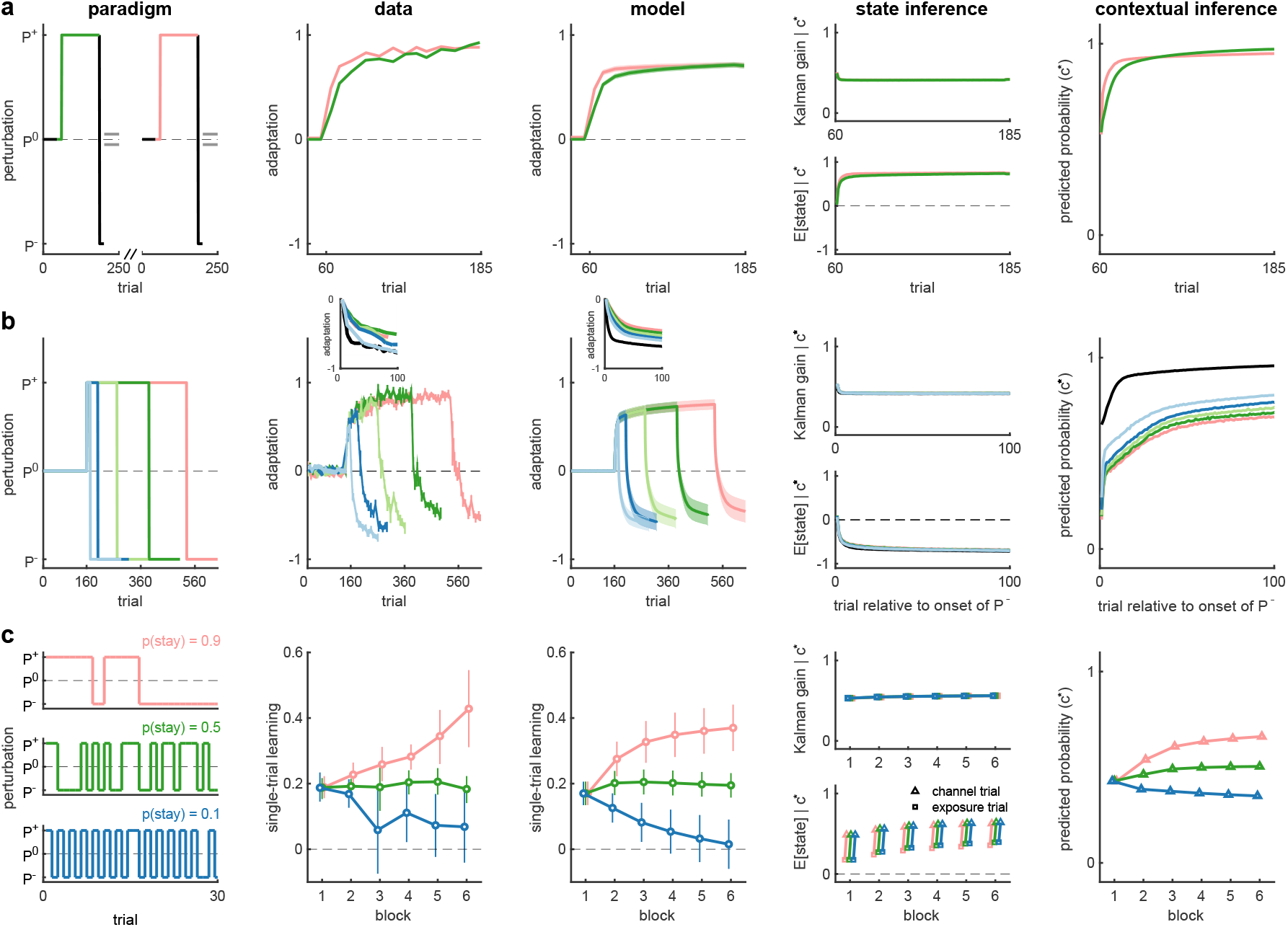
Contextual inference underlies apparent changes in the learning rate. The COIN model applied to three phenomena: savings (**a**), anterograde interference (**b**) and the effect of environmental consistency (**c**). Column 1: experimental paradigms (lines as in previous figures, colors highlight key comparisons). Column 2: experimental data replotted from Ref. 13 (**a**), Ref. 14 (**b**) and Ref. 9 (**c**). Column 3: output of COIN model averaged over 40 parameter sets obtained from fits to individual participants in the experiments shown in previous figures. Error bars show s.e.m. based on the number of participants in the original experiments. Columns 4-5: internal inferences in the COIN model with regard to the context (*c*\*) that is most relevant to the perturbation to which adaptation is measured. Specifically, *c*\* is the context with the highest responsibility on the given trial (that associated with *P*^+^ in **a** and *P^−^* in **b**) or, as in Fig. 3d (also single-trial learning), the context with the highest predicted probability on the second channel trial of a triplet (that associated with *P*^+^, **c**). Column 4: Kalman gain (top) and mean predicted state (bottom) for the relevant context *c*\*. Column 5: Predicted probability of the relevant context *c*\*. Insets in **b**(columns 2-3) show data aligned at the point at which adaptation crosses zero during the *P^−^* phase for ease of comparison. Black lines in **b** represent initial adaptation to *P*^+^ and have been sign inverted in columns 2-3 and the bottom panel of column 4. Data in **c** shows averages within blocks, with the bottom panel in column 4 showing separate averages for exposure and subsequent channel trials.

The COIN model showed savings (Fig. 4a): learning the same perturbation a second time (even after washout) is faster than the first time (e.g. Refs. 3,13,15,16). In contrast, linear time-invariant state space models (such as the dual-rate model) cannot explain savings after washout ^17^. Savings in the COIN model is a natural consequence of the dynamics of contextual inference: *P*^+^ is expected with higher probability during the second exposure after having experienced it during the first exposure (Fig. 4a, contextual inference, compare pink and green). Conversely, the Kalman gain, i.e. the learning rate for the state (see Suppl. Mat.), remained unchanged between the two exposures (Fig. 4a, state inference, top panel, pink and green overlapping).

The COIN model also shows anterograde interference (Fig. 4b) in which learning a perturbation (*P*^−^) is slower if an opposite perturbation (*P*^+^) has been previously learned (e.g. Refs. 14,18), with the amount of interference increasing with the length of experience of the first perturbation (Fig. 4b, data). Critically, this slowness is not due to simply starting from an oppositely adapted state, as it persists even if the speed of adaptation is measured only from the point when adaptation passes through zero (Fig. 4b, data, inset). The COIN model reproduces the interference (Fig. 4b, model) because more extended experience with *P*^+^ makes it less probable that a transition to any other inferred context, including *c*^−^ (the context associated with *P*^−^), might happen (Fig. 4b, contextual inference), thereby decreasing the expression of the state of *c*^−^, and thus slowing apparent adaptation to *P*^−^. Again, state inference alone did not predict a difference between conditions (Fig. 4b, state inference).

The persistence of the environment has also been shown to affect single-trial learning (Fig. 4c, paradigm and data) ^9,12^. More consistent environments (Fig. 4c, pink) lead to an increase in single-trial learning compared to more variable environments (Fig. 4c, blue). Again, in contrast to classical interpretations that are based on the learning rate of the state being modified by environmental consistency ^9^, the COIN model reproduces these results (Fig. 4c, model) without changes in the Kalman gain (Fig. 4c, state inference, top). Specifically, the extent of memory updating is independent of consistency (Fig. 4c, state inference, bottom), and instead it is the expression of memories that differs across conditions: more (less) consistent perturbations lead to higher (lower) probabilities with which the model predicts contexts to persist from one trial to the next (Fig. 4c, contextual inference). The COIN model was also able to explain changes in single-trial learning across a number of different experimental paradigms that varied environmental consistency in Gonzalez Castro et al. ^12^ (Fig. S14).

Importantly, these results together with those presented in Fig. 3 and Fig. S11 suggest that what is typically considered as “single-trial learning” is in fact a mixture of two processes (see Suppl. Text): the updating of memories on the exposure trial, which can be considered genuine learning as it involves a change in inferred state(s), and the expression of those states in the subsequent trial (based on contextual inference).

## Cognitive processes underlying contextual inference

In addition to providing a comprehensive account of the phenomenology of motor learning, the COIN model also suggests how specific cognitive mechanisms contribute to the underlying computations. For example, associating working memory with the maintenance and updating of context probabilities explains why and how a working memory task can effectively lead to evoked recovery in a modified version of the spontaneous recovery paradigm ^19^ (see Suppl. Text and Fig. S15). Furthermore, identifying explicit and implicit forms of learning with state and specific parameter inferences in the model, respectively, explains the complex time courses of explicit and implicit components of visuomotor learning ^20–22^ (see Suppl. Text and Fig. S16).

## Discussion

In summary, the COIN model puts the problem of learning a repertoire of memories, rather than individual memories, center stage. Once this more general problem is considered, contextual inference becomes a key computation that unifies seemingly disparate data sets, such as spontaneous and evoked recovery, the interplay of sensory cues and state feedback in memory updating, savings, anterograde interference and apparent changes in learning rate with environmental consistency. In comparison, previous models either lack a notion of multiple contexts altogether ^3,9^, or contextual inference and its effects on memory creation, updating or expression remain partially heuristic, lacking the principles of the COIN model ^2,6,8,23^. As a consequence, these models can only account for a subset of these results (Table S3), which they were often hand-tailored to address. Specifically, models postulating multiple, simultaneous learning processes with different, but fixed rates can explain spontaneous, but not evoked recovery, the effects of environmental consistency, or our memory updating experiment ^3,8^. Conversely, previous models that include the possibility of multiple contexts (or modules) fail to account for both spontaneous and evoked recovery ^1,2,6^. Models assuming a single process (in a single context) with dynamically changing rates do not account for either spontaneous / evoked recovery or memory updating ^9^.

While creating new memories in the COIN model is essential for learning a motor repertoire, reorganization of memories may also be necessary ^24^. In contrast to the online processes we studied here (memory creation, updating, and expression), the COIN model could be extended, for example, by the offline pruning or merging of existing memories, during sleep or periods of inactivity.

Previous work exploring humans’ complex and hierarchical internal models have typically studied higher-level (e.g. conceptual ^25,26^ or causal ^27^) forms of cognition and characterized learning either on a developmental time scale ^25^, or just through its end result ^26,27^. Crucially, our results demonstrate that the motor system provides an ideal opportunity to track the dynamical emergence of such internal models over the course of learning, even within a single session, by performing detailed subject-by-subject and trial-by-trial analyses under controlled perturbations. We suggest that these results will be relevant to all forms of learning in which experience can be usefully broken down into discrete contexts – in the motor system and beyond.

## Acknowledgments

This work was supported by the European Research Council (ERC) under the European Union’s Horizon 2020 research and innovation programme (grant agreement No 726090 to M.L.) and the Wellcome Trust (Investigator Awards 212262/Z/18/Z to M.L and 097803/Z/11/Z to D.M.W), Royal Society (Noreen Murray Professorship in Neurobiology to D.M.W), Engineering and Physical Sciences Research Council (studentship to J.B.H). We thank J. Ingram for technical support. Authors’ contributions: J.B.H developed the model, implemented the model, performed the experiments, analyzed the data and performed simulations. J.B.H and D.M.W designed the behavioral experiments. All authors were involved in the conceptualization of the study, developed techniques for analyzing the model, interpreted results and wrote the paper. The authors have no competing interests. All data, code, and materials used in the analysis will be available in a public repository to any researcher for purposes of reproducing or extending the analysis.

## Supplementary materials

## Materials and Methods

### Participants

A total of 40 neurologically-healthy participants (18 males and 22 females; age 27.7 ± 5.6 yr, mean ± s.d.) were recruited to participate in two experiments, which had been approved by the Cambridge Psychology Research Ethics Committee and the Columbia University IRB (AAAR9148). All participants provided written informed consent and were right-handed according to the Edinburgh handedness inventory ^28^. To provide sufficient power, sample sizes were chosen on the basis of the typical between-participant variability observed in similar motor adaptation studies (e.g. 3,9,11,21). Note that our analyses were performed both at a single participant level as well as a group level.

### Experimental apparatus

All experiments were performed using a vBOT planar robotic manipulandum with virtual-reality system and air table ^7^. The vBOT is a modular, general-purpose, two-dimensional planar manipulandum optimized for dynamic learning paradigms. The vBOT’s handle position was measured using optical encoders sampled at 1 kHz and torque motors allowed forces to be generated at the handle and updated at the same rate. Participants grasped the handle of the manipulandum with their right hand while their forearm was supported on an air sled, which constrained arm movements to the horizontal plane and reduced friction. A monitor mounted horizontally face-down above the vBOT projected images via a horizontal mirror so that visual feedback was overlaid in the plane of movement. In the spontaneous/evoked recovery experiment, the mirror prevented direct vision of the hand and forearm. In the memory updating experiment, a semi-silvered mirror was used and a lamp illuminated the hand from below the mirror with the illumination adjusted so that both the vBOT, hand, arm and virtual images were clearly visible. This was done to ensure that participants had an accurate estimate of the state of their hand and arm (as in Ref 11).

On each trial, the vBOT could either generate no forces (*P*^0^, null field), a velocity-dependent curl force field (*P*^+^ or *P*^−^ perturbation depending on the direction of the field) or a force channel (*P*^c^, channel trials). For the curl force field, the force generated on the hand was given by

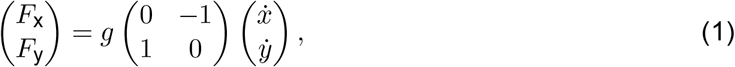

where *F*_x_, *F*_y_, 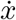 and 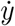 are the forces and velocities at the handle in the *x* (transverse) and *y* (sagittal) directions respectively. The gain *g* was set to ± 15 Ns m^−1^, where the sign of *g* specified the direction of the curl field (counterclockwise or clockwise which were assigned to *P*^+^ and *P*^−^, counterbalanced across participants). On channel trials, the hand was constrained to move along a straight line from the home position to the target. This was achieved by simulating forces associated with a stiff spring and damper, with the forces acting perpendicular to the long axis of the channel. A spring constant of 3,000 N·m^−1^ and a damping coefficient of 140 Ns m^−1^ were used. Channel trials clamped the kinematic error close to zero and were used to measure the participant’s level of adaptation to the *P*^+^ and *P*^−^ perturbations based on the forces they generated into the channel walls ^29,30^.

### Experiment 1: Spontaneous and evoked recovery

Sixteen participants either performed a spontaneous or an evoked recovery condition. In both conditions participants made repeated movements of a cursor, always aligned with the center of the handle, from a home position to a target. On each trial participants first aligned the cursor (blue cm radius disc) with a home position (0.5 cm radius circle) situated in the midline approximately 30 cm in front of the participant’s chest. The trial started when the cursor was within 0.5 cm of the home position and had remained below a speed of 0.5 cm·s^−1^ for 0.1 s. After a 0.3 s delay, a target (0.5 cm radius circle) appeared 12 cm in front of the home position (distally in the *y* direction) and a tone indicated that the participants should initiate a reaching movement to the target. The trial ended when the cursor had remained within 0.5 cm of the target and below a speed of 0.5 cm·s^−1^ for 0.1 s. If the peak speed of the movement was less than 50 cm·s^−1^ or more than 70 cm·s^−1^, a low-pitch tone sounded and a ‘too slow’ or ‘too fast’ message was displayed, respectively. At the end of each trial the participant relaxed and the vBOT actively returned the hand to the home position.

### Spontaneous recovery condition

In the spontaneous recovery condition, participants (n = 8) performed a version of the standard spontaneous recovery paradigm ^3^. The paradigm consisted of a pre-exposure phase (5 blocks/50 trials) with a null field (*P*^0^). This was followed by an exposure phase (12 blocks/120 trials, with an additional 5 exposure trials after the 45 s rest break given after block 6) with *P*^+^ (the direction of the force field assigned to *P*^+^ was counterbalanced across participants). In the pre-exposure and exposure phases, to assess adaptation each block of 10 trials had one channel trial (*P*^c^) in a random location (not the first). After the exposure phase, participants were rapidly de-adapted in a counter-exposure phase by applying 15 trials with the opposite perturbation (*P*^−^). This was followed by a long series of 150 channel trials (*P*^c^).

### Evoked recovery condition

Participants (n = 8) in the evoked recovery condition performed a modified version of the spontaneous recovery paradigm which differed in that the 3^rd^ and 4^th^ trials of the channel phase were replaced with *P*^+^ trials (Fig. 2d).

### Experiment 2: Memory updating

This experiment is based on a paradigm in which sensory cues allow multiple memories to be simultaneously learned ^11^. Participants (n = 24) grasped the handle of the vBOT and a virtual object (solid green rectangle, 16 × 3 cm) was displayed centred on the hand and translated with hand movements Fig. 3a). The object had two potential control points (blue 0.4 cm radius discs) ± 7 cm lateral to the centre of the object. On each trial, participants first aligned the centre of the object (indicted by a yellow cross) with the home position (0.5 cm radius circle) situated in the midline approximately 30 cm in front of the participant’s chest.

The trial started after the center of the object was within 0.5 cm of the home position and had remained below a speed of 0.5 cm·s^−1^ for 0.1 s. After a 0.3 s delay, a target (a circle with a radius of 0.5 cm) appeared 12 cm away (distally along the *y* axis) in front of either the left or right control point. A tone indicated that the participants should initiate a reaching movement to the target. Participants were instructed to move the corresponding control point to the target. That is, if the target was aligned with the left control point, they should move the left control point to the target, and conversely for the target aligned with the right control point. Crucially, because each target was aligned with its respective control point, the hand had to move to the same location to attain either target. The different targets and control points provide different sensory cues for the trial. The trial ended when the control point had remained within 0.5 cm of the target for 0.1 s below a speed of 0.5 cm·s^−1^. If the peak speed of the movement was less than 50 cm·s^−1^ or more than 70 cm·s^−1^, a low-pitch tone sounded and a ‘too slow’ or ‘too fast’ message was displayed, respectively. At the end of each trial, the vBOT actively returned the hand to the home position

Each trial could be made in a null field (*P*^0^), in one of the two perturbations (*P*^+^ or *P*^−^) or in a channel (*P*^c^) and with either the left or right sensory cue (target-control point). We indicate the sensory cue used on a trial by a subscript (e.g. 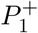 and 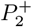 for the *P*^+^ perturbation with the left and right sensory cue, respectively). The experiment consisted of three phases: pre-training, training and post-training.

The training phase (see details below) consisted or exposure to two cue-perturbation pairs (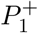 and 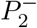) so that participants could be expected to associate each cue with its corresponding perturbation.

In the pre-training and post-training phases (Fig. 3c) participants performed blocks of trials which consisted of 8, 10 or 12 *P*^0^ trials with an equal number of each sensory cue in a pseudorandom order (for post-exposure phase the number of *P*^0^ trials was reduced to 2, 4 or 6). These trials were used to bring adaptation close to baseline so that we could assess single-trial learning in the triplet of trials that followed the *P*^0^ trials. The first and third trial in the triple were always channel trials with sensory cue 1 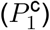 and the middle trial of the triplet (‘exposure’ trial) was one of the four possible combinations of perturbation (force-field direction) and sensory cue (control point): 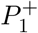, 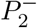, 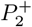 and 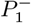. Therefore the first two exposure trial types (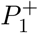 and 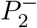) were the same as those experienced in the training phase and we refer to them as ‘consistent’ and the latter two (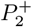 and 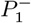) are ‘inconsistent’ with the training as they match a cue with a perturbation not experienced during training. Within each sequence of 4 blocks, each of these combinations was experienced once and the four blocks were repeated 4 times in pre-exposure and 8 times in post-exposure. Importantly, there was no consistent relationship between sensory cues and perturbations in the pre-training phase, as each triplet type was presented an equal number of times.

In the training phase (Fig. 3b), each sensory cue was consistently and repeatedly associated with one perturbation (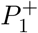 and 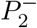) and single-trial learning was occasionally assessed using consistent exposure trials only. To do this participants performed 24 blocks of trials consisting of:

- 2 channel trials (one 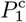 and 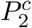 order counterbalanced across consecutive blocks)
- 32 force-field trials (equal number of 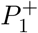 and 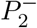 within each 8 trials in a pseudorandom order)
- 2 channel trials (one 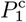 and 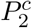) order reversed from first two channel trials to assess adaptation after learning
- 14, 16 or 18 washout trials with (equal number of 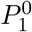 and 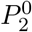 in a pseudorandom order)
- 1 triplet (with exposure either 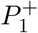 or 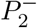 with order counterbalanced across consecutive blocks)
- 6, 8 or 10 washout-field trials (sampled without replacement)
- 1 triplet (exposure of 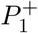 or 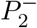 that was not experienced on the previous triplet).

When the number of null-field trials could vary we sampled without replacement from the options and replenished whenever they emptied. The order in which each triplet type was presented in a block was counterbalanced across blocks. A 60 s rest break was given after every 3 blocks during the training phase. After each rest break, 8 null-field trials were performed in which the sensory cues were presented in a pseudorandom order.

The control point assigned to sensory cue 1 (used on all triplet channel trials) and sensory cue 2 was counterbalanced across participants as was the direction of force field assigned to *P*^+^ and *P*^−^.

Prior to the experiment participants performed a familiarization phase of 80 trials consisting of null-field trials and channel trials for each contextual cue in a pseudorandom order.

### Data analysis

On channel trials, we calculated adaptation as the proportion of the force field that was compensated for. This was taken as the slope of the regression (with zero offset) of the time series of actual (signed) force generated into the channel walls against the time series of forces (based on the hand velocity in the channel) that would fully compensate for the perturbation had it been present ^3^. For this analysis, we used the portion of the movement where the hand velocity was greater than 1 cm·s^−1^. Single-trial learning was calculated as the change in adaptation between the first and second channel trial of a triplet.

To identify changes in single-trial learning between triplets, repeated-measures ANOVAs were performed. All statistical tests were two-sided with significance set to *P* < 0.05. Data analysis was performed using MATLAB R2020a.

## COIN Model

### Generative model

Fig. S1 shows the graphical model for the generative model. At each time step *t* = 1,…, *T* there is a discrete latent variable (the context) *c_t_* ∈ {1,…, ∞} that evolves according to Markovian dynamics:

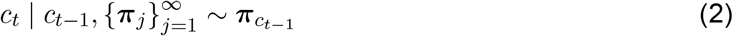

where 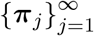 is the context transition probability matrix and *π_j_* is its *j*^th^ row containing the probabilities of transitioning from context *j* to all contexts (including itself). In principle, there are an infinite number of rows and columns in this matrix. However, in practice, generation and inference can both be accomplished using finite-sized matrices by placing an appropriate prior on the matrix (see below).

Each context *j* is associated with a continuous (scalar) latent variable 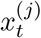 (the state, e.g. the strength of a force field or the magnitude of a visuomotor rotation) that evolves according to its own linear-Gaussian dynamics independently of all other states:

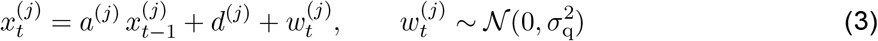

where *a*^(*j*)^ is the state retention factor, *d*^(*j*)^ is the state drift and 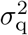 is the variance of the process noise (shared across contexts). Each state is assumed to have existed for long enough for its prior to be its stationary distribution: 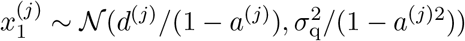.

At each time step, a continuous (scalar) observation *y_t_* (the state feedback) is emitted from the state associated with the current context:

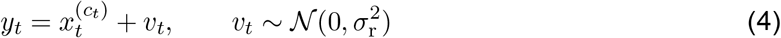

where 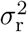 is the variance of the measurement noise (also shared across contexts).

In addition to the state feedback, a discrete observation (the sensory cue) *q_t_* ∈ {1,…, ∞} is also emitted. The distribution of sensory cues depends on the current context:

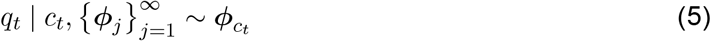

where 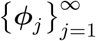 is the cue emission probability matrix (which, in principle, is also doubly infinite in size but can be treated as finite in practice) and *ϕ_j_* is its *j*^th^ row containing the probabilities of emitting all cues from context *j*.

In order to make this infinite-dimensional switching state-space model well-defined, we place hierarchical Dirichlet process ^31^ priors on the context transition and cue emission probability matrices. The context transition probability matrix is generated in two steps. First, a set of global transition probabilities 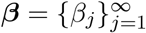 are generated via a recursive, stochastic ‘stick-breaking’ construction:

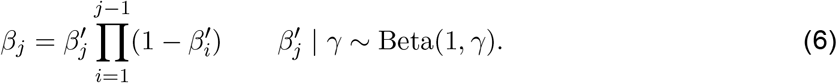

We denote the distribution of the global transition probabilities by *β* ~ GEM(*γ*). Note that 0 ≤ *β_j_* ≤ 1 and 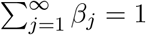, as required for a set of probabilities. The hyperparameter *γ* determines the effective number of contexts: large *γ* implies a large number of small-probability contexts, while small *γ* implies a smaller number of relatively large-probability contexts.

Each row of the context transition matrix is then generated using a ‘sticky’ variant ^32^ of the Dirichlet process:

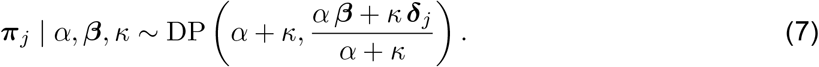

where *δ*_j_ is a one-hot vector with the *j*^th^ element set to 1 and all other elements set to 0. The concentration (inverse variance) parameter *α* + *κ* controls the variability of transition probabilities around the base (mean) distribution (*αβ* + *κδ_j_*)/(*α* + *κ*), with large *α* + *κ* reducing this variability. The self-transition bias (stickiness) hyperparameter *κ* increases the probability of a self-transition (*c_t_* = *c*_*t*−1_ = *j*) and expresses the fact that a context often persists for several time steps before switching, such as when an object is manipulated for an extended period of time or a surface is walked on. Note that the rows of the context transition matrix are dependent as their expected values (the base distributions of the corresponding Dirichlet processes) contain a shared term, the global transition distribution. This dependency captures the intuitive notion that contexts that are more common in general will be transitioned to more frequently from all contexts. In the limit of *α*, *κ* → ∞, the rows become identical except for the location of the self-transition bias term (i.e. 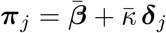, where 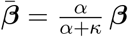 and 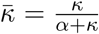), and the generative model implied by the Dirichlet process Kalman filter ^6^ is recovered as a special case.

The cue emission probability matrix 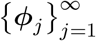 is generated using an analogous (non-sticky) hierarchical construction:

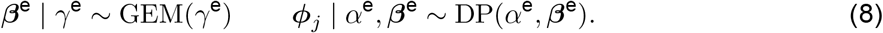

where *γ*^e^ determines the distribution of the global cue emission probabilities *β*^e^, and *α*^e^ determines the across-context variability of local cue emission probabilities around the global cue emission probabilities.

In order to allow full Bayesian inference over the model parameters, we also placed a prior on the parameters governing the state dynamics (***f***^(*j*)^ = [*a*^(*j*)^ *d*^(*j*)^]^⊤^) that was a bivariate normal distribution (truncated between 0 and 1 for *a*^(*j*)^):

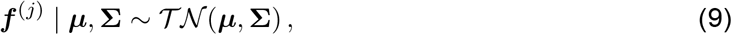

where ***μ*** = [*μ*_a_ *μ*_d_]^⊤^ and 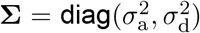 is a diagonal covariance matrix. Under the assumption that positive and negative drifts are equally probable, we set *μ*_d_ to zero.

#### The Chinese restaurant franchise

Here we present an alternative representation of the hierarchical Dirichlet process that provides a mechanism for generating sequences of contexts and sensory cues in the COIN model as well as a framework for inference. In addition, this representation provides intuitions for how the generative process works as trials are experienced.

In the Chinese restaurant franchise ^31^, there are an infinite number of restaurants each with an infinite number of tables. Each table serves only one dish from an infinite global menu shared by all the restaurants (hence the franchise). The same dish can be served on multiple tables in the same restaurant as well as in multiple restaurants. Each customer enters a restaurant and is seated at a table where a dish is served.

In the COIN model, customers correspond to trials and will arrive in the same temporal order as the trials. For both the context transitions and the cue emissions, the restaurant that the customer enters corresponds to the current context (i.e. if the current context is *j*, the customer enters restaurant *j*). For the context transitions, the dish served at the table at which the customer sits corresponds to the next context (i.e. if dish *c* is served, a transition to context *c* occurs). For cue emissions, the dish served at the table corresponds to the sensory cue emitted on that trial (i.e. if dish *q* is served, cue *q* is emitted). Note that separate Chinese restaurant franchises are used for context transitions and cue emissions.

Although there an infinite number of restaurants, tables and dishes in the franchise, to generate a finite amount of data, we only need to consider the finitely many occupied tables and the dishes served at those tables (i.e. the contexts and sensory cues already experienced), as well as one empty table in each occupied restaurant and one novel dish (allowing for a novel context or sensory cue).

##### Table assignment and dish selection

Upon entering a restaurant, each customer sits at an occupied table with probability proportional to the number of people already sitting at that table or a new table with probability proportional to *α*. This has the effect that tables with many customers attract even more customers, and since the table determines the next context (or sensory cue), this makes commonly experienced transitions (or cues) increasingly likely in the future.

The first customer to sit at a new table chooses the dish for that table. The customer chooses a previously-served dish with probability proportional to the number of other tables in the franchise serving that dish or a new dish with probability proportional to *γ*.

The hyperparameter *α* controls how the number of tables grows as a function of the number of customers. With small *α*, there will be few tables, and so the distribution of dishes served at those tables will depend strongly on the restaurant (i.e. context transitions and cue emissions will depend strongly on the current context). With large *α*, there will be many tables, leading to a more similar distribution of dishes across the restaurants (i.e. context transitions and cue emissions will be less dependent on the current context).

The hyperparameter *γ* controls how the number of dishes grows as a function of the number of tables. With small *γ*, most tables will have the same dish, and so the number of dishes (i.e. contexts and cues) will grow slowly over trials. With large *γ*, most tables will have different dishes, and so the number of dishes (i.e. contexts and cues) will grow rapidly over trials.

##### Loyal customers

For the context transitions, the process we have described so far has no self-transition bias. To include such a bias (as in the COIN model), the Chinese restaurant franchise can be extended to include loyal customers ^32^.

Each restaurant now has a specialty dish whose index is the same as that of the restaurant (e.g. dish *j* is the specialty dish of restaurant *j*). The specialty dish is available in all restaurants, but is more popular in the dish’s namesake restaurant. As we shall see, this leads to family loyalty to a restaurant. Consider contexts *c*_*t*−1_, *c_t_* and *c*_*t*+1_, which we shall refer to as the grandparent, parent and child, respectively. The parent enters restaurant *j*, determined by the grandparent *c*_*t*−1_ = *j*, and sits at table *τ* that serves dish *k_jτ_*. The dish the parent eats determines the restaurant that the child eats at, that is *k_jτ_* = *c_t_*. The increased popularity of the specialty dish means that children are more likely to eat at the same restaurant as their parent. Hence, multiple generations often eat at the same restaurant.

To simplify inference in the Chinese restaurant franchise with loyal customers, a distinction is made between a *considered* dish and a *served* dish. The first customer to sit at a table chooses a dish for the table without acknowledging the increased popularity of the specialty dish of the restaurant. However, with some probability, this considered dish is overridden (perhaps by a waiter’s suggestion) and the specialty dish is served instead. This process is described as follows:

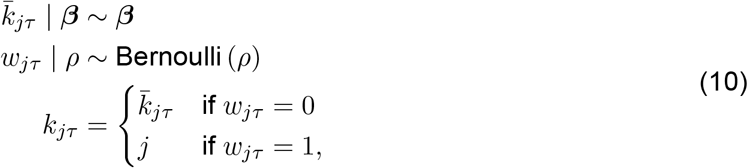

where 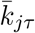 is the considered dish, *w_jτ_* is an override variable, *ρ* = *κ*/(*α* + *κ*) and *k_jτ_* is the served dish. When *κ* = 0, the served dish (which is always the same as the considered dish, as *w_jτ_* = 0 with probability one) is distributed as *k_jτ_* | *β* ~ *β*, where *β* represents the overall popularity or ratings of the dishes. When *κ* ≠ 0, the increased popularity of the specialty dish leads to the following modified dish ratings:

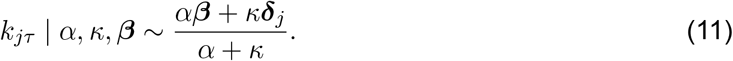

### Inference with particle learning

The goal of inference is to estimate a joint posterior distribution over the context, the state in each context and the parameters governing the state transitions, the context transitions and the sensory cue emissions at each point in time based on the state feedback and sensory cues observed so far. To perform posterior inference in an online (i.e. recursive) manner, we use a sequential Monte Carlo (simulation-based) method known as particle learning ^33,34^.

Particle learning extends standard particle filtering methods by incorporating the estimation of static parameters via a fully-adapted filter that utilizes conditional sufficient statistics for the posterior of the parameters. To sequentially compute a particle approximation to the joint posterior distribution of states, contexts and conditional sufficient statistics for the parameters, an essential state vector is constructed and is used together with a predictive distribution and propagation rule to build a resampling-sampling framework.

Central to particle learning is the essential state vector, *z_t_*, which contains samples and/or sufficient statistics of the states, context and parameters. Online state filtering and parameter learning is equivalent to sequential filtering of the essential state vector:

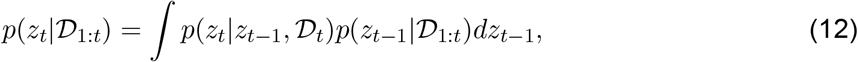

where 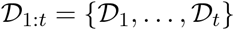 is the sequence of observations up to time *t*.

Particle methods use a system of *P* particles to form a discrete approximation to 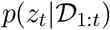 via

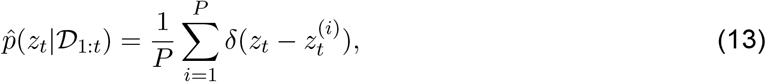

where *δ*(·) is a Dirac delta function. A recursive formula for obtaining 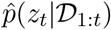 from 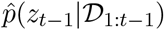 is suggested by the following decomposition of Eq. 12:

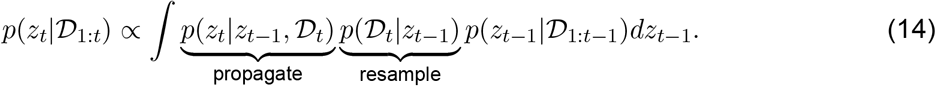

First, in a *resample* step, particles are sampled with replacement from a multinomial distribution with weights proportional to the predictive distribution 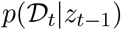. This produces a particle approximation to the smoothed distribution 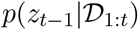 by replicating/discarding particles based on how well they predicted the observations at time *t* Then, in a *propagate* step, the resampled particles are propagated via the evolution equation 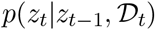. A final *sample* step can also be performed in which the parameters, *θ*, are sampled from their posterior distribution conditioned on the propagated essential state vectors. Although this last step is optional, without it the diversity of parameters would reduce with each resampling step until all particles shared the same parameters, a problem known as degeneracy. A single time step of particle learning is summarized in Algorithm 1.

In the COIN model, the essential state vector 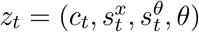 contains the context, the sufficient statistics (mean and variance) for the states 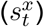, the sufficient statistics for the parameters 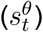 and the parameters (*θ*). Following the direct assignment algorithm of Teh et al. ^31^, we do not instantiate the context transition and cue emission probability matrices. Instead, we instantiate (sample) the global transition and emission distributions and integrate out the context transition and cue emission probability matrices (i.e. compute their expected values) conditioned on these instantiated global distributions. Hence, in the COIN model, 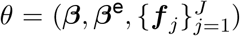. The propagate step in Algorithm 1 can be decomposed into three separate steps:

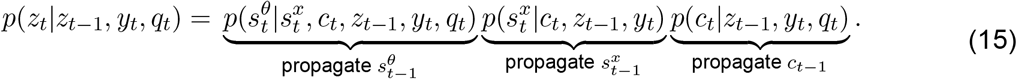

**Algorithm 1:**
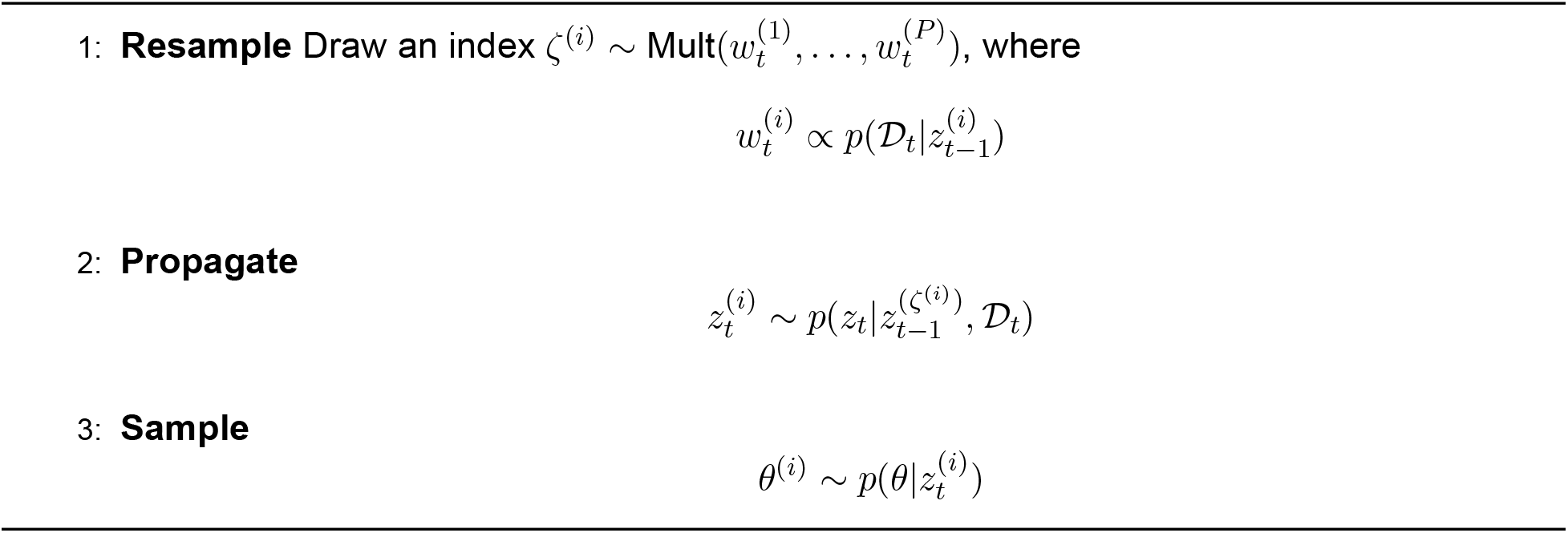
The general particle learning algorithm.

First, the context is propagated; then the sufficient statistics for the states are propagated conditioned on the context and the state feedback; and then the sufficient statistics for the parameters are propagated conditioned on the context, the sufficient statistics for the states, the state feedback and the sensory cue.

We now describe the resample, propagate and sample steps for the COIN model in detail.

#### Resample

Given the particle approximation 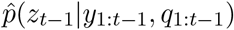, the updated smoothed approximation 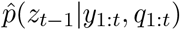 is obtained by resampling particles with weights *w_t_* proportional to the predictive distribution:

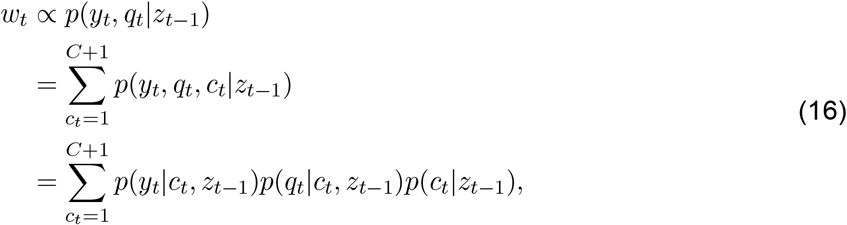

where *C* is the number of contexts instantiated up to *t* − 1. Note that the sum over contexts is to *C* + 1 to include the possibility that the latest observations were generated by a new context, the (*C* + 1)^th^ context. The context transition term of the predictive distribution is given by

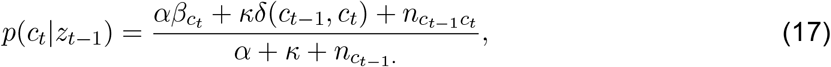

where 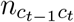 denotes the number of transitions from context *c*_*t*−1_ to context *c_t_* up to time *t*−1. Dots represents marginal counts. For example, 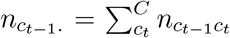 is the number of transitions out of *c*_*t*−1_) depends on the global transition probability 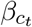. Thus, when the global transition distribution is updated (see below), the inferred transition probabilities from all contexts are also updated. An important consequence of this is that transition probabilities learned in one context will generalize to all contexts.

The cue emission term has a similar formulation to the context transition term:

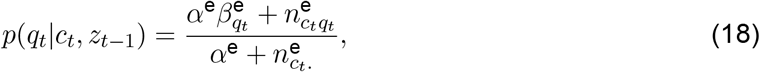

where 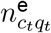 denotes the number of emissions of cue *q_t_* in context *c_t_* up to time *t* − 1, 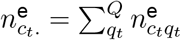 is the number of cues emitted in context *c_t_* up to time *t* − 1 and *Q* is the number of cues emitted up to time *t* − 1.

The state feedback term depends on the estimate of the state in each context and is given by

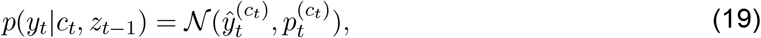

where 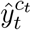 and 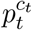 are the mean and variance of the predicted state feedback for context *c_t_* given by the time update equations of the Kalman filter (Algorithm 2).

**Algorithm 2:**
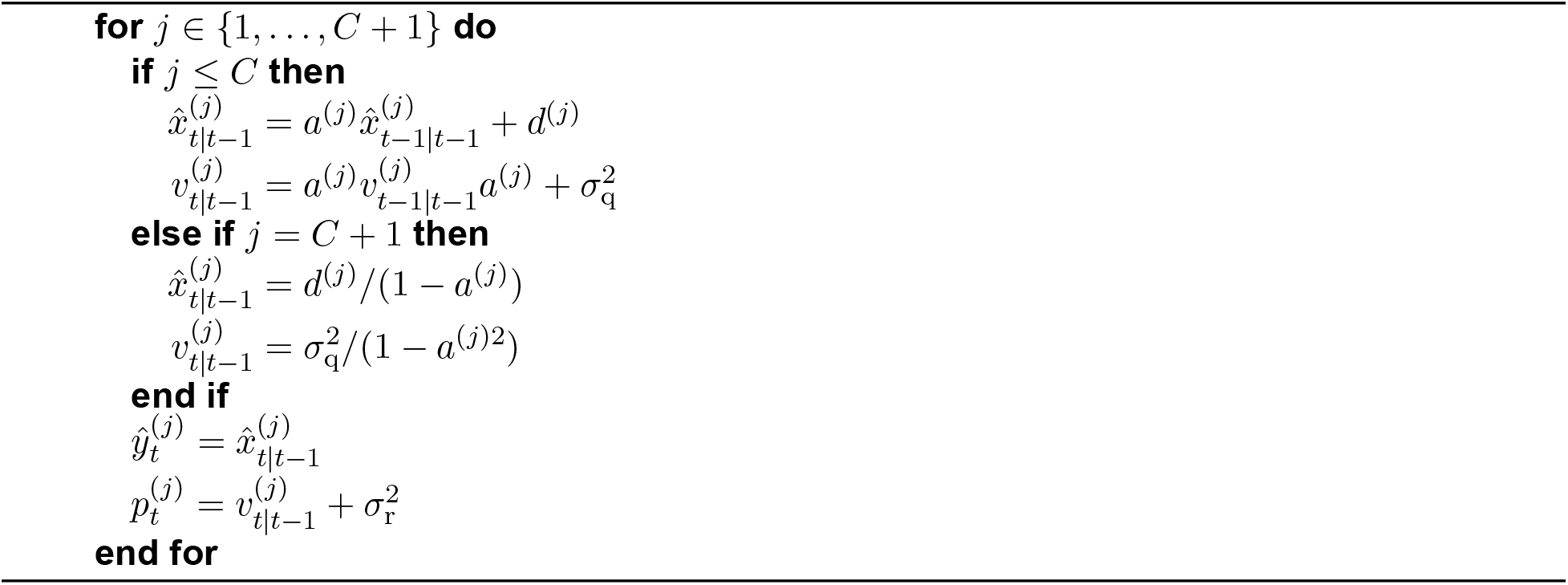
State and state feedback prediction for instantiated contexts and a novel context. For a novel context, the mean and variance of the predicted state are the moments of the stationary distribution conditioned on state retention and drift parameters sampled from the prior.

#### Propagate the context and the state

The particles of the smoothed approximation 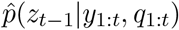 are propagated via the three steps outlined in Eq. 15.

##### Propagate the context

The context is propagated by sampling *c_t_* ∈ {1,…, *C* + 1} from the filtering distribution:

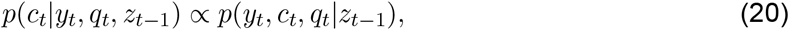

where *p*(*y_t_*, *c_t_*, *q_t_*|*z*_*t*−1_) is given in Eq. 16. These filtered probabilities are the so-called ‘responsibilities’.

If *c_t_* = *C* + 1 (i.e. the context is new), *C* is incremented and *β* is transformed by sampling *b* ~ Beta(1, *γ*) and assigning 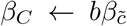 and 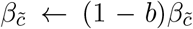, where 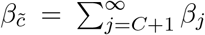. Similarly, if *q_t_* = *Q* +1 (i.e. the sensory cue is new), *Q* is incremented and *β*^e^ is transformed by sampling *b*^e^ ~ Beta(1, *γ*^e^) and assigning 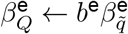 and 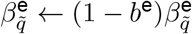, where 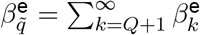.

##### Propagate the sufficient statistics for the states

Conditioned on the sampled context variable, the sufficient statistics (mean and variance) for the state of each instantiated context are propagated via the measurement update equations of the Kalman filter (Algorithm 3).

**Algorithm 3:**
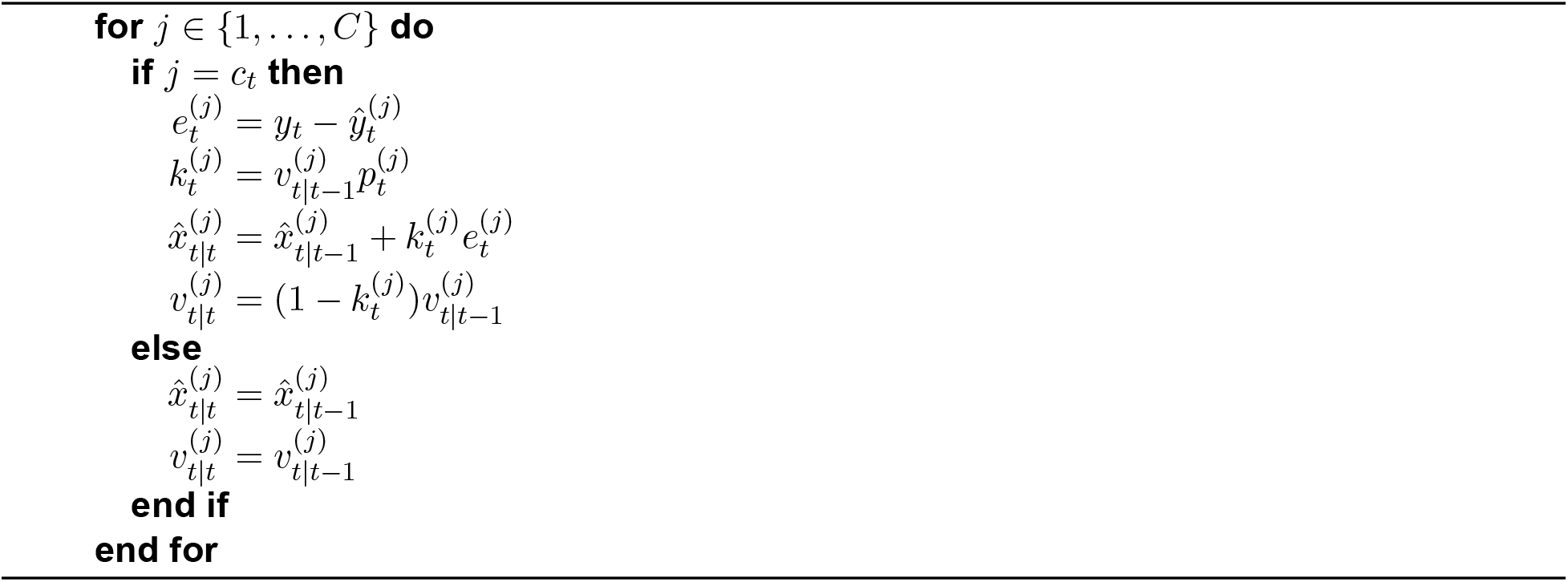
State filtering. The difference between the actual and predicted state feedback (i.e. the prediction error) is used to update the predicted state for the active context (i.e. the context responsible for generating the state feedback). The prediction error is scaled by the Kalman gain, which is close to 0 when 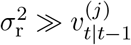 and close to 1 when 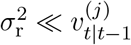.

#### Propagate the sufficient statistics for the parameters

##### Propagate the sufficient statistics for the state retention and drift parameters

For each *j* ∈ {1,…, *C*}, a pair of states 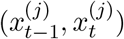 are sampled from

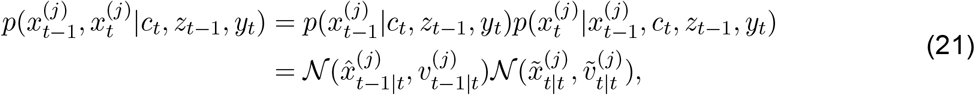

where 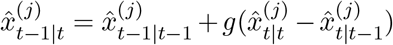, 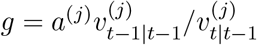, 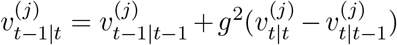, 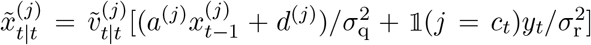 and 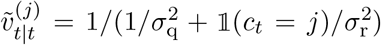. Here the indicator function 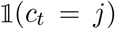 is equal to 1 if the sampled value of *c_t_* = *j* and 0 otherwise. The sufficient statistics for the state retention and drift parameters are then propagated as follows:

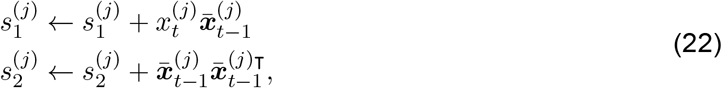

where 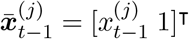.

##### Propagate the sufficient statistics for the context transition and cue emission parameters

The context transition counts and the cue emission counts are propagated by incrementing 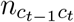 and 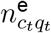, respectively.

#### Sample the parameters

##### Sample the state retention and drift parameters

For each *j* ∈ {1,…, *C*}, the hyperparameters of the posterior distribution of the state retention and drift parameters are computed:

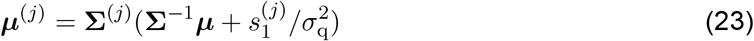

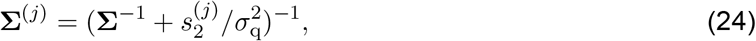

and a new set of state retention and drift parameters are sampled from

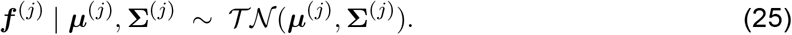

##### Sample the global emission distribution parameters

To sample *β*^e^, a Chinese restaurant process is first simulated to sample each 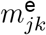 (the number of tables in restaurant *j* serving dish *k*). For each *j* ∈ {1,…, *C*} and *k* ∈ {1,…, *Q*}, 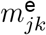 and *n* are initialized to 0. Then, for 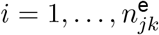 (i.e. for each customer in restaurant *j* eating dish *k*), a sample is drawn from

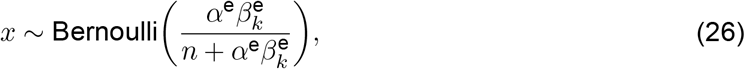

*n* is incremented, and if 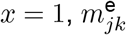 is incremented.

Conditioned on 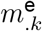 (the total number of tables serving dish *k*), the global emission distribution is sampled from

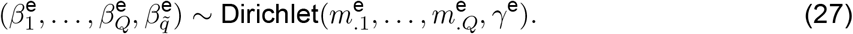

##### Sample the global transition distribution parameters

To sample *β*, a Chinese restaurant process is first simulated to sample each *m_ij_* (the number of tables in restaurant *i serving* dish *j*). For each (*i, j*) ∈ {1,…, *C*}^2^, *m_ij_* and *n* are initialized to 0. Then, for *i* = 1,…, *n_ij_* (i.e. for each customer in restaurant *i* eating dish *j*), a sample is drawn from

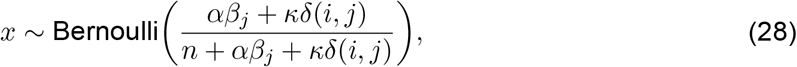

*n* is incremented, and if *x* = 1, *m_ij_* is incremented.

To obtain 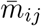 (the number of tables in restaurant *i considering* dish *j*), *w_i._* (the number of override variables in restaurant *i*) is sampled from

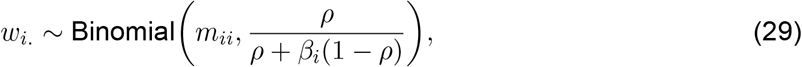

and subtracted from *m_ii_*:

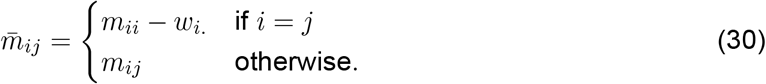

Conditioned on 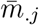 (the total number of tables considering dish *j*), the global transition distribution is sampled from

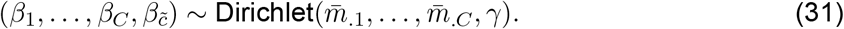

#### COIN model implementation

We applied the above inference algorithm to a sequence of noisy state feedback observations and sensory cues (if present, numbered according to the order they were presented in the experiment). On each trial, the state feedback was assigned a value of 0 (null-field trials), +1 (*P*^+^ force-field trials) or −1 (*P*^−^ force-field trials) plus i.i.d. zero-mean Gaussian observation noise with variance 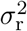. Because both motor noise and sensory noise influence observed movement kinematics (state feedback), we set 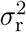 to 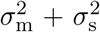 under the assumption that motor and sensory noise are i.i.d. Gaussian variables (with variances 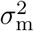 and 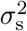, respectively) that sum to produce the final observation noise. We assumed that sensory noise is typically no more than around one tenth of the perturbation magnitude, and hence we set *σ*_s_ to 0.03 to reduce the number of free parameters in the model.

The algorithm was initialized with *C* = 0, *Q* = 0, *β*_1_ = 1, 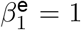 and the sufficient statistics for the parameters set to 0.

On channel trials (*P*^c^), the state (e.g. the magnitude of a force field or visuomotor rotation) is not observed. Therefore, we omitted state feedback on these trials. This was achieved by modifying the general inference algorithm of the model in the following ways:

1. The state feedback likelihood term (Eq. 19) did not contribute to the weights used to resample particles (Eq. 16) or the probabilities used to propagate the context (Eq. 20).
2. The measurement update steps were omitted when updating the state estimate (Algorithm 3); that is for all *j* ∈ {1,…, *C*}, 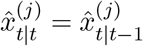 and 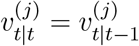. Hence, on channel trials, each state estimate is updated based only on the inferred dynamics (retention and drift) ascribed to that context and there is no error-based learning.
3. To propagate the sufficient statistics for the retention and drift parameters, a pair of states 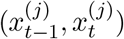 are sampled from a distribution that is equivalent to Eq. 21 but that does not condition on *y_t_* (and hence does not condition on *c_t_* either):

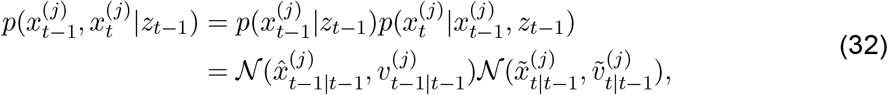

where 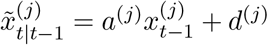 and 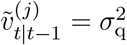.

To reduce the number of free parameters in the model, we set *γ* = *γ*^e^ = 0.1. The only exception to this was model validation in which we set *γ* = *γ*^e^ = 0.3 to generate distributions of contexts and cues that are typical of motor learning experiments.

To place a finite limit on the maximum number of contexts that the COIN model can instantiate, we truncated the stick-breaking process of the GEM. We chose a truncation level of 10 as this number was greater than the true number of contexts in any of the experiments we modeled (typically 2-3). As an exception, in the ntext setting of model validation, the truncation level was set to 1 (see Validation of the COIN model). Note that when the truncation level is set to 1, the nonparametric switching state-space model reduces to a ntext (i.e. non-switching) state-space model. Moreover, if the parameters of the state transition dynamics are also known (i.e. not learned online), the Kalman filter is recovered as a special case of the COIN model.

Adaptation (*a_t_*) on trial *t* was modeled as the net predicted state (*u_t_*) plus Gaussian motor noise:

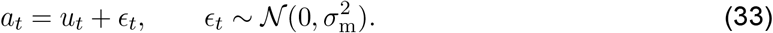

The net predicted state was obtained by summing the predicted states for each instantiated context and a novel context (weighted by their predicted probabilities) and averaging across particles. The predicted probabilities are given by

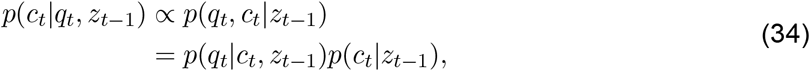

where *p*(*q_t_*|*c_t_*, *z*_*t*−1_) and *p*(*c_t_*|*z*_*t*−1_) are defined in Eqs. 17 and 18.

#### Modeling visuomotor rotation experiments in the COIN model

In visuomotor rotation experiments, the cursor moves in a different direction to the hand (which is occluded from vision). This introduces a discrepancy between the location of the hand as perceived by vision and proprioception. To model this discrepancy, we included a bias parameter in the state feedback (equation 4):

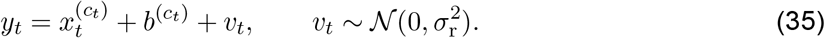

To support Bayesian inference, we placed a normal distribution prior over this parameter:

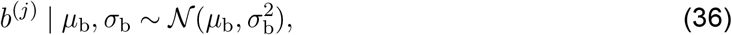

where *μ*_b_ was set to zero as positive and negative biases are equally likely, and *σ*_b_ was set to 70^−1^ by hand to match the empirical data in Fig. S16a.

The inference algorithm was correspondingly modified and extended in the following ways:

1. The predicted state feedback (Algorithm 2) became 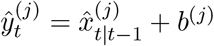.
2. The sufficient statistics for the bias parameter are propagated as follows:

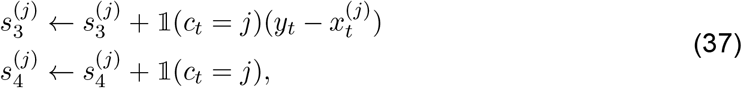

where 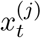 is sampled from 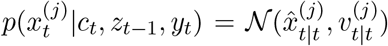. Note that this step is omitted on channel trials as there is no state feedback.
3. The hyperparameters of the posterior distribution of the bias parameter are

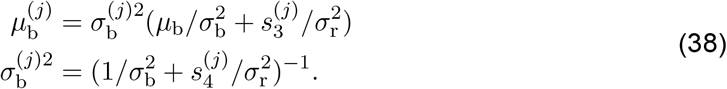
4. A new bias parameter is sampled from the updated posterior at the end of each trial:

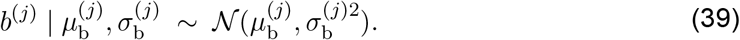

#### Simulating existing data sets

The paradigms in Fig. 4 and Fig. S14 were simulated using the 40 sets of parameters fit to individual participants’ data in both experiments. One hundred simulations (each conditioned on a different noisy state feedback sequence) were performed for each parameter set. The results shown are based on the average of all of these simulations.

The paradigms in Fig. S16 and Fig. S15 were variations of the standard spontaneous recovery paradigm. Therefore, we simulated these paradigms using the parameters fit to the average spontaneous and evoked recovery data sets (Table S1). One hundred simulations (each conditioned on a different noisy state feedback sequence) were performed. The results shown are based on the average of these simulations.

##### Savings paradigm

We used the paradigm described in Coltman et al. ^13^ to simulate savings in the COIN model (Fig. 4a). Participants completed two force-field learning sessions that were separated by a 5-minute break. Each session consisted of a pre-exposure phase (60 trials) with a null field (*P*^0^). This was followed by an exposure phase (125 trials) with a velocity-dependent curl field (*P*^+^). After the exposure phase, participants performed a counter-exposure phase (15 trials) with the opposite curl field (*P*^−^). This was followed by a series of 50 channel trials (*P*^c^). In addition, channel trials were randomly interspersed throughout the exposure phase (approximately 1 in every 10 trials).

##### Anterograde interference paradigm

We used the paradigm described in Sing and Smith ^14^ to simulate anterograde interference in the COIN model (Fig. 4b). The paradigm consisted of a pre-exposure phase (160 trials) with a null field (*P*^0^). This was followed by an exposure phase of variable length (13, 41, 112, 230, or 369 trials) with a velocity-dependent curl field (*P*^+^). After the exposure phase, participants performed a counter-exposure phase (115 trials) with the opposite curl field (*P*^−^). Channel trials were randomly interspersed throughout the exposure and counter-exposure phases (approximately 1 in every 7 trials).

##### Environmental consistency paradigms

We used the paradigm described in Experiment 1 in Herzfeld et al. ^9^ to simulate the effect of environmental consistency on single-trial learning in the COIN model (Fig. 4c). The paradigm consisted of a pre-training phase (156 trials) with a null field (*P*^0^) interspersed with triplets (1 in every 13 trials). This was followed by a training phase composed of 25 blocks (45 trials each). Each block of the training phase consisted of a sequence of 30 perturbation trials, 2 channel trials, 10 washout trials, and 1 triplet. During the perturbation trials, either *P*^+^ or *P*^−^ was presented. Across trials, the perturbation either switched with probability *p*(switch) or remained the same with probability *p*(stay) = 1 − *p*(switch). Three groups performed the experiment with p(stay) set to 0.9, 0.5 and for the groups who experienced the slowly, medium and rapidly switching environment, respectively.

As an additional demonstration of the effect of environmental consistency on single-trial learning in the COIN model (Fig. S14), we simulated the P1N1, P1, P7, and P20 environments of the force-field adaptation task described in Gonzalez Castro et al. ^12^. The paradigm consisted of a pre-exposure phase (200 trials) with a null field (*P*^0^). In the anti-consistent environment (P1N1), participants experienced 50 cycles each with a single *P*^+^ trial, followed by a single *P*^−^ trial, followed by 11–13 *P*^0^ trials. In the inconsistent environment (P1), participants experienced 45 cycles with a single *P*^+^ trial, followed by 10-12 *P*^0^ trials. In the moderately consistent environment (P7), participants experienced 27 cycles with seven *P*^+^ trials, followed by 15–18 *P*^0^ trials. In the highly consistent environment (P20), participants experienced 27 cycles with 20 *P*^+^ trials, followed by 28–32 *P*^0^ trials. To assess single-trial learning during exposure to the environments, channel trials were randomly interspersed before and after the first *P*^+^ trial in a subset of the force-field cycles.

##### Working memory and evoked recovery

We investigated the effect of a working memory task on contextual inference in the COIN model (Fig. S15) by simulating a force-field adaptation task (Experiment 1) in Keisler and Shadmehr ^19^. The paradigm consisted of a pre-exposure phase (192 trials) with a null field (*P*^0^). This was followed by an exposure phase (384 trials) with a velocity-dependent curl field (*P*^+^). After the exposure phase, participants performed a counter-exposure phase (20 trials) with the opposite curl field (*P*^−^). Participants then completed either a memory task (memory group) or a non-memory task (non-memory group). This was followed by a series of 192 channel trials (*P*^c^). Channel trials were randomly interspersed throughout the pre-exposure and exposure phases (1 in every 8 trials).

In the memory task, participants were shown 12 word pairs (e.g., “COMFORT-ATOM”, “LEGEND-BLANK”). Immediately after viewing the words, participants were then shown one word from each pair and instructed to say the corresponding word aloud. In the non-memory task, participants were shown strings of letters (e.g., “kdinedlr”) and were instructed to say aloud the number of vowels in each string.

We hypothesized that context probabilities, which are updated recursively in the COIN model, are maintained and updated in working memory. The effect of the working memory task is to erase these estimated probabilities from memory so that participants instead infer the context based on the stationary distribution, which represents the expected frequency of each context. Therefore, on the first trial of the channel phase (i.e. directly after the working memory task), we set the predicted probabilities to their values under the stationary distribution (calculated from the expected value of the context transition matrix under the Dirichlet posterior). For the non-memory task, the COIN model was simulated as for a standard spontaneous recovery paradigm.

##### Explicit and implicit visuomotor learning

We investigated explicit and implicit learning in the COIN model (Fig. S16) by simulating a visuomotor rotation task (Report condition of Experiment 2) described in McDougle et al. ^21^. The paradigm consisted of a pre-exposure phase (100 trials) in which cursor feedback was veridical (*P*^0^). This was followed by an exposure phase (200 trials) in which the cursor was rotated by 45° in the clockwise direction (*P*^+^, note we use this to represent a positive rotation in this visuomotor paradigm). After the exposure phase, participants performed a counter-exposure phase (20 trials) in which the cursor was rotated by 45° in the counterclockwise direction (*P*^−^). This was followed by a series of 100 visual error clamp trials (*P*^c^) in which the cursor moved straight to the target regardless of the participant’s hand trajectory. During the pre-exposure, exposure and counter-exposure phases, the target was flanked by a 360° ring of numbered visual landmarks spaced 5.625° apart. Starting at trial 91 of the pre-exposure phase, participants were instructed to report verbally before each reach the landmark that they planned to push the manipulandum toward to make the cursor hit the target. These reported aiming directions were interpreted as the explicit component of learning. Implicit learning was quantified by subtracting the explicit component from the actual movement direction on each trial. After the end of the counter-exposure phase, participants were told to stop using any aiming strategy that they had developed and reach directly for the target during the remaining visual error clamp phase. A control group performed the identical paradigm but without any reporting of aim direction.

To simulate learning in a visuomotor rotation experiment in the COIN model, we included an additional parameter to reflect measurement bias (the difference between hand location perceived by proprioception and vision), which was inferred online (see Modeling visuomotor rotation experiments in the COIN model).

#### Deterministic state-space models

We also fit a class of *n*-rate deterministic state-space models to the data in the spontaneous/evoked recovery experiment. These models frame motor adaptation as sequential estimation of a task perturbation (e.g., the magnitude of a force field) using *n* separate adaptive states, each of which has its own own retention factor and learning rate. For the two-state (dual-rate) model *n* = 2, and for the three-state model *n* = 3. The individual states can be arranged into a state vector:

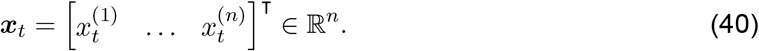

The motor output on trial *t* is the sum of the elements in the state vector:

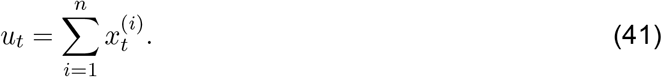

The error on trial *t* is the difference between the motor output and the task perturbation:

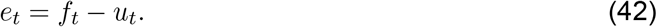

The task perturbation, *f*, was zero for null-field trials, +1 for *P*^+^ field trials and 1 for *P*^−^ field trials. The state vector is updated across trials:

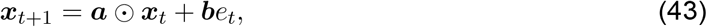

where ***a*** = [*a*^(1)^…*a*^(*n*)^]^⊤^ ∈ ℝ^*n*^ is a retention vector that governs trial-by-trial decay, ***b*** = [*b*^(1)^…*b*^(*n*)^]^⊤^ ∈ ℝ^n^ is a learning-rate vector that governs error-dependent adaptation and ⊙ denotes element-wise multiplication. For a task perturbation of +1, equation 43 can be rewritten as

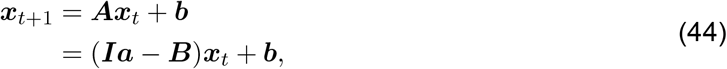

where ***I*** ∈ ℝ^*n*×*n*^ is the identity matrix and each column of ***B*** ∈ ℝ^*n*×*n*^ is equal to ***b***. We used this reparameterization to ensure that fitted parameters lead to stable learning as assessed through the eigenvalues of ***A*** (see below).

#### Model fitting

In both experiments, we fit the COIN model to the data of individual participants by fitting the set of parameters *ϑ* so as to maximize the data log likelihood. In the spontaneous/evoked recovery experiment, *ϑ* = {*σ*_q_, *μ*_a_, *σ*_a_, *σ*_d_, *α, ρ, σ*_m_}, and in the memory updating experiment, which included sensory cues, an additional parameter was fit so that *ϑ* = {*σ*_q_, *μ*_a_, *σ*_a_, *σ*_d_, *α, ρ, σ*_m_, *α*^e^}. The likelihood is given by

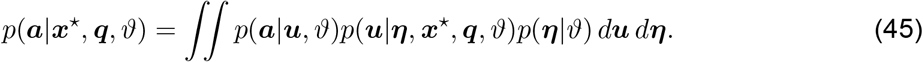

Here 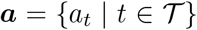 is the noisy motor output (adaptation) of the participant on the subset 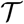 of trials that were channel trials, 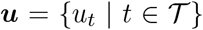 is the noiseless motor output of the model on the same subset of trials, 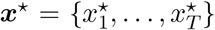 and ***q*** = {*q*_1_,…, *q_T_*} are, respectively, the perturbations and sensory cues (if applicable) presented to the participant and ***η*** = {*η*_1_,…, *η_T_*} is the observation noise. The actual observation noise that the participant perceived is unknown to us and so is marginalized out (equation 45). We approximated the likelihood using Monte Carlo integration:

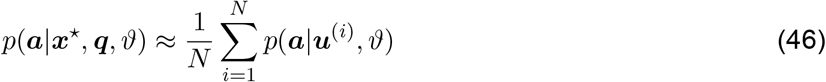

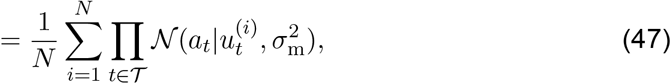

where each *u*^(*i*)^ ~ *p*(***u***|***η***^(*i*)^, ***x***^⋆^, ***q***, *ϑ*) was obtained by running the COIN model inference algorithm conditioned on an observation noise sequence ***η***^(*i*)^ sampled from 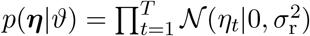. Note that this objective is stochastic because we sample observation noise. Consequently, to fit the COIN model, we used Bayesian adaptive direct search (BADS) ^35^, a Bayesian optimization algorithm that alternates between a series of fast, local Bayesian optimization steps and a systematic, slower exploration of a mesh grid. Optimization was performed from 30 random initial parameter settings with *P* = 100 particles and *N* = 100 observation noise sequences. Once each optimization was complete, we re-calculated the log likelihood using *P* = 1000 particles and *N* = 1000 observation noise sequences to obtain a lower-variance estimate of the log likelihood. This estimate was used to choose the best fit out of 30 for each participant.

To fit the COIN model to group average data (spontaneous/evoked recovery experiment), we defined the likelihood as

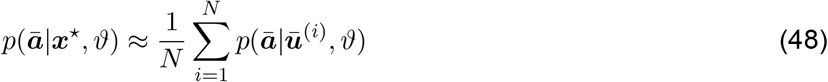

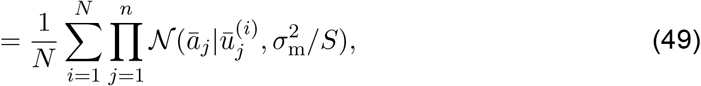

where 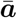 is the average adaptation data across participants, 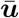 is the average noiseless motor output across participants and *S* is the number of participants in the group. Note that each element of 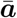 and 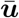 is indexed by the channel trial number (1,…, *n*) as opposed to the specific trial on which the channel trial occurred, as the latter was randomized across participants. The noiseless motor output for each participant was obtained, as above, by running the COIN model inference algorithm for each participant conditioned on an observation noise sequence sampled from *p*(***η***|*ϑ*). To fit the model to the average spontaneous recovery group data and the average evoked recovery group data using the same set of parameters, we optimized the sum of the log likelihoods of these groups.

To fit the deterministic state-space models, we minimized the mean squared error between the model data (equation 41) and the adaptation data measured on channel trials. Under the assumption that the adaptation data is the model data plus i.i.d. Gaussian noise, this is equivalent to maximizing the likelihood 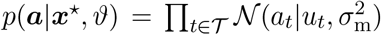, where 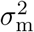 is the mean squared error. To ensure stable solutions, we constrained the eigenvalues of the matrix ***A*** in equation 44 to be between 0 and 1. Optimization was performed from 30 random initial parameter settings using both MATLAB’s fmincon and BADS. We report the best solution found by either optimizer.

#### Model comparison

To perform model comparison for individual participants, we calculated the Bayesian information criterion:

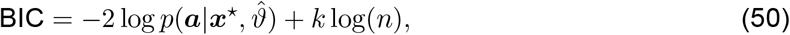

where 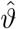 is the maximum likelihood estimate of the parameters, *k* is the number of parameters and *n* is the number of data points (channel trials). The first term in the BIC penalizes underfitting, whereas the second term penalizes model complexity, as measured by the number of free parameters in the model. Taking the difference in BIC values for two competing models approximates twice the log of the Bayes factor^36^. A BIC difference of greater than 4.6 (a Bayes factor of greater than 10) is considered to provide strong evidence in favor of the model with the lower BIC value ^37^. Note that the BIC penalizes model complexity more heavily than the Akaike information criterion (AIC) and corrected AIC (AICc), and hence, relative to AIC and AICc, BIC handicaps the COIN model as it has more parameters than the dual-rate model.

To perform model comparison at the group level, we calculated the group-level BIC, which is the sum of BICs over individuals ^38^.

#### Inferring internal representations of the COIN model fit to adaptation data

To examine the internal representations of the COIN model fit to adaptation data, we inferred the sequence of beliefs about the context, states and parameters, as encapsulated in the essential state vector *z*_1:*T*_. For each participant, this inference was conditioned on their observed adaptation data *a*_1:*T*_, their maximum likelihood COIN model parameters *ϑ* and the sequences of perturbations 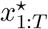 and sensory cues *q*_1:*T*_ presented to them. Thus we inferred the posterior distribution 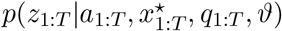. This inference can be performed recursively in time using particle filtering methods, in which beliefs are first sampled from the prior (by running the COIN model, which does not condition on the adaptation data) and then weighted by the likelihood (the probability of adaptation data given the sampled beliefs). Note that this involves a two-level hierarchy of particle methods: at the bottom level of the hierarchy particle learning is used to simulate inference from the perspective of the participant conditioned on state feedback and sensory cues, while at the top level of the hierarchy particle filtering is used to simulate inference from the perspective of the experimenter conditioned on the participant’s adaptation data.

Our assumptions about how adaptation data are generated are as follows. The beliefs of a participant evolve as a hidden Markov process with initial distribution *p*(*z*_1_|*ϑ*), conditional transition distribution *p*(*z_t_*|*z*_*t*−1_, *y_t_*, *q_t_*, *ϑ*) and observation distribution 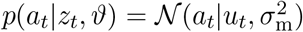. The conditional transition distribution denotes one update of the COIN model from trial *t* − 1 to time *t* conditioned on the state feedback and sensory cue observations at trial *t*.

A recursive formula for obtaining 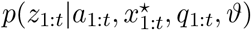 from 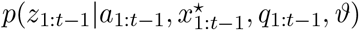 is

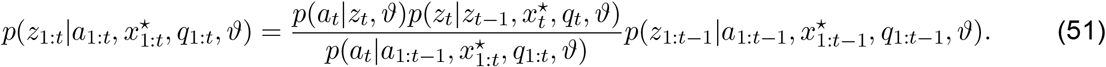

The marginal transition distribution in the numerator is obtained by integrating out the state feedback, which is unknown to us, from the conditional transition distribution:

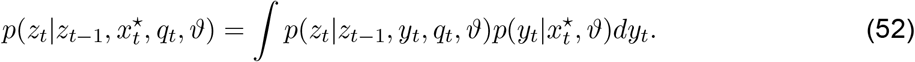

This integral can be approximated using Monte Carlo integration:

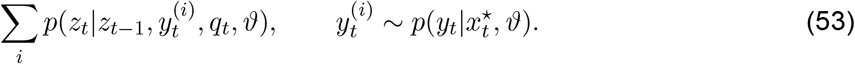

The recursive update in equation 51 is analytically intractable, and hence we use particle filtering methods. The importance distribution from which we draw samples of *z*_1:*T*_ has the following recursive form:

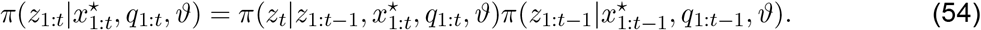

Note that this importance distribution does not condition on the adaptation data and so is termed the prior importance distribution. We define the importance function 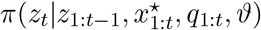 to be the marginal transition distribution 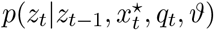, as this is straightforward to sample from. To correct for the use of an importance distribution, samples of *z*_1:*t*_ drawn from 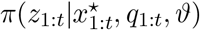 are assigned importance weights given by

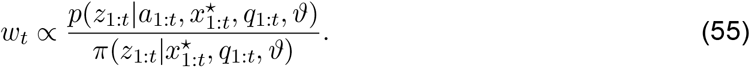

This yields a set of *N* weighted particles that approximates the target posterior:

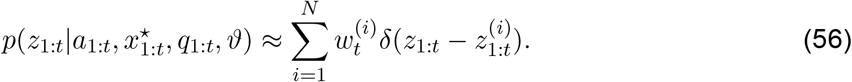

The recursive nature of equations 51 and 54 allows the weights in turn to be computed in a recursive manner:

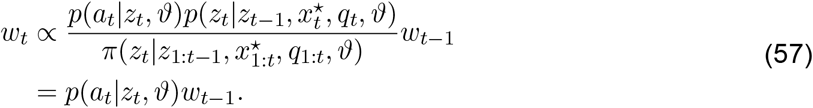

This weight update rule demonstrates that samples that place higher probability on the adaptation observations are assigned greater weight. Note that if trial *t* is not a channel trial, there is no adaptation observation for this trial and so *w_t_* ∝ *w*_*t*−1_. The computation of the importance weights involves multiplying incremental weights. Over many trials, this results in only a few particles having significant weight, a problem known as degeneracy. To avoid degeneracy, whenever the effective sample size (*ESS*) falls below a predefined threshold (half the total number of particles), we resample particles with probabilities proportional to their weights. The *ESS* is a standard metric based on the normalised weights and is typically defined as

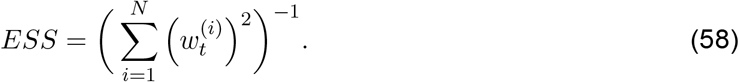

When all particles have equal weight, *ESS* = *N*, and when only one particle has nonzero weight, *ESS* = 1. The particle filtering algorithm is summarized in Algorithm 4 for a single trial.

The estimate of the posterior expectation of *z*_1:*T*_ is

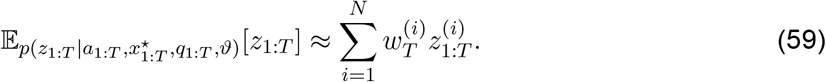

This method was used to generate the model data plotted in Fig. 2c,e and Fig. 3 and Figs. S8 and S11 to S12.

#### COIN model analysis

Particle methods present a challenge when it comes to COIN model analysis as contexts sampled by different particles cannot be directly mapped onto one another. This is because each particle samples its own context sequence, and this sequence may be different from the sequences sampled by other particles and/or the ground-truth context sequence. In addition, different particles can instantiate a different number of contexts, adding further complexity to the problem of finding a correspondence between contexts across particles. Hence, contexts that have the same label (assigned based on the order in which they were instantiated) in different particles do not necessarily correspond to the same inferred or ground-truth context. This means that contexts across particles cannot be simply equated based on the order of their instantiation. To address this issue we took two approaches. In some instances (Figs. 3 and 4 and Figs. S11 and S14), we focused our analysis on one specific context (*c*\*) that could be easily defined in all particles (e.g. the context with the largest predicted probability or responsibility). In all other instances, we fit models of reduced complexity to the particle system as a whole to extract simpler representations of the internal workings of the model. We now describe this process in detail.

**Algorithm 4:**
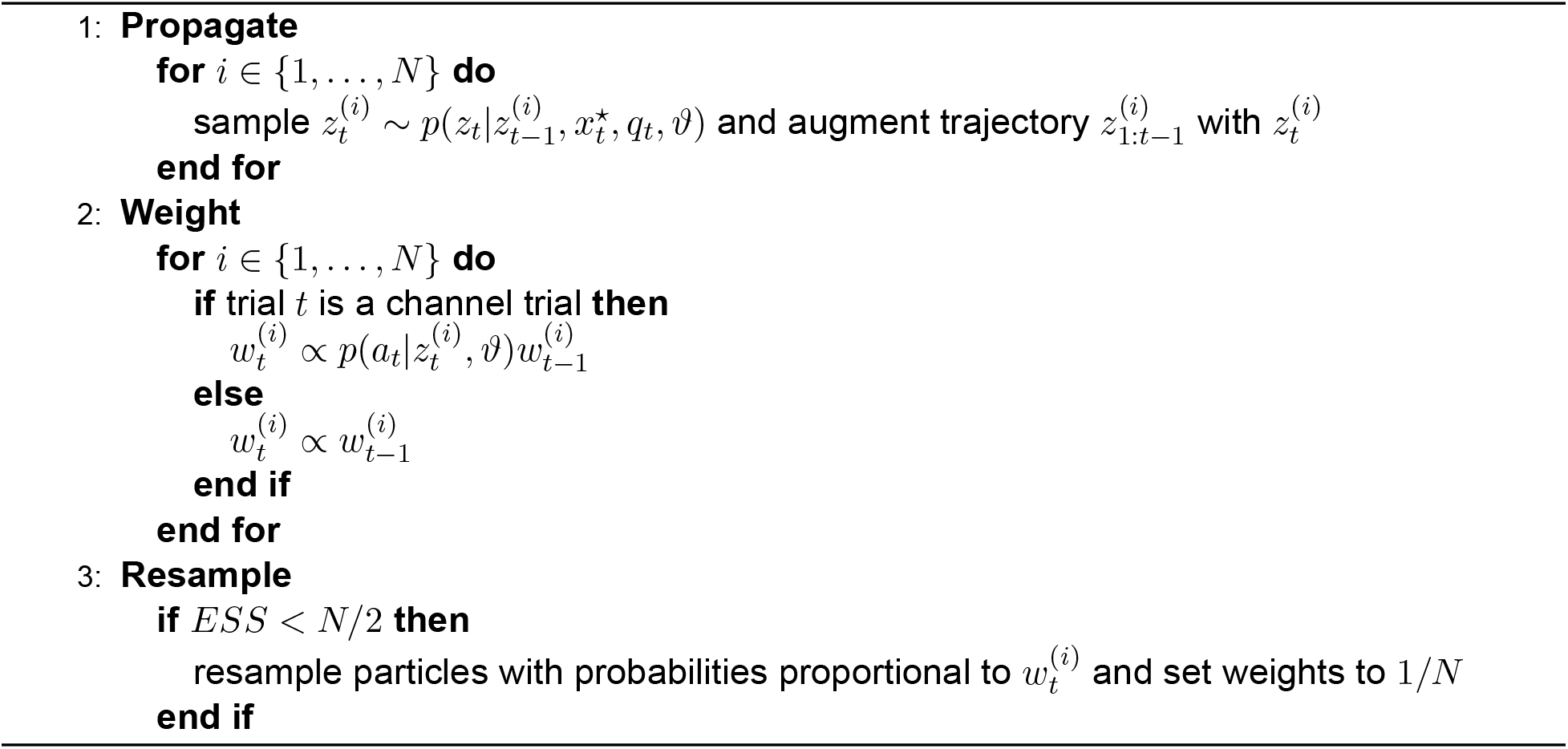
Particle filtering to infer a participant’s internal beliefs conditioned on their adaptation data.

To extract representations of the continuous variables (states and state parameters in e.g. Fig. 1 and Fig. S2), on each trial we first constructed a target density by marginalizing over all contexts, particles and observation noise sequences. For example, for the predicted state distribution, the target density is a mixture of Gaussians:

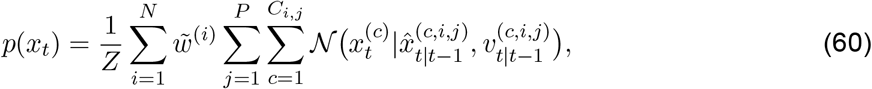

where *C*_i,j_, denotes the number of contexts instantiated by particle *j* conditioned on observation noise sequence *i*, *P* is the number of particles, *N* is the number of observation noise sequences, and *Z* is a normalization constant. The weight 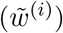 assigned to observation noise sequence *i* is 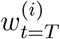 (Eq. 57) for simulations conditioned on adaptation data (see Inferring internal representations of the COIN model fit to adaptation data) or 1*/N* for simulations not conditioned on adaptation data. We performed component reduction on this mixture by approximating it with another mixture of Gaussians that had far fewer components. The key idea is that each component in the reduced-component mixture represents the distribution of the state in a specific context. The reduced-component mixture was fit by optimizing the parameters (mean and variance) of each component using expectation-maximization. The number of components (*K*) in the fitted mixture was set to the maximum number of contexts across all particles in all observation noise sequences—a number that we expect to be at least as great as the number of significant modes in the target density. The weight of component *k* was set proportional to the number of particles (across all observation noise sequences) that had at least *k* contexts. Once we fit the mixture, we kept *M* of the components (where *M* ≤ *K*) and discarded the rest. Specifically, we kept component *k* if the proportion of particles that had at least *k* contexts was more than 0.5 (a simple majority rule), otherwise it was discarded. The reason we fitted more components than we intended to keep was to avoid the potential for unmodeled modes, however small, distorting the fit of components to the largest modes of the target density. The posterior distributions of the drift, retention and bias parameters were extracted in an analogous way.

Note that these mixture components do not have context labels assigned to them. Context labels were assigned in a sequential manner across trials. On trial *t* = 1, only one component existed, and this component was labeled as context 1. Then on each trial *t* = 2,…, *T*, each component was labeled based on its similarity to the components on the previous trial *t* − 1, which were by now labeled. Specifically, we considered all possible permutations of labels and chose the permutation that minimized the KL divergence between components of the same label (on trials *t* − 1 and *t*), summed across labels. Whenever the number of components increased from trial *t* − 1 to *t*, a new label was introduced, and this label was assigned to whichever context had not yet been labeled based on the KL-divergence metric.

To extract the responsibilities (Fig. 1), on each trial we constructed two histograms, one for the responsibilities of instantiated contexts and one for the responsibility of a novel context. Each histogram was formed by binning the responsibilities computed by each particle conditioned on each observation noise sequence, with the counts contributed by observation noise sequence *i* weighted by 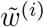. We then considered all valid probability vectors with *M* + 1 elements (i.e., all possible combinations of *M* + 1 bin centers that summed to 1), where *M* − 1 elements were taken from the histogram for instantiated contexts and 1 element was taken from the histogram for a novel context. Each valid probability vector was assigned a weight proportional to the product of the counts in the corresponding *M* + 1 bins. The weighted average of these vectors was taken as the responsibilities. Note that these responsibilities do not have context labels assigned to them (except for the novel context). To assign context labels to them, we considered all possible permutations of labels and chose the permutation that minimized the KL divergence between the true marginal state distribution (defined here as the sum of the predicted state distributions for each context weighted by the responsibilities) and the approximate marginal state distribution (defined here as the sum of the predicted state distributions for each context obtained and labeled as described above, weighted by the responsibilities obtained here). The stationary context probabilities and predicted probabilities were extracted in an analogous way.

To extract the context transition matrix (Fig. S2), on each trial we constructed three histograms of transition probabilities, one for each type of context transition (transitions from an instantiated context to itself [inst-self], transitions from an instantiated context to a different instantiated context [inst-inst] and transitions from an instantiated context to a new context [inst-new]). The bin counts contributed by observation noise sequence *i* were weighted by 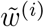. We considered all valid probability vectors with *M* + 1 elements, where *M* − 1 elements were taken from the ‘inst-inst’ histogram, 1 element was taken from the ‘inst-self’ histogram and 1 element was taken from the ‘inst-new’ histogram. As above, each valid probability vector was assigned a weight proportional to the product of the corresponding bin counts. Our goal was to extract *M* vectors, each of which represents a row in an *M* × *M* + 1 transition matrix (where context *M* + 1 is the novel context). Therefore, we clustered the weighted set of probability vectors into *M* components by fitting a mixture of Dirichlet distributions using maximum likelihood estimation of the parameters of each component in the mixture. The number of components in the mixture and the component weights were determined as above for the reduced-component mixture of Gaussians. The mean of each fitted component was used to construct a row of the transition matrix (see below). Note that these transition probabilities do not have context labels assigned to them (except for the novel context) beyond non-specific labels such as self-transition and non-self-transition. To label the contexts, we first considered all permutations of the rows of the matrix along with all permutations of the elements within each row (with the exception of self-transitions and transitions to a new context, which were placed along the diagonal and in the last column of the matrix, respectively). We chose the matrix that minimized the KL divergence between the true marginal state distribution (defined here as the sum of the predicted state distributions for each context, weighted by the sum of the rows of the transition matrix, weighted by the responsibilities on the previous trial) and the approximate marginal state distribution (defined here as the sum of the predicted state distributions for each context obtained and labeled as described above, weighted by the sum of the rows obtained here, weighted by the responsibilities on the previous trial obtained and labeled as described above).

To extract the cue emission matrix (Fig. S2), on each trial we constructed *Q* + 1 histograms of emission probabilities, one for each of the *Q* cues observed so far and one for a novel cue. The bin counts contributed by observation noise sequence *i* were weighted by 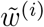. For the *M* rows of the matrix, we considered all valid probability vectors with *Q*+1 elements, where each element was taken from a different histogram. As above, each valid probability vector was assigned a weight proportional to the product of the corresponding bin counts. Our goal was to extract *M* vectors, each of which represents a row in an *M* × *Q* + 1 emission matrix (where cue *Q* + 1 is the novel cue). Therefore, we clustered the weighted set of probability vectors into *M* components by fitting a mixture of Dirichlet distributions using maximum likelihood estimation of the parameters of each component in the mixture. The number of components in the mixture and the component weights were determined as elsewhere. The mean of each fitted component was used as a row of the emission matrix. Note that these emission probabilities have cue but not context labels assigned to them. To assign a context label to each row, we considered all possible permutations of the rows (where row position determines the context label) and chose the permutation that minimized the KL divergence between the true marginal cue distribution (the sum of the cue distributions for each context weighted by the predicted probabilities) and the approximate marginal cue distribution (the sum of the rows obtained here weighted by the predicted probabilities obtained and labeled as described above).

#### Validation of the COIN model

We validated our approximate inference algorithm on synthetic data generated under the generative model. Data was generated in two settings that differed in terms of the upper bound on the number of possible contexts (determined by the truncation level of the stick-breaking process). In the single-context setting only one context was possible (truncation level of 1). In the multiple-context setting, up to 10 contexts were possible (truncation level of 10). For the single-context and multiple-context settings, we generated 4000 and 2000 synthetic data sets, respectively, of 500 time steps duration each. The parameters and hyperparameters used to generate these data sets are shown in Table S1 and were chosen so that the distributions of the numbers of contexts and cues (Fig. S4) were typical of motor learning experiments.

Each data set had a sequence of time-varying latent variables (contexts and states) and observations (state feedback and sensory cues) as well as a set of static parameters for the state of each context (state retention factor and state drift). We applied our inference algorithm to the sequence of observations and at each time step calculated a posterior predictive p-value for each of the time-varying latent variables, observations and parameters. For continuous variables (state feedback, states, parameters), the posterior predictive p-value was calculated by evaluating the cumulative distribution function (CDF) of the predictive probability distribution at the true value of the variable. For discrete variables with integer-valued support (contexts, sensory cues), the posterior predictive p-value was calculated as

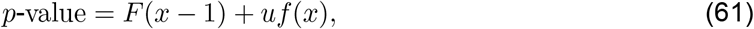

where *f*() is the predicted probability mass function, *F*() is the associated cumulative mass function, *x* is the true value of the variable and 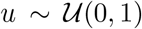 is a uniform random variable on (0, 1). Crucially, if the predictive probability distributions/functions are well calibrated, the distribution of posterior predictive p-values will be uniformly distributed between 0 and 1, and hence the cumulative probability of posterior predictive p-values will lie on the identity line (Figs. S3 and S4).

Under the Markov exchangeable prior of the COIN model, the distribution of context sequences is invariant under permutations of the transitions. Hence context labels (defined by the order in which a context was instantiated/generated) are arbitrary and cannot necessarily be equated between inferred contexts (i.e. contexts instantiated by the inference algorithm) and ground truth contexts (i.e. contexts generated by the generative model) or between particles. This ‘label switching problem’ ^39^ complicates model validation with respect to latent variables that depend on the context (i.e. the state of each context, the parameters for the state of each context and the context itself). We addressed this issue in several ways. In the single-context setting (Fig. S3), we circumvented the label switching problem by limiting the number of contexts to 1. However, this precludes validation of inferences about the context itself. In the multiple-context setting (Fig. S4), we evaluated the state of the current context 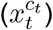 and its associated parameters (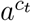 and 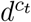) with respect to the CDF of the marginal predictive distributions (i.e. after integrating out the context with respect to the predicted context probabilities). To validate inferences about the context itself, inferred contexts were relabeled by minimizing the Hamming distance (the number of time steps at which context labels differ) between the relabeled context sequence and the true context sequence. Thus we mapped the inferred contexts to a set of labels that maximized the overlap with the ground truth context sequence. This was done using the Kuhn-Munkres (Hungarian) algorithm ^40^ and was performed separately for each particle and at each time step (based on the sequence of contexts up to and including the current time step).

#### Parameter and model recovery

We used the parameters from the fits of the COIN and dual-rate models to the data for each participant in the spontaneous recovery (SR) and evoked recovery (ER) experiments to generate 10 synthetic data sets for each subject from the corresponding paradigm (SR and ER) and model class (COIN and dual-rate). For a given set of parameters in a given experiment, the only source of variability in the dual-rate model across different synthetic data sets was motor noise. In contrast, for the COIN model, in addition to motor noise, sensory noise also provided a source of variability across data sets. We then fit each synthetic data set with both the COIN and dual-rate model as we did with real data (see above).

For model recovery (Fig. S10), we examined the proportion of times the difference in BIC between the COIN and dual-rate fits favored the true (vs. incorrect) model class that was used to generate the data.

For parameter recovery (Fig. S9), we compared the COIN model parameters that were used to generate synthetic data (‘true’ parameters) with the COIN model parameters fit to these synthetic data sets.

#### Responsibility-weighted learning rate

A key prediction of the COIN model is that memory updating should depend on contextual inference (Figs. 1 and 3). This is because the COIN model assumes that only one perturbation—the perturbation associated with the current context—influences the state feedback. Hence, if the current context is known, only the estimate of the perturbation associated with the current context should be updated after observing the state feedback:

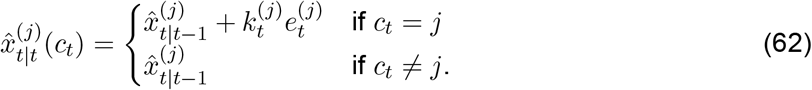

However, in general, the current context is not known. After integrating out the unknown context, the expected value of each update is

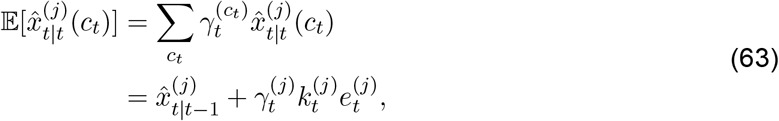

where 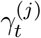 denotes the responsibility of context *j*. The responsibility scales the Kalman gain, producing an effective learning rate 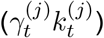 that lies between 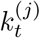 (when 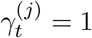, i.e. certain that *c_t_* = *j*) and zero (when 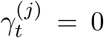, i.e. certain that *c_t_* ≠ *j*). Thus contextual inference is key to Bayes-optimal memory updating (Fig. 1f).

Although the notion that error signals should be scaled by responsibilities is not unique to the COIN model ^1,8^, in the memory updating experiment, we provide the first experimental evidence of this computation (Fig. S11), an achievement made possible by the recently-developed triplet assay of single-trial learning ^9,12^.

#### An analytic approximation to single-trial learning in the COIN model

Here we derive a simple and intuitive approximation to single-trial learning in the COIN model so as to provide insights into the memory updating experiment (Fig. 3). Single-trial learning is defined as

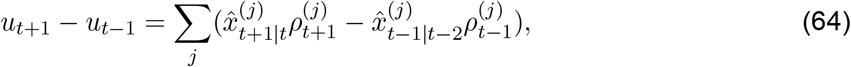

where 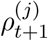 is the predicted probability of context *j* on trial *t* + 1. To aid the derivation, we make use of the following set of simplifying assumptions:

i. There is no decay or drift of state estimates across trials.
ii. All state estimates are zero on the first channel trial of the triplet, which implies that errors on the exposure trial are one.
iii. The Kalman gain is the same for all contexts.

Under these assumptions, single-trial learning can be simplified to

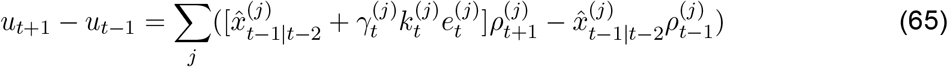

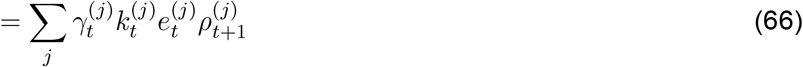

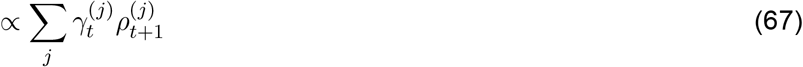

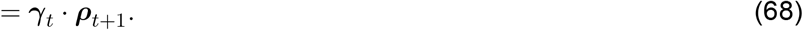

Here ***γ***_*t*_ and ***ρ***_*t*+1_ are vectors of responsibilities and predicted probabilities, respectively. Therefore, single-trial learning is approximately proportional to the dot product of the responsibilities on the exposure trial of the triplet (which determine how much each memory is updated, see Responsibility-weighted learning rate) and the predicted probabilities on the following channel trial (which determine how much each updated memory is subsequently expressed). Intuitively, this dot product is greater when the memories that are updated more are also the ones that are subsequently expressed more. In the memory updating experiment, we confirmed that single-trial learning is indeed well approximated by this dot product (Fig. S11). Moreover, the presentation of a sensory cue on the second channel trial of each triplet allowed us to reveal the effects of differential updating of a single memory by encouraging predicted probabilities to be all-or-none. In this setting, single-trial learning is proportional to the responsibility of the memory on the exposure trial; that is, when 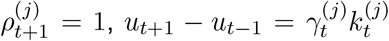. Again, we confirmed that this indeed is the case (Fig. S11)

The effects of force fields, sensory cues and context transition probabilities on single-trial learning can be explained in a unified manner using this simple dot-product metric. In the memory updating experiment (Fig. 3), ***ρ***_*t*+1_ is constant across the four triplet types, as we present the same sensory cue on the channel trials of all triplets, but 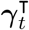 varies, as we present different combinations of force fields and sensory cues on the exposure trials of the triplets. In contrast, in the environmental-consistency experiments (Fig. 4c and Fig. S14), 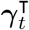 is constant, as the same force field is presented on the exposure trial of the triplets and there are no sensory cues, but ***ρ***_*t*+1_ varies, as the context transition probabilities differ across the environments.

## Supplementary Text

### Mathematical analysis of spontaneous and evoked recovery in the COIN model

Here we develop a mathematical analysis of how the main features of spontaneous and evoked recovery emerge in the COIN model. Specifically, the main features we wish to explain are that spontaneous recovery is 1. non-monotonic, with a smooth but transient increase in adaptation, followed by decay, 2. which asymptotes (at least within the time scale of the experiment) above zero, and evoked recovery shows 3. very rapid (almost instantaneous) increase to a higher level of adaptation than spontaneous recovery, followed by monotonic decay, 4. which also asymptotes above zero.

In general, state and contextual inference in a switching state-space model, such as the COIN model, is analytically intractable. However, inference can be performed analytically under the following assumptions: (i) there are no state feedback observations (as on channel trials); (ii) the inferred parameters of the state and context transition dynamics are constant; and (iii) the number of contexts does not change. In this special case, state estimates are updated according to the state dynamics ascribed to each context:

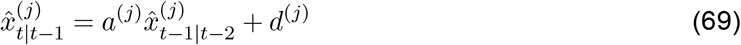

and context probabilities (***ρ***) are updated (independently of the states) according to the context transition matrix (**Π**):

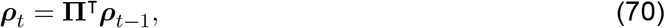

Assumptions (i)-(iii) are at least approximately true during the channel trial phase of the spontaneous and evoked recovery paradigms, i.e. when our main explicanda occur. Specifically, assumption (i) is true as there is no state feedback. Assumption (ii) is approximately true as the inferred parameters governing the state and context transition dynamics are updated relatively little over the timescale relevant for spontaneous and evoked recovery late in learning. Assumption (iii) is approximately true as new contexts tend not to be inferred when state feedback is omitted.

Based on these approximations, we simulated state and contextual inference during the channel phase of the spontaneous and evoked recovery paradigms (Fig. S5). We ran the simulations with two contexts using parameters *a*^(1)^ = *a*^(2)^ = 0.95, *d*^(1)^ = −*d*^(2)^ = 0.0075 and

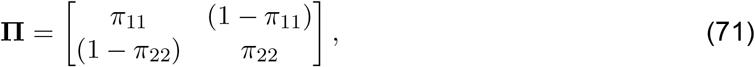

where *π*_11_ = 0.999 and *π*_22_ = 0.9, reflecting the fact that nearly all transitions in the experiment are self-transitions, and that the context associated with *P*^+^ has been experienced more often than the context associated with *P*^−^.

For spontaneous recovery, on trial (immediately following *P*^−^), the state estimates associated with *P*^+^ (context 1, red) and *P*^−^ (context 2, orange) are equal but opposite (Fig. S5a), and the context probabilities are equal (Fig. S5b, solid lines). Hence, adaptation is initially at baseline (Fig. S5c, solid line). Then, based on Eq. 69, the state estimates converge exponentially (at the same rate) to their steady-state values, 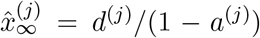. In particular, for context 1, this means a monotonically decreasing decay towards a non-zero asymptote (Fig. S5a, red) because experience in *P*^+^ is compatible with a positive steady-state (which we incorporated by our choice of a positive drift rate, *d*^(*j*)^, in Eq. 69). At the same time, based on Eq. 70, the predicted probabilities converge exponentially to their values under the stationary distribution, lim_*t*→∞_ *p*(*c_t_* = *j*) (Fig. S5b, solid lines). Context 1 is more probable than context 2 under the stationary distribution as *P*^+^ was experienced for more trials than *P*^−^ during the experiment. Hence, the predicted probability of context 1 monotonically increases (Fig. S5b, solid red). The net result of these updates is that there is an initial rise in adaptation due to the increasing contribution of the state associated with context 1, followed by a fall in adaptation toward a non-zero baseline due to the decay of this state toward a non-zero steady-state (Fig. S5c, solid line). Therefore, the classic non-monotonic nature of spontaneous recovery arises because the dynamics of contextual inference (responsible for the initial rise in adaptation) are faster than the dynamics of state inference (responsible for the subsequent fall in adaptation). Critically, as long as the inferred state dynamics reflect the statistics of the experiment in which, by design, the true state of each context never changes (the *P*^+^ and *P*^−^ perturbations are constant), the dynamics of contextual inference are bound to be faster than the dynamics of state inference, and the steady-state of adaptation (on the time scale of the experiment) to be above zero. Thus, non-monotonic spontaneous recovery (feature 1) with a decay that does not reach zero (feature 2), as seen in the experimental data (Fig. 2c), is a robust feature of the COIN model. Indeed, the simulation of the full model without the approximations we introduced above for analytical tractability also shows such spontaneous recovery (Fig. 2b, bottom right, and c; also for individual subjects whose data shows spontaneous recovery, Fig. S8) with all three main properties that our analysis here suggests are key for obtaining this result. Specifically, (a) state estimates associated with *P*^+^ and *P*^−^ approximately cancel at the beginning of the *P*^c^ phase (when weighted with their corresponding context probabilities) and then monotonically converge to a positive and negative steady-state, respectively (Fig. 2b, bottom left); (b) the corresponding context probabilities may be similar initially but then diverge, such that the probability associated with *P*^+^ grows toward a near-one steady-state, while that associated with *P*^−^ shows the opposite trend, decaying toward a near-zero baseline (Fig. 2b, top right); (c) the dynamics of contextual inference are markedly faster than those of state estimation (Fig. 2b, cf. top right and bottom left).

For evoked recovery, we assume the learner is certain they are in context at the end of the second evoker (*P*^+^) trial, and from then on, during the *P*^c^ trials, their contextual inferences evolve according to the same dynamics as in spontaneous recovery. For a direct comparison with spontaneous recovery, we also kept everything else (parameters, state estimates) identical to the simulation of spontaneous recovery. (This included ignoring more subtle differences in state inferences between the two paradigms; cf. Fig. 2b and d, bottom left.) Because context is also much more probable than context 2 under the stationary context probabilities (see above), the context probabilities did not change much with updating from their initial values (Eq. 70), and so the probability of context 1 and context 2 respectively remained high and low throughout the simulation (Fig. S5b, dashed; 1 cf. Fig. 2d, top right). Hence, adaptation largely reflected the dynamics of the state of context 1 (Fig. S5a, red; cf. Fig. 2d, bottom left), decaying exponentially from a level of adaptation that was necessarily higher than that reached in spontaneous recovery to a non-zero asymptote (Fig. S5c, dashed). Thus, the model produced a rapid, strong (feature 3), and long-lasting (feature 4) recovery, as seen in both the experimental data (Fig. 2e) and in the full model (Fig. 2d, bottom right, and e; Fig. S8). In other words, our analysis suggests that evoked recovery can be understood as a limiting case of spontaneous recovery when – spurred by evoker trials – the dynamics of contextual inference converge very rapidly to the stationary context probabilities. As such, the same three key properties that underlie spontaneous recovery ensure that the main features of evoked recovery also robustly emerge in the COIN model.

### Working memory in the COIN model

A working memory task performed just before the channel trial phase has been shown to interfere with spontaneous recovery, and in fact to create an effect that is reminiscent of evoked recovery, such that *P*^+^ adaptation returns immediately to a high level following *P*^−^, already on the first *P*^c^ trial (Fig. S15a, Ref. 19). In the dual-rate model, this effect has been attributed to a selective diminishing of the adaptation of the fast learning process ^19^. We simulated the COIN model with the parameters obtained from the fit to the average spontaneous and evoked recovery data sets (also used in Fig. 2b,d). The COIN model reproduces the effect by modeling the working memory task as selectively abolishing the memory of the context probabilities on the last *P*^−^ trial (Fig. S15b-d). This means that on the first *P*^c^ trial, predicted context probabilities are based on general knowledge of how frequently different contexts are expected to be encountered in the future (stationary distribution), rather than on which contexts are likely to follow the context specifically encountered on the last trial (compare colored circles between middle right panels of Fig. S15c and d). Because *P*^+^ has been the most frequent trial type, the probability of its associated context under the stationary distribution is very high, and hence there is a strong re-expression (evoked recovery) of the memory for this context. This suggests that the belief over contexts may require working memory for maintenance.

### Explicit versus implicit learning in the COIN model

Recent studies have shown that motor learning has both explicit and implicit components which exhibit markedly different time-courses ^20,22^. For example, in a paradigmatic example using a visuomotor rotation task, a measure of explicit learning was obtained by asking participants to report the direction in which they planned to move prior to moving, and implicit learning was then measured as the difference between the actual direction they moved and this explicit judgement ^21^. In a spontaneous recovery paradigm, explicit learning showed non-monotonic behavior during the *P*^+^ phase, fast increase followed by slow decay (Fig. S16a). In contrast, implicit learning showed slower and monotonic increase during the *P*^+^ phase. Due to these differences in the form of adaptation, explicit and implicit learning have been suggested to correspond respectively to the fast and slow processes of the dual-rate model ^20^. However, this mapping is unable to account for the rapid drop and recovery of supposedly slow implicit learning seen during the subsequent *P*^−^ and *P*^c^ phases.

In order to simulate these experiments, we adapted the COIN model to account for a critical difference between visuomotor and force-field learning: visuomotor but not force-field learning (which is the primary paradigm we use to test the predictions of the COIN model in the main text) introduces a discrepancy between the hand’s proprioceptive and visual location. Due to this discrepancy, a fundamental credit-assignment problem arises ^8^ as to whether the observed cursor deviation is due to a perturbation on the motor system or a bias (miscalibration) in the sensory system. This was naturally captured in the COIN model by introducing a bias between the state and sensory feedback as another latent parameter in each context, which was learned together with the parameters that govern the evolution of the state in that context (Methods). We hypothesized that participants would have explicit access to the state representing their belief about the visuomotor rotation, but that they would not have access to their sensory bias which would reflect the implicit component of learning.

We simulated the COIN model with the parameters obtained from the fit to the average spontaneous recovery and evoked data sets (also used in Fig. 2b,d and Fig. S15) plus an additional parameter representing the standard deviation of the prior on the bias (Methods). Fig. S16c, d & e show the bias, state and predicted probability for each context. The average bias across contexts weighted by the predicted probabilities (Fig. S16f) showed a slow monotonic increase during the *P*^+^ phase with a drop and recovery during the *P*^−^ and *P*^c^ phase. As hypothesized, the profile is very similar to that of the implicit component of learning (Fig. S16a-b, light green). However, the average state across contexts (Fig. S16g) did not show the experimentally observed characteristic overshoot of the explicit component (Fig. S16a, dark green). Instead, examining the state of the context with the highest responsibility (Fig. S16d, colored bar in the bottom, and thin black line, also shown as dark green line in (Fig. S16b) revealed that it had a strikingly similar time course to the explicit component of learning (Fig. S16a & b dark green). This is because the state and the bias interact competitively within a context to account for the total state feedback, and hence as the bias estimate increases, the state estimate decreases, giving rise to the characteristic non-monotonicity. As the experimental definition of explicit and implicit components guarantees that they sum to total adaptation (see above), we also defined motor output in the model as the sum of the explicit (state of the context with the highest responsibility) and implicit components (Fig. S16b, solid pink). Taken together, this version of the COIN model reproduced the important qualitative features of explicit, implicit, and total adaptation in the experiment (compare Fig. S16a and b). 1 (Although there were quantitative differences, e.g. in the overall speed of learning, note that all but one parameter were fit to rather different force-field learning experiments and so a quantitatively precise match could not be expected.) In particular, the different time courses of explicit versus implicit components arose naturally in the model. This is because, in the COIN model, parameters (i.e. bias) are assumed to be constant over the lifetime of a context, whereas states can change dynamically, and thus their estimates are updated more slowly (Fig. S16c, f) than those of states (Fig. S16d, g) – inherently giving rise to multiple time scales of learning. Moreover, the average bias across contexts in the COIN model (Fig. S16f, cyan, and b, light green) also tracked the rapid drop and recovery of implicit learning during the *P*^−^ and *P*^c^ phase (Fig. S16a, light green) that the dual-rate model cannot explain. This arises from the same contextual inference-based mechanism that also underlies other aspects of spontaneous recovery (Fig. 2). Specifically, the rapid fall in the implicit component of learning during the *P*^−^ phase is due to the increased expression of the associated context (Fig. S16e, orange) that has a negative bias (Fig. S16c, orange). On entering the *P*^c^ phase, there is a re-expression of the context associated with *P*^+^ (Fig. S16e, red) that has a positive bias (Fig. S16c, red).

Interestingly, in order for the COIN model to be consistent with experimental data, our definition of total adaptation in this experiment (average bias plus the state of *just one context*, the most responsible one) needed to be different from what would have been directly consistent with the way it is originally defined in the model (the net predicted state feedback, here corresponding to the average bias plus the average state *across all contexts*). However, this experiment was also conducted differently from the other experiments we modeled. In particular, in this paradigm, an explicit judgment was solicited at the beginning of each trial before motor output was required. We reasoned that the explicit commitment of where they will aim would determine where participants eventually aim in their motor output (measured as total adaptation), thus explaining why only the reported state (corresponding to the explicit judgment in the model) and not the average state is reflected in motor output. This is in line with previous studies showing that an explicit commitment affects subsequent decision making ^41^. Moreover, this reasoning made a further prediction: in a (control) variant of the same visuomotor rotation experiment in which no explicit judgments are solicited, total adaptation should have a different time course as it now should reflect the average state not just the explicitly reported state. This did indeed seem to be the case in the data (albeit slightly, not reaching statistical significance ^21^): learning of *P*^+^ was slower and adaptation to *P*^−^ was not as completed as in the original version of the task. These differences were qualitatively reproduced by the COIN model when total adaptation was modeled as usual, using the average state across contexts (Fig. S16a-b, dashed pink).

Importantly, learning a measurement bias is equivalent to performing sensory recalibration, which is known to occur during adaptation to a visuomotor rotation. For example, after learning a visuomotor rotation with their right hand, a participant can be asked to use their non-adapted left hand to point to where they sensed their right hand was at the end of a reach ^42^. Consistent with a sensory recalibration, participants incorrectly estimate the location of their right hand location, pointing closer to where the cursor was than to the actual location of their right hand.

In summary, rather than mapping explicit and implicit learning to fast and slow processes, which only differ quantitatively, the COIN model suggests that they may map to qualitatively different components of learning: state variables and parameters (in this case, bias), respectively. This mapping then implies that participants may be explicitly aware of the state (most probable rotation angle) but not the bias (discrepancy between proprioceptive and visual location of the hand). The inability to report the average motor state estimate across contexts, and instead report the state of the most probable context as their explicit judgment, is in line with many studies that examine motor and perceptual judgments and find clear dissociations (e.g. ^43,44^).

### Comparison with other theories of contextual inference

There are deep analogies between the context-dependence of learning in the motor system and other learning systems, both in terms of their phenomenologies and the computational problems they are trying to solve. The segmentation of continuous experience into discrete “contexts” or 1 “events” is a fundamental aspect of a variety of learning systems, such as associative learning ^45^ and episodic memory ^46^. Recently, such context-dependent learning has been successfully formalized within the framework of Bayesian non-parametric models ^6,47–49^. However, these insights have not been brought to bear on the motor system. Instead, theories of motor learning either did not have a notion of context ^3,9,50^ or used heuristics for how motor learning parses experience into discrete contexts, and how it expresses and updates the memories laid down for each context ^2^. In those few cases in which contextual motor learning was considered within a principled probabilistic framework, the generative models underlying learning were not sufficiently rich (e.g. they lacked state or context transition dynamics) to be able to express fundamental properties of the environment that are also critical for explaining a number of learning phenomena ^8,23^.

Table S3 summarizes the ability of dominant single-context and multiple-context models to explain the main data sets we have modeled. The COIN model performs contextual inference in a more principled and comprehensive way than previous models of contextual learning (both of motor and other memories), which was key to explaining the experimental data we and others collected. The COIN model is principled because it uses a coherent Bayesian inversion of a well-defined, rich generative model of the environment to compute context probabilities. Critically, these graded context probabilities implement a soft assignment of observations to contexts, such that all memories are partially updated and expressed on each trial, which was crucial in explaining memory expression in spontaneous recovery (Fig. 2), memory updating in single-trial learning (Fig. 3), savings (Fig. 4a), anterograde interference (Fig. 4b), changes in apparent learning rates with environmental consistency (Fig. 4c). In contrast, previous models perform hard context assignments such that only one context is expressed ^2^ or updated ^2,6^ on each trial and thus are unable to account for many of these phenomena. The COIN model is also comprehensive in that contextual inferences are based on all relevant information that is available to the brain: sensory cues, state feedback, and the history of contexts (encoded in context transition probabilities). In contrast, previous models either did not learn context probabilities at all ^2^, which was crucial for explaining spontaneous recovery, or did not exploit the Markovian nature of context transitions when learning transition probabilities ^6,8^, which was crucial for explaining environmental-consistency effects (Fig. 4c & Fig. S14). For example, the Dirichlet process Kalman filter (DP-KF) model proposed by Gershman et al. ^6^ learns transition probabilities by counting the number of times each context has been experienced, rather than the number of transitions between each pair of contexts (as in the COIN model). Hence, the DP-KF only learns the overall probabilities of contexts (independent of the previous context). Therefore, as the three groups in the environmental-consistency study experienced *P*^+^ and *P*^−^ equally often (and only the transition probabilities varied between the groups, Fig. 4c), this model cannot account for the differences in single-trial learning seen in the data. Yet other models, namely the MOSAIC model ^23^, learn context transition probabilities but, unlike the COIN model, they do so in a non-hierarchical way such that the transition probabilities learned in one context do not generalize to other contexts (i.e. each row of the context transition matrix is updated independently of all other rows). This results in this model not being able to capture anterograde interference (Fig. 4b) as the transition probabilities learned in the *P*^+^ context will have no effect once in the *P*^−^ context and therefore cannot affect the expression of *P*^−^. Finally, models that do not include contextual inference at all are not able to explain both spontaneous and evoked recovery correctly ^3,9^ 1 (Fig. 2c,e), and are inherently unable to predict the effect of explicit contextual sensory cues on memory updating Fig. 3.

---

**Figure S1.**
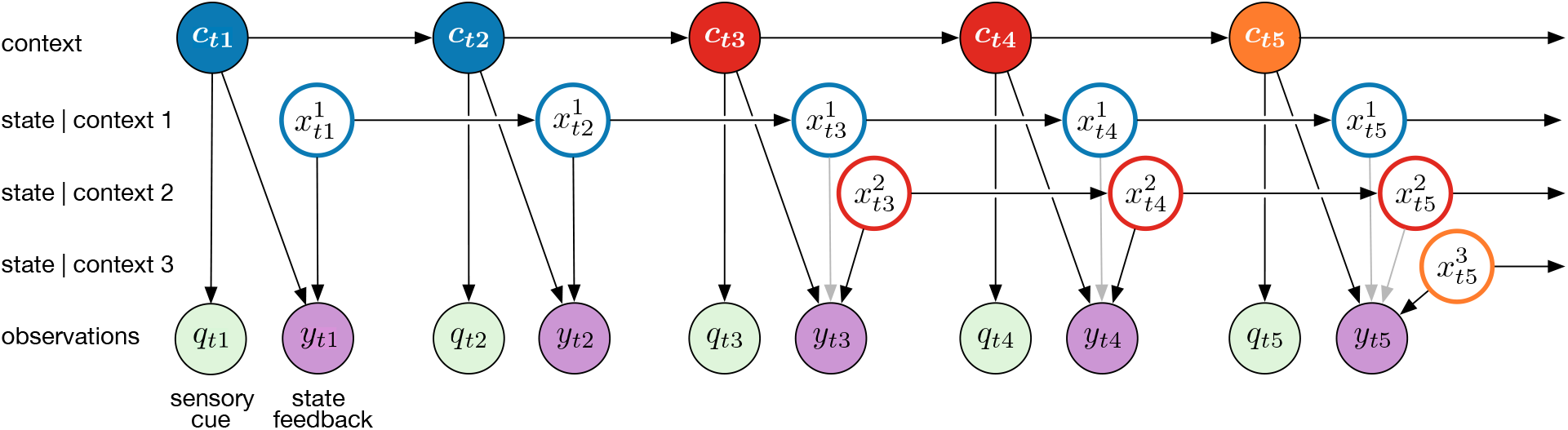
COIN model. Graphical model of the generative process of the COIN model. A (potentially) infinite number of discrete contexts, *c_t_* (colors), exist that transition as a Markov process. Each context *j* is associated with a state, 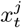, that evolves as a linear-Gaussian (stochastic) dynamical system independent of the states of the other contexts. For clarity, only three states are shown and each state is only shown from when the context associated with it first becomes active, even though all states exist at all times. The current context can lead to the emission of a discrete sensory cue, *q_t_*, and also generates state feedback, *y_t_*, associated with its state (i.e. only the state associated withe active context contributes to the state feedback; black vs. gray arrows). The context and the states are hidden from the learner who must infer them from observed sensory cues and state feedback.

**Figure S2.**
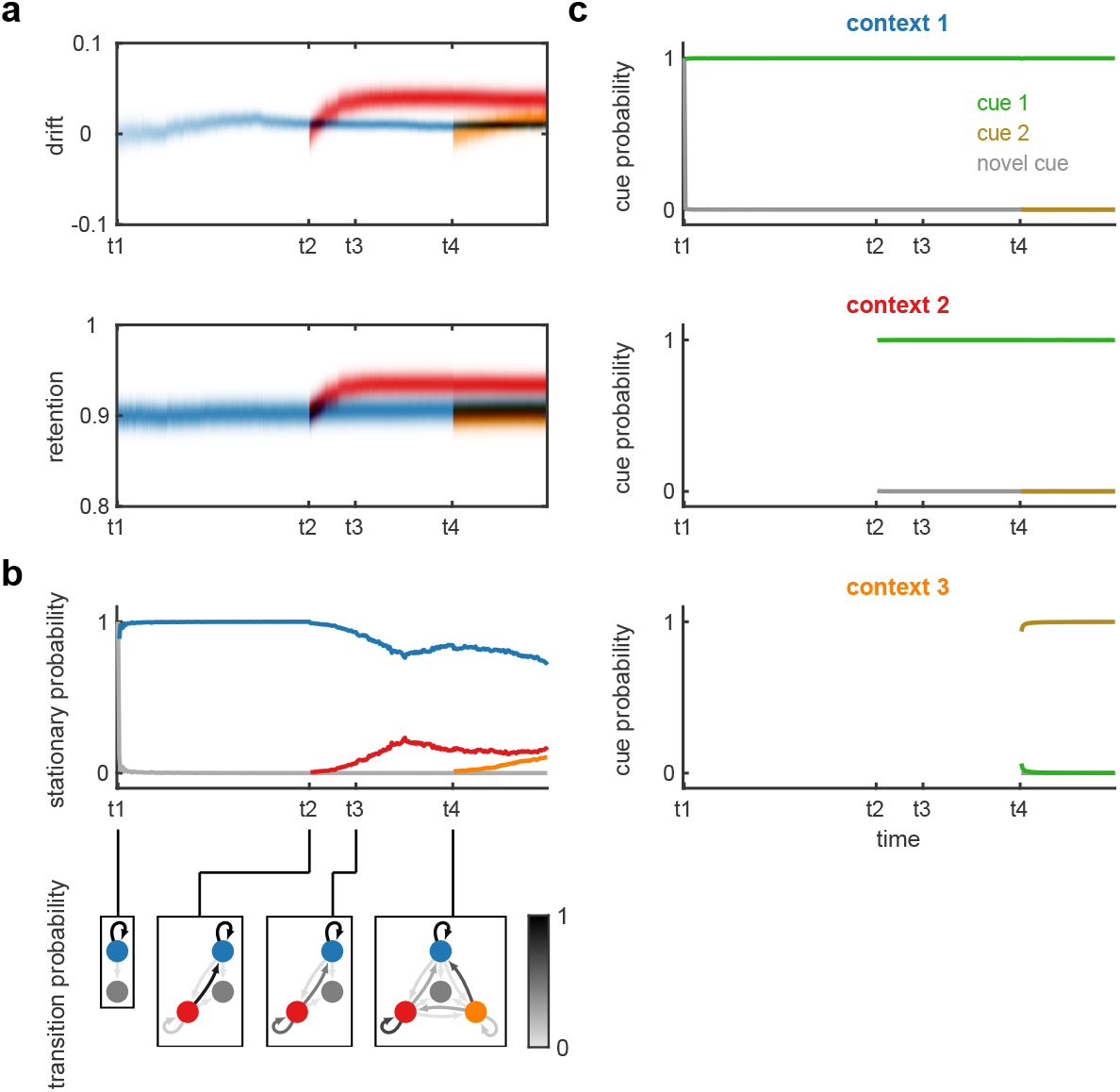
Parameter inference in the COIN model. In addition to inferring states and contexts (shown in Fig. 1), the COIN model also infers the parameters of within-context state dynamics (**a**) as well as the parameters governing transitions between contexts (**b**) and cue emissions for contexts (**c**). **a,** Posterior distribution of drift and retention parameters for the three instantiated contexts (colors as in Fig. 1). Note that drift and retention are estimated to be larger for the red context that is associated with the largest perturbation. **b,** Stationary probability for each instantiated context (line colors) and the novel context (gray) representing the expected proportion of time spent in each context given the current estimate of the transition matrix. Insets show Markov chain representations of the inferred transition probability matrices at four key time points with arrow shading showing the transition probabilities. Note that the model infers these transition probability matrices (at all time points) but the stationary probabilities illustrated here are not explicitly computed by the model: we derived them from the transition probability matrices to show the model’s current prediction for the overall probability of encountering each context in the future. **c,** Inferred cue emission probabilities for the three instantiated contexts (panels) and cues (line colors). Note that full (Dirichlet) posterior distributions are computed over both transition (**b**) and cue emission probabilities (**c**) in the model, but for clarity here we only show the means of these posterior distributions.

**Figure S3.**
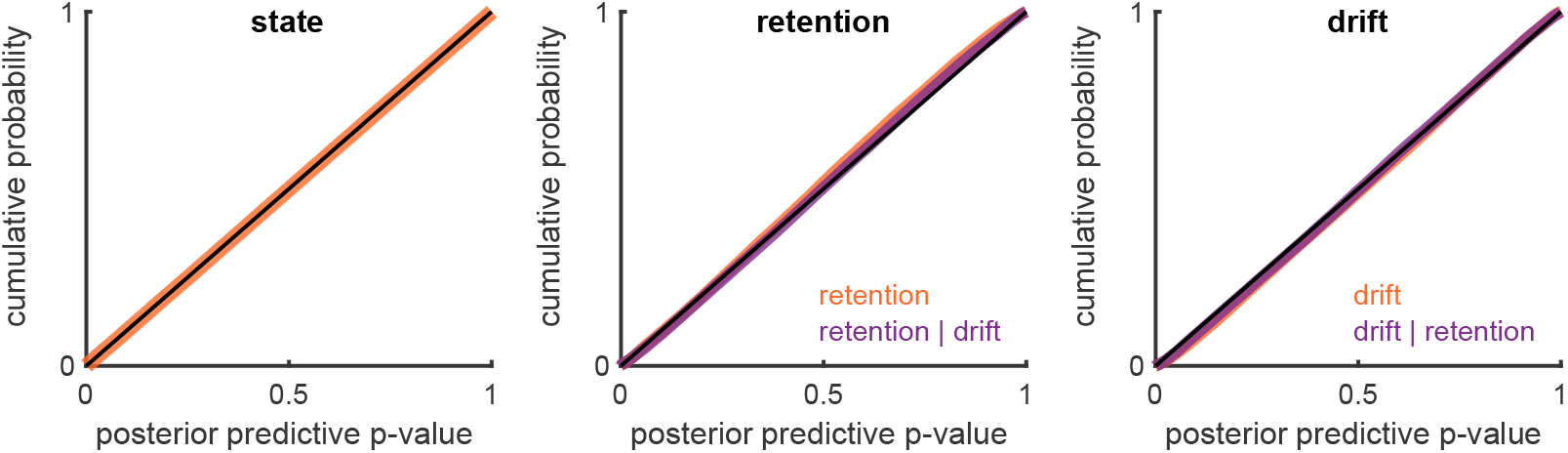
Validation of the inference algorithm of the COIN model with a single context. We computed inferences in the COIN model with a single context based on synthetic inputs generated by its generative model (Fig. S1). Plots show the cumulative distributions of posterior predictive p-values of the state variable (left), and the parameters governing its dynamics (retention, middle; and drift, right). The posterior predictive p-value is computed by evaluating the c.d.f. of the model’s posterior over the given quantity at the true value of that quantity (as defined by the generative model). Empirical distributions of posterior predictive p-value were collected across 4000 simulations (with different true dynamics parameters), with 500 time steps in each simulation (during which the true state changes, but the dynamics parameters are constant). Note that although true dynamics parameters do not change during a simulation, inferences in the model about them will still generally evolve, and so a new posterior p-value is generated in each time step even for these quantities. If the model implements well-calibrated probabilistic inference under the correct generative model, all these empirical distributions should be uniform. This is confirmed by all cumulative distributions (orange and purple curves) approximating the identity line (black diagonal). Orange curves show posterior predictive p-values under the corresponding marginals of the model’s posterior. To give additional information about the model’s joint posterior over dynamics parameters, we also show the posterior predictive p-value (cumulative) distribution of each parameter conditioned on the true value of the other one (purple curves).

**Figure S4.**
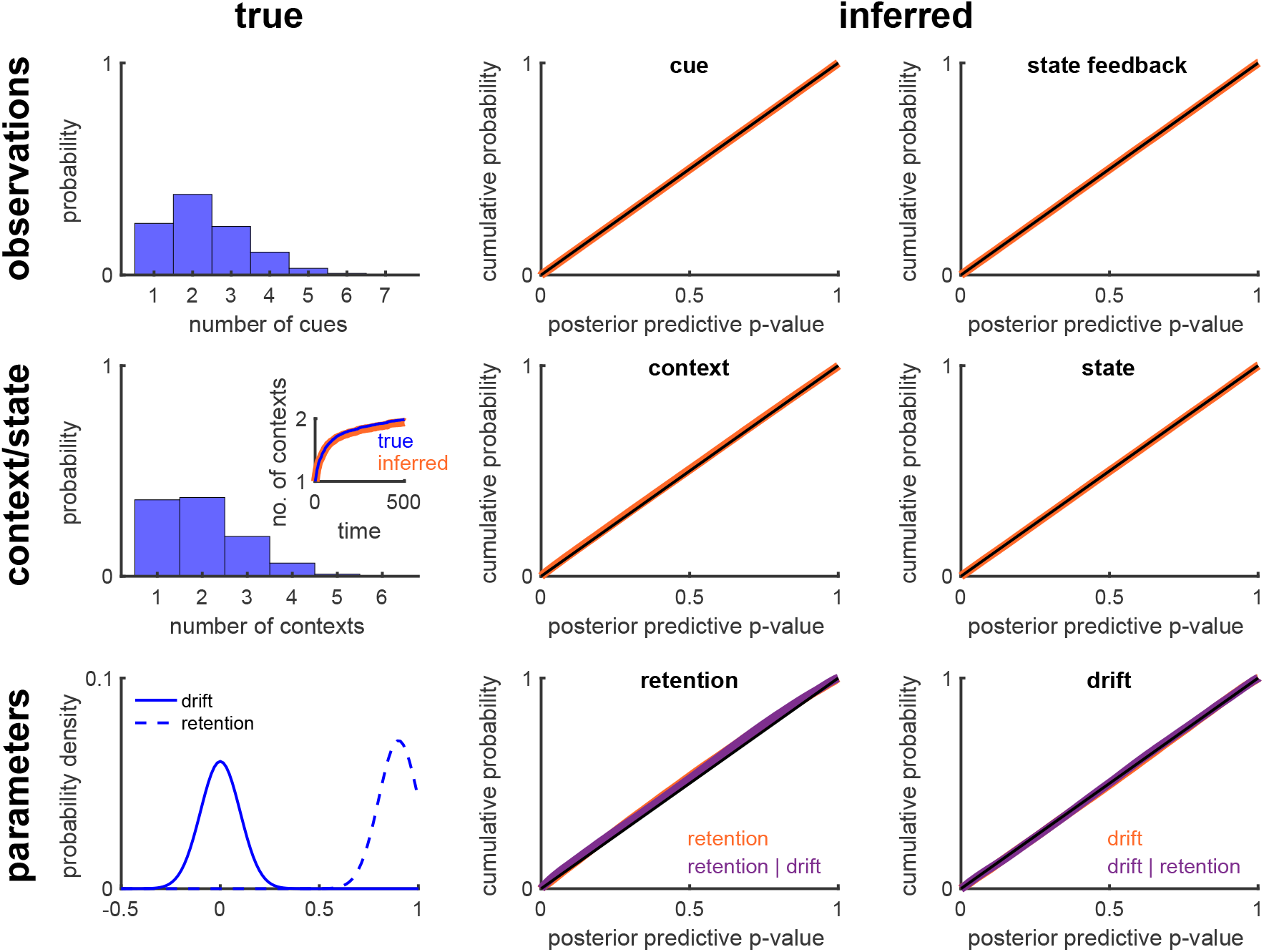
Validation of the inference algorithm of the COIN model with multiple contexts. Simulations with synthetic data as in Fig. S3 but with multiple contexts allowed both during data generation and inference. Empirical distributions of posterior predictive p-value were collected across 2000 simulations (with different true retention and drift parameters), with 500 time steps in each simulation (during which not only states evolve but also contexts transition, and sometimes new contexts are created). Left column shows the true distributions of sensory cues, contexts and parameters. Inset shows the growth of the number of contexts over time in both during generation (true) and inference (inferred). Middle and right columns show the cumulative probabilities of the posterior predictive p-values (pooled across data sets and time steps) for the observations (top row), contexts and state (middle row) and parameters (bottom row). To calculate the posterior predictive p-values for the context, inferred contexts were relabeled by minimizing the Hamming distance between the relabeled context sequence and the true context sequence using the Hungarian algorithm (see Suppl. Mat.). For the parameters, the posterior predictive p-values were calculated with respect to both the marginal distributions (retention and drift) and the conditional distributions (retention drift and drift retention) as in Fig. S3. The cumulative probability curves approximate the identity line (thin black line) showing that the inferred posterior probability distributions are well calibrated.

**Figure S5.**
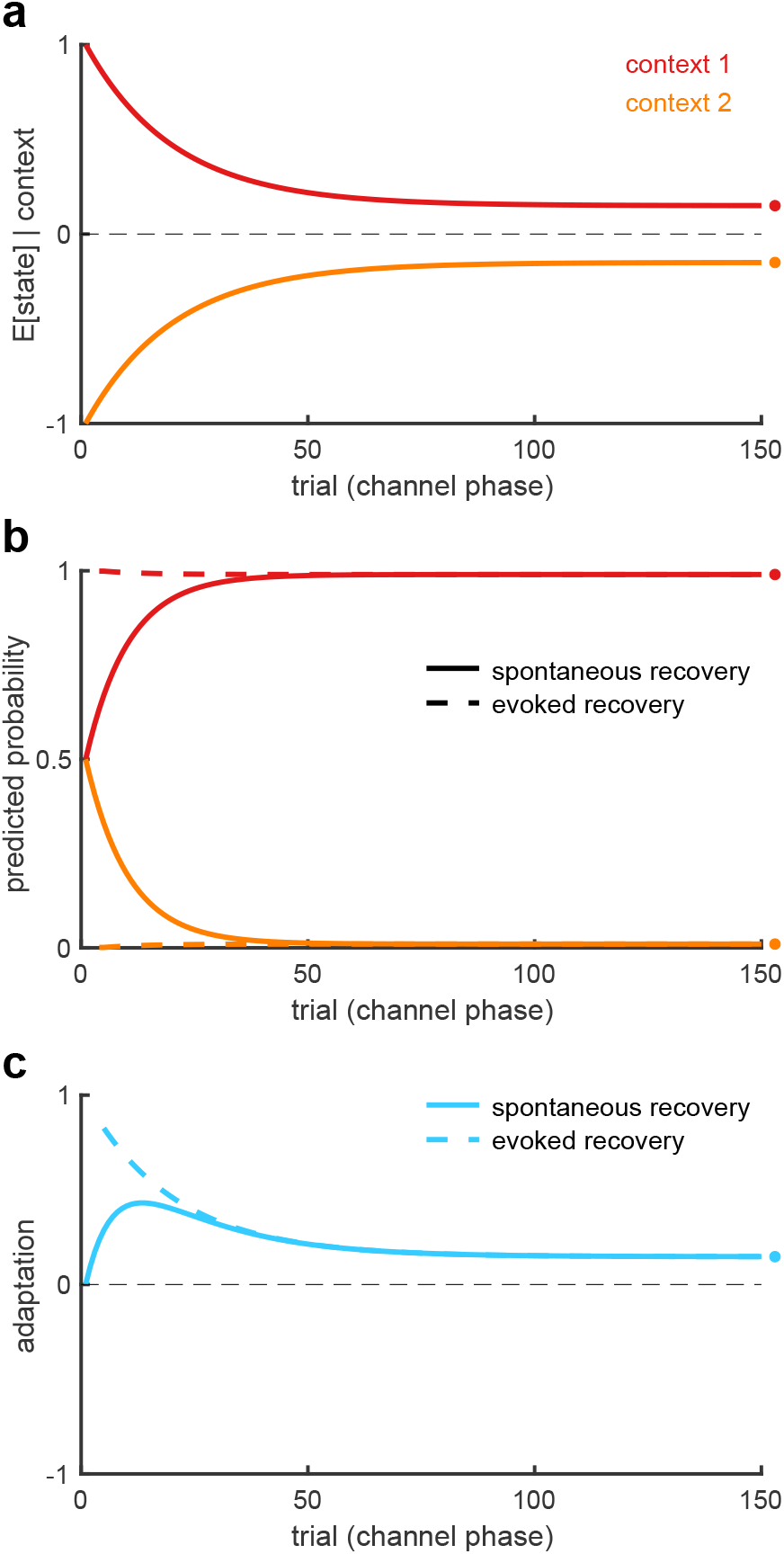
Mathematical analysis of spontaneous and evoked recovery. The channel phase of spontaneous and evoked (after the two *P*^+^ trials) recovery simulated in a simplified setting (Suppl. Text) with two contexts that are initialized to have equal but opposite state estimates (**a**) and equal (spontaneous recovery, solid) or highly unequal (evoked recovery, dashed) predicted probabilities (**b**). For the two contexts the retention parameters are assumed to be constant and equal, and the drift parameters are assumed to be constant, of the same magnitude but opposite sign. Mean adaptation (**c**), which in the COIN model is the average of the state estimates (**a**) weighted by the corresponding context probabilities (**b**), shows the classic non-monotonic pattern of spontaneous recovery (solid, cf. Fig. 2b-c) and the characteristic abrupt rise of evoked recovery (dashed, cf. Fig. 2d-e). Note that although in the full model, state estimates are different between evoked and spontaneous recovery following the two *P*^+^ trials, here we assumed they are the same (no separate solid and dashed lines in **a**) for simplicity and to demonstrate that the difference in mean adaptation between the two paradigms (**c**) can be accounted for by differences in contextual inference alone (**b**, cf. Fig. 2b and d, top right insets). Circles on the right show steady-state values of inferences and the adaptation. Note that in both paradigms, adaptation is predicted to decay to a non-zero asymptote (see also Fig. S6).

**Figure S6.**
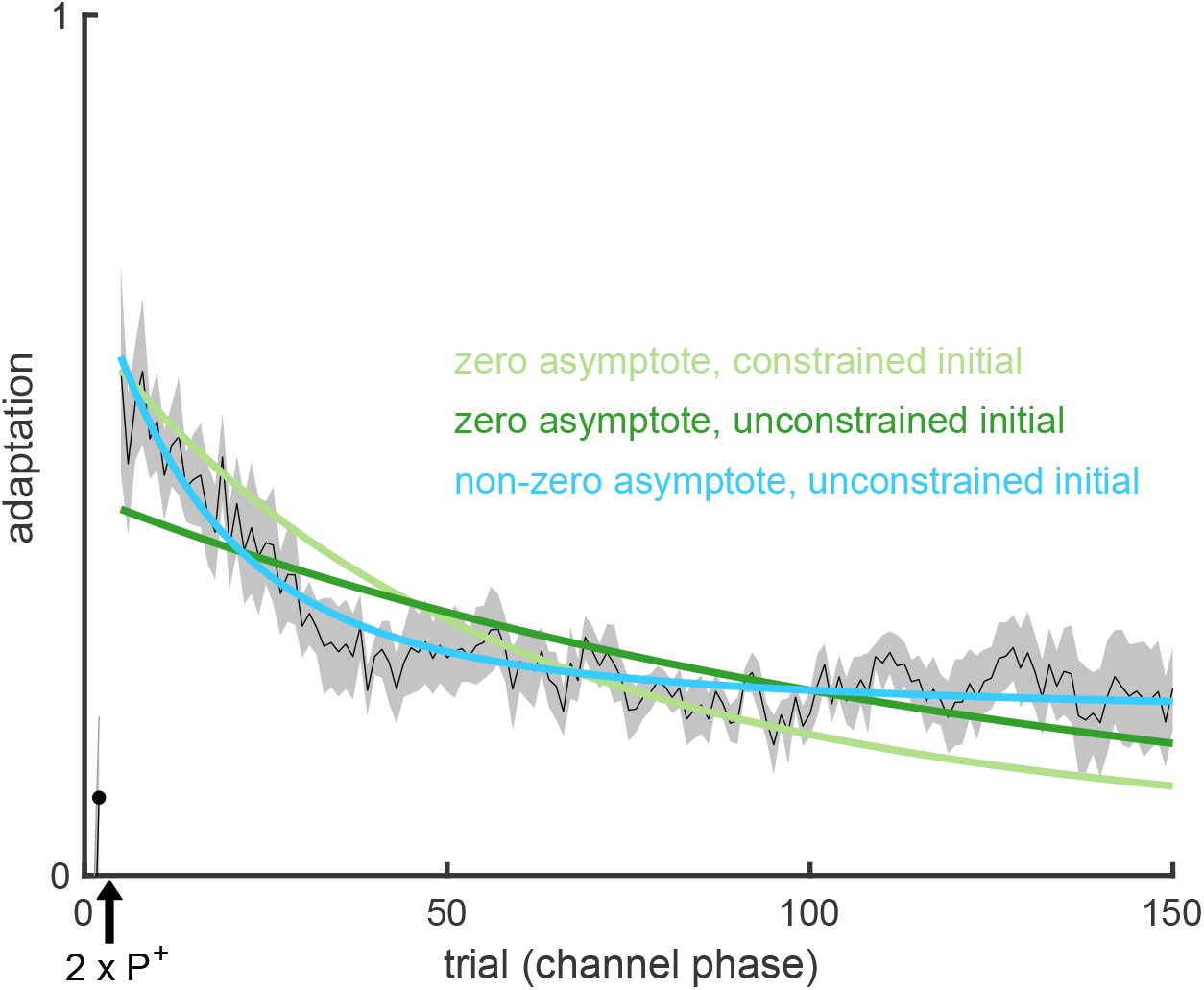
Evoked recovery does not decay exponentially to zero. According to the COIN model, adaptation in the channel trial phase of evoked recovery can be approximated by exponential decay to a non-zero (i.e. positive) asymptote (Fig. 2e, Fig. S5, Suppl. Text). To test this prediction, we fit an exponential function that either decays to zero (light and dark green) or decays to a non-zero (constrained to be positive) asymptote (cyan) to the adaptation data of individual participants in the evoked recovery group after the two *P*^+^ trials (black arrow). The two zero-asymptote models differ in terms of whether they are constrained to pass through the datum on the first (channel) trial or not. The mean fits across participants for the models that decay to zero (green) fail to track the mean adaptation (black, ± s.e.m. across participants), which shows an initial period of decay followed by a period of little or no decay. The mean fit for the model that decays to a non-zero asymptote (cyan) tracks the mean adaptation well and was strongly favored in model comparison (Δ group-level BIC of 944.3 and 437.7 compared to the zero-asymptote fits with constrained and unconstrained initial values, respectively). Note that fitting to individual participants excludes the confound of finding a more complex time course (e.g. one with non-zero asymptote) only due to averaging across participants that each show a different simple time course (e.g. all with zero asymptote but different time constants).

**Figure S7.**
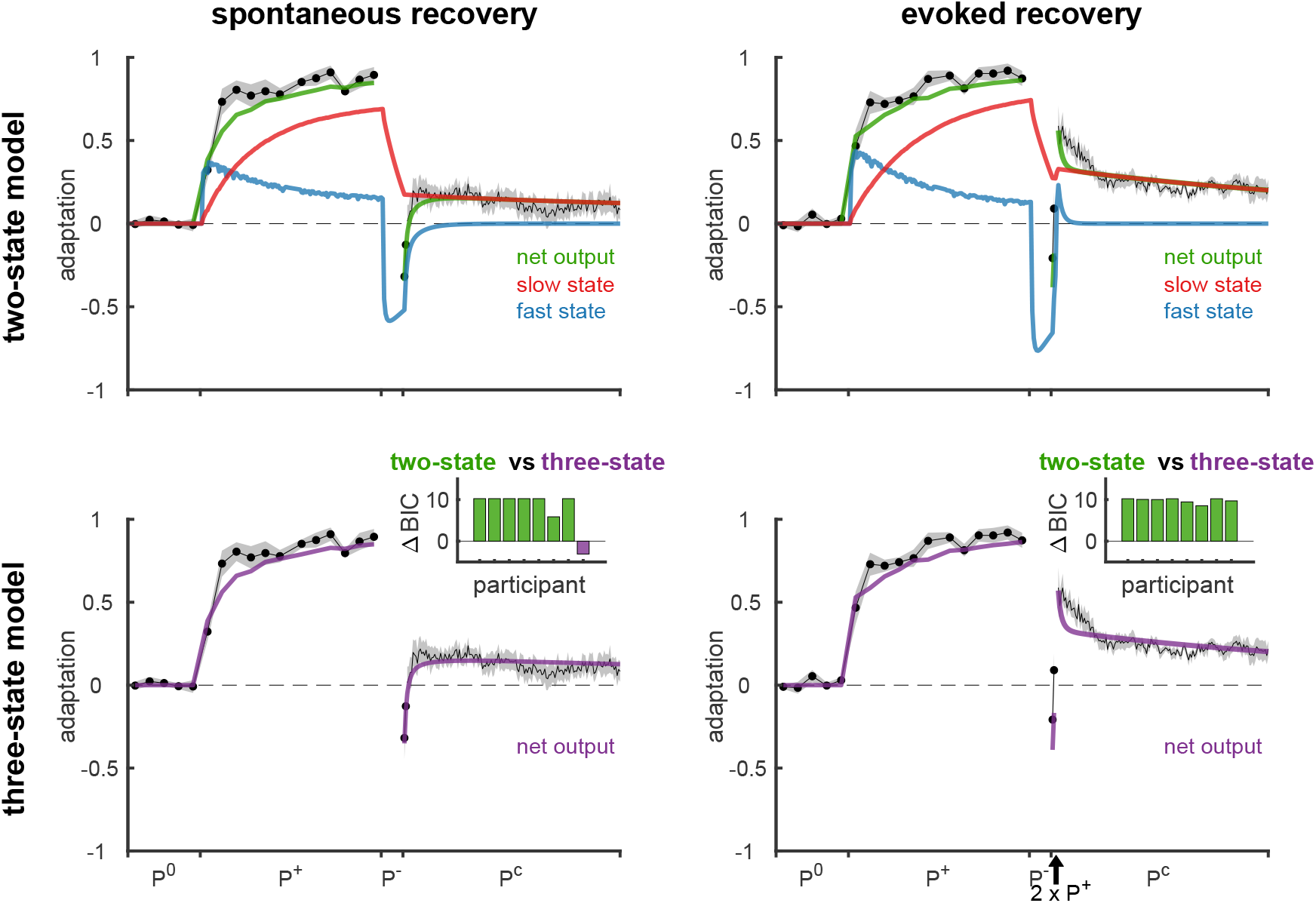
State-space model fits to adaptation data from the spontaneous and evoked recovery groups. Solid lines show the mean fits across participants of the two-state model (top row) and the three-state model (bottom row) to the spontaneous recovery (left column) and evoked recovery (right column) data sets. Mean ± s.e.m. adaptation on channel trials shown in black (same as in Fig. 2c and e). Insets show differences in BIC between the two-state model and the three-state model for individual participants (positive values in green indicate evidence in favor of the two-state model, and negative values in purple indicate evidence in favor of the three-state model). At the group level, the two-state model was far superior to the three-state model (group-level BIC of 64.2 and 78.4 for the spontaneous and evoked recovery groups, respectively). Individual states are shown for the two-state model (top, blue and red). Both the fast and slow processes adapt to *P*^+^ during the extended initial learning period. The *P^−^* phase reverses the state of the fast process, but not of the slow process, so that they cancel when summed resulting in baseline performance. Spontaneous recovery during the *P*^c^ phase is then explained by the fast process rapidly decaying, revealing the state of the slow process that has remained partially adapted to *P*^+^. Note that this explanation assumes that the fast and slow processes contribute equally to the motor output at all times. This is fundamentally different from the expression and updating of multiple context-specific memories in the COIN model, which are dynamically modulated over time according to ongoing contextual inference.

**Figure S8.**
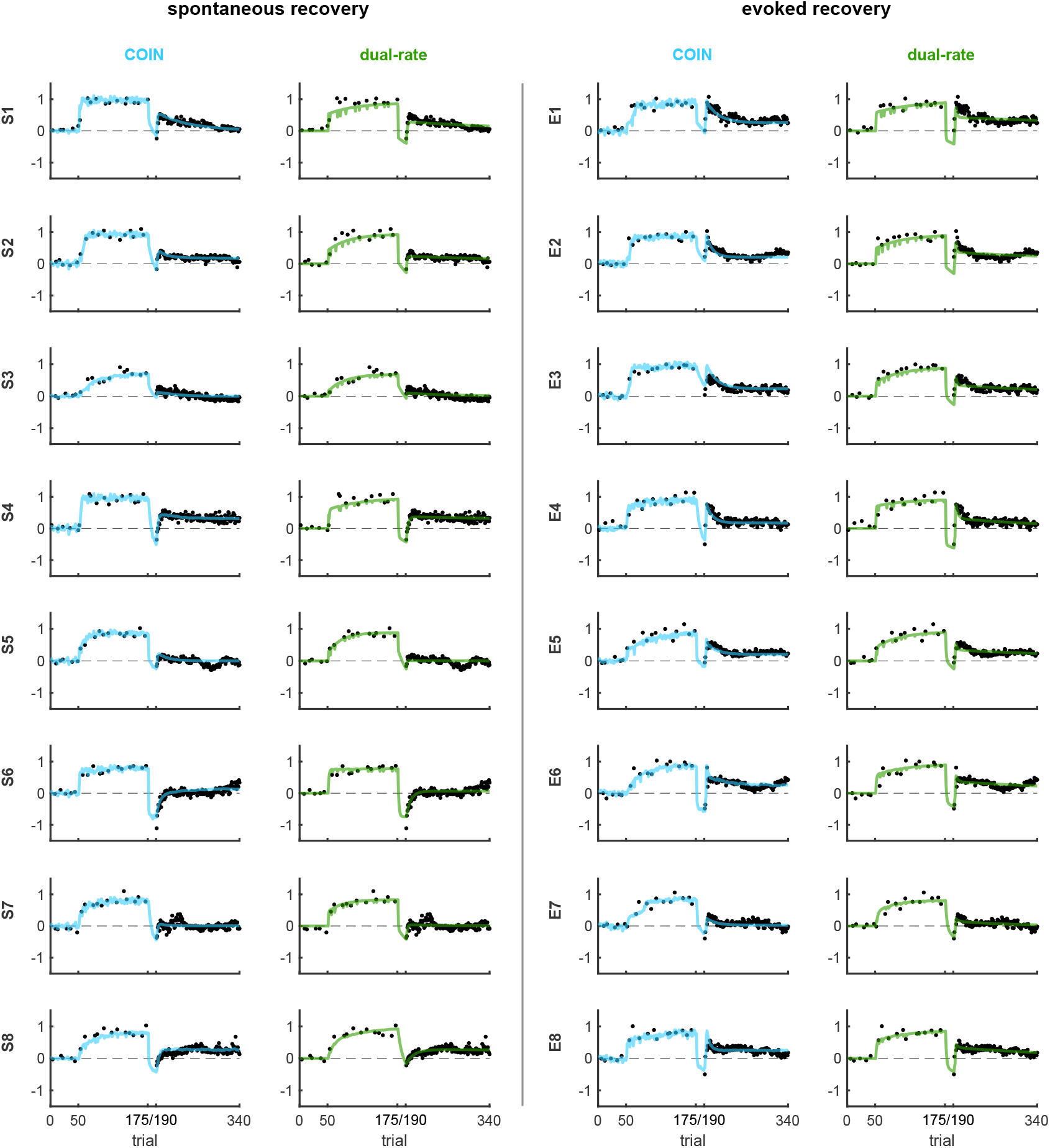
COIN and dual-rate model fits for individual participants in the spontaneous and evoked recovery groups. Data and model predictions are shown for individual participants as in Fig. 2c and e for across-participant averages. Participants in the S and E groups are ordered by decreasing BIC difference between the dual-rate and COIN model (i.e. S1’s and E1’s data most favor the COIN model), as in insets of Fig. 2c and e. Note that the COIN model can account for much of the heterogeneity of spontaneous (e.g. from large in S1 to minimal in S6) and evoked recovery (e.g. from large in E1 to minimal in E7).

**Figure S9.**
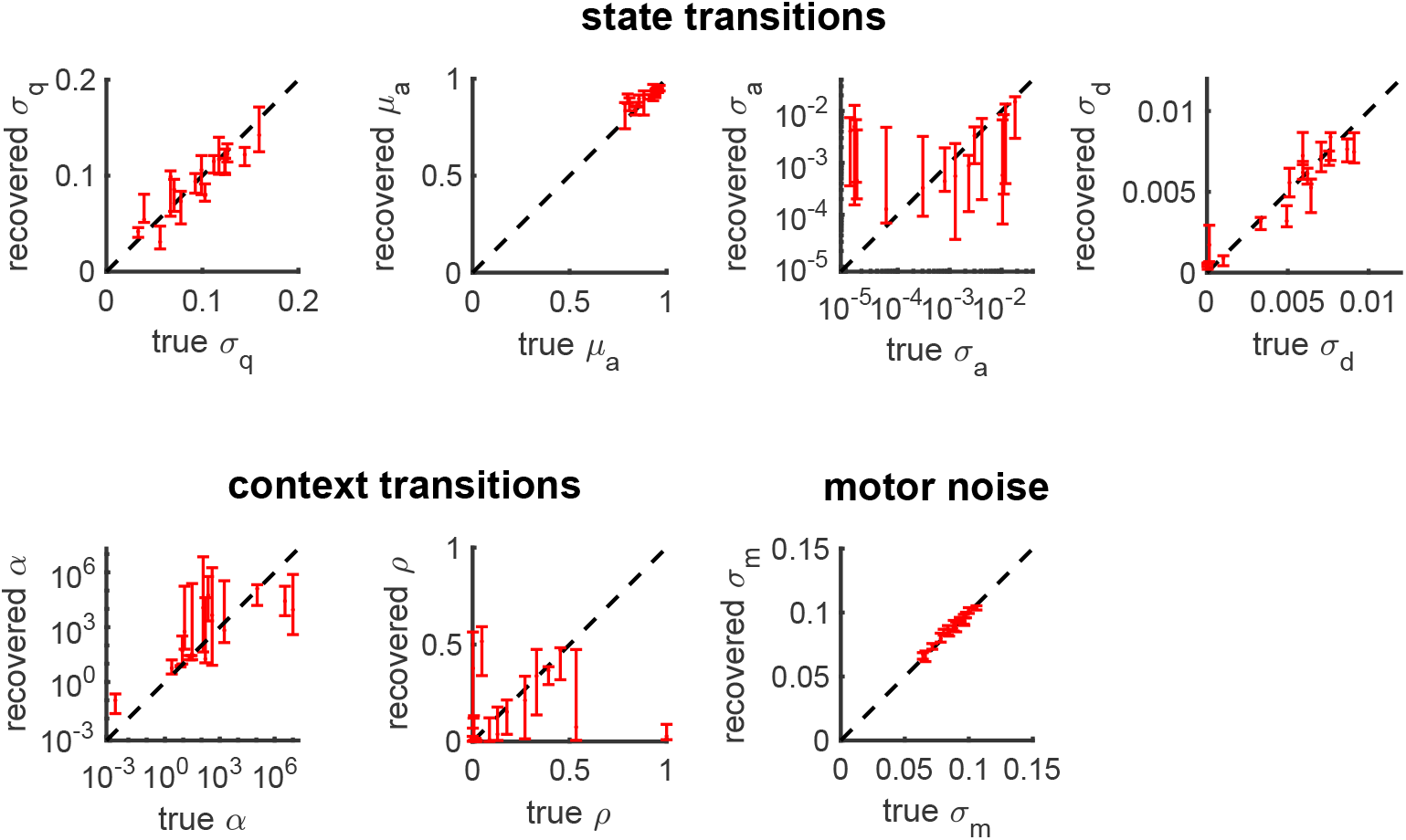
Parameter recovery in the COIN model. Plots show the COIN model parameters that were recovered (y-axes) from fits to 10 synthetic data sets generated with the COIN model parameters (true, x-axes) obtained from the fits to each participant in the SR and ER experiments (Methods). Vertical bars show the interquartile range of the recovered parameters for each participant. While several parameters are recovered with good accuracy (*σ*_q_, *μ*_a_, *σ*_d_, *σ*_m_), others are not (*α*, and in particular *σ*a and *ρ*). We expect that with richer paradigms and larger data sets, all parameters would be recovered accurately. Most importantly, despite partial success with recovering individual parameters, model recovery shows that recovered parameter sets taken as a whole can be used to accurately identify whether data was generated by the dual-rate or COIN mechanism (Fig. S10). Note that we make no claims about individual parameters in this paper as our focus is on model class recovery.

**Figure S10.**
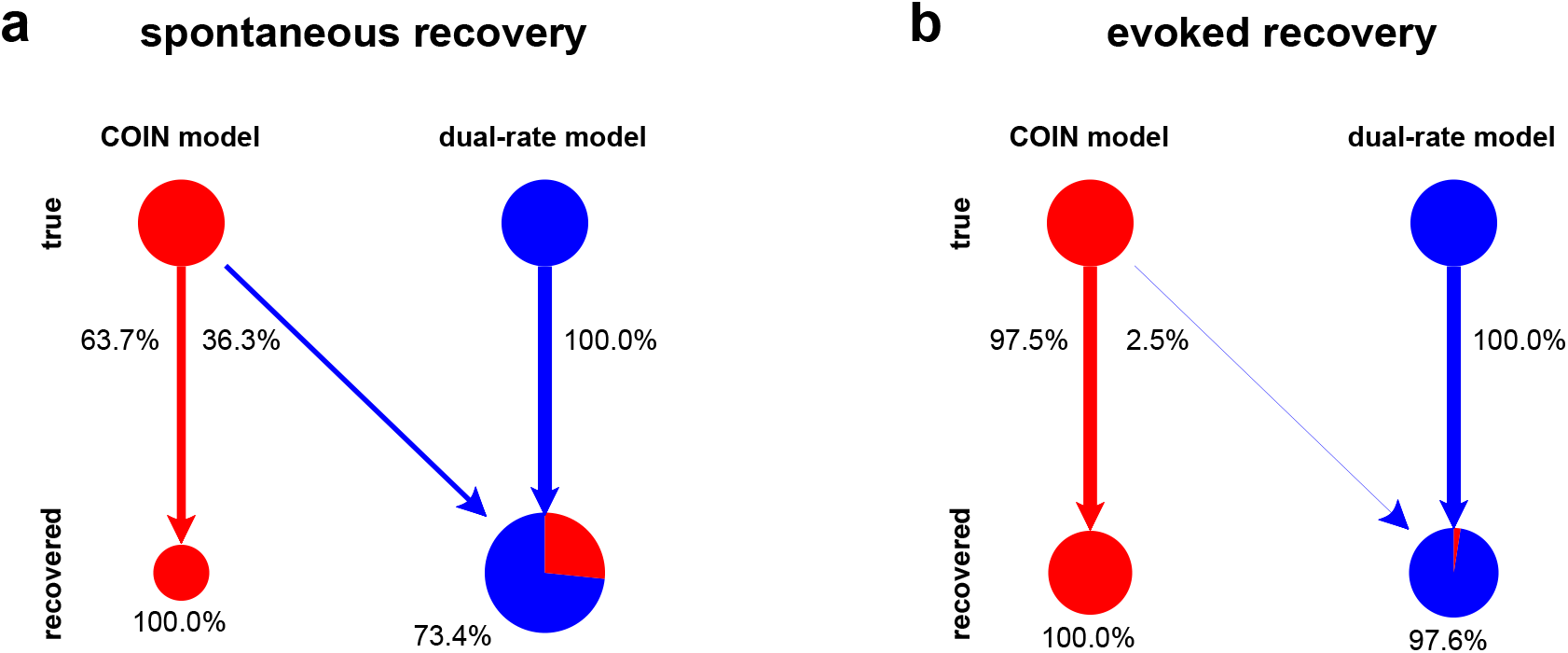
Model recovery for spontaneous (a) and evoked recovery experiments (b). Synthetic data sets were generated using one of two models (COIN model, red; dual-rate model, blue) and then the same model comparison method that we used on real data (Fig. 2c, e, insets) was used to recover the model that generated each synthetic data set (Methods). Arrows connect true models (used to generate synthetic data, disks on top) to models that were recovered from their synthetic data (pie-chart disks at bottom). Arrow color indicates identity of recovered model, arrow thickness and percentages indicate probability of recovered model given true model. Bottom disk sizes and pie-chart proportions respectively show total probability of recovered model and posterior probability of true model given recovered model (assuming a uniform prior over true models), with percentages specifically indicating posterior probability of the correct model.

**Figure S11.**
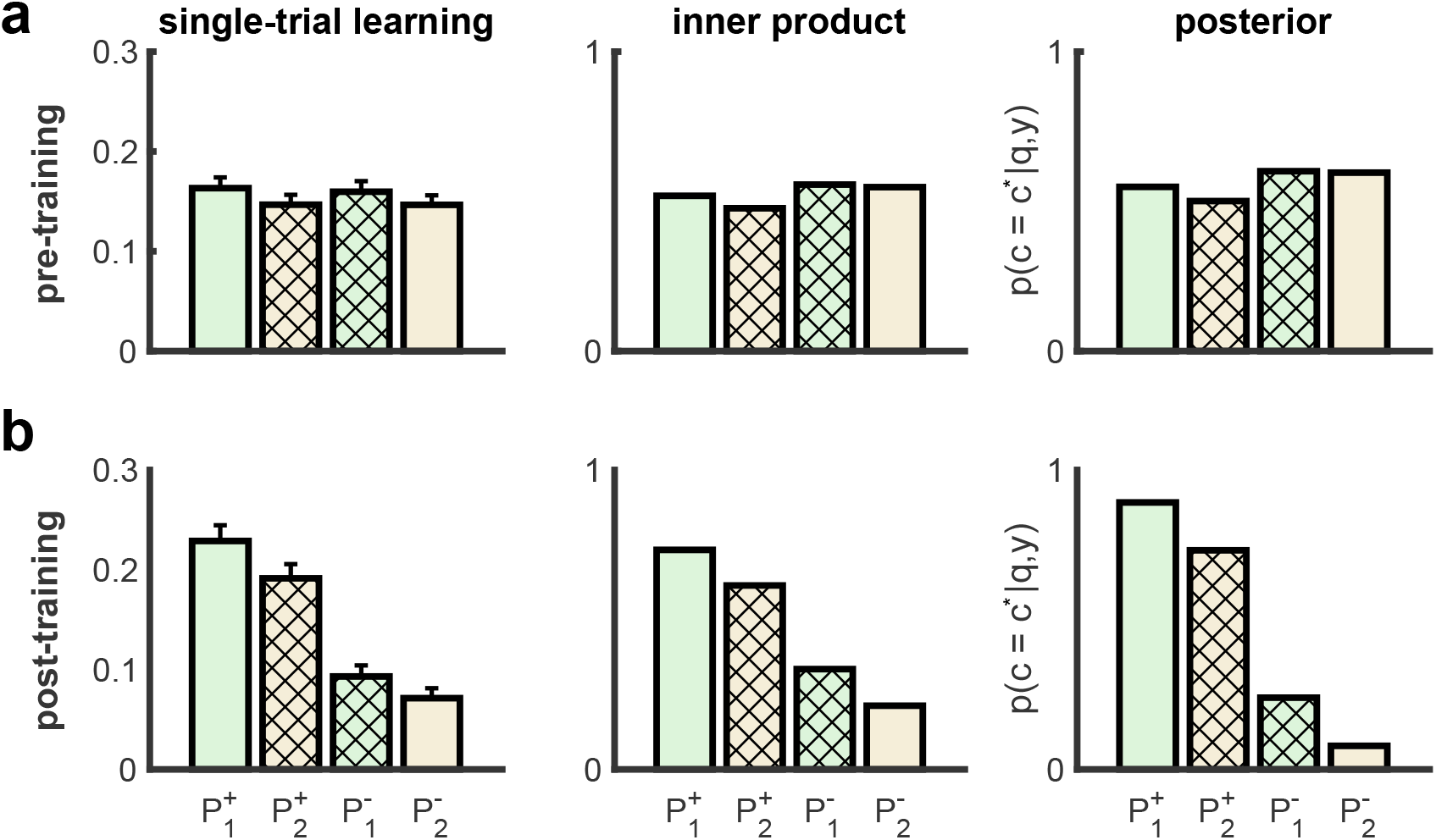
Single-trial learning reflects posterior contextual inference and can be approximated as a dot product. **a,** Single-trial learning for the four cue-perturbation triplets before the training phase in the memory updating experiment. Data shows mean ± s.e.m. across participants. Single-trial learning is approximately proportional to a dot product between the vector of responsibilities on the exposure trial of the triplet and the vector of predicted probabilities on the subsequent channel trial (see Suppl. Mat. for derivation). The model posterior on the exposure trial is also shown for the context that was predominantly expressed on the final channel trial of the triplet (*c*\*). **b,** Single-trial learning as in (**a**) after training.

**Figure S12.**
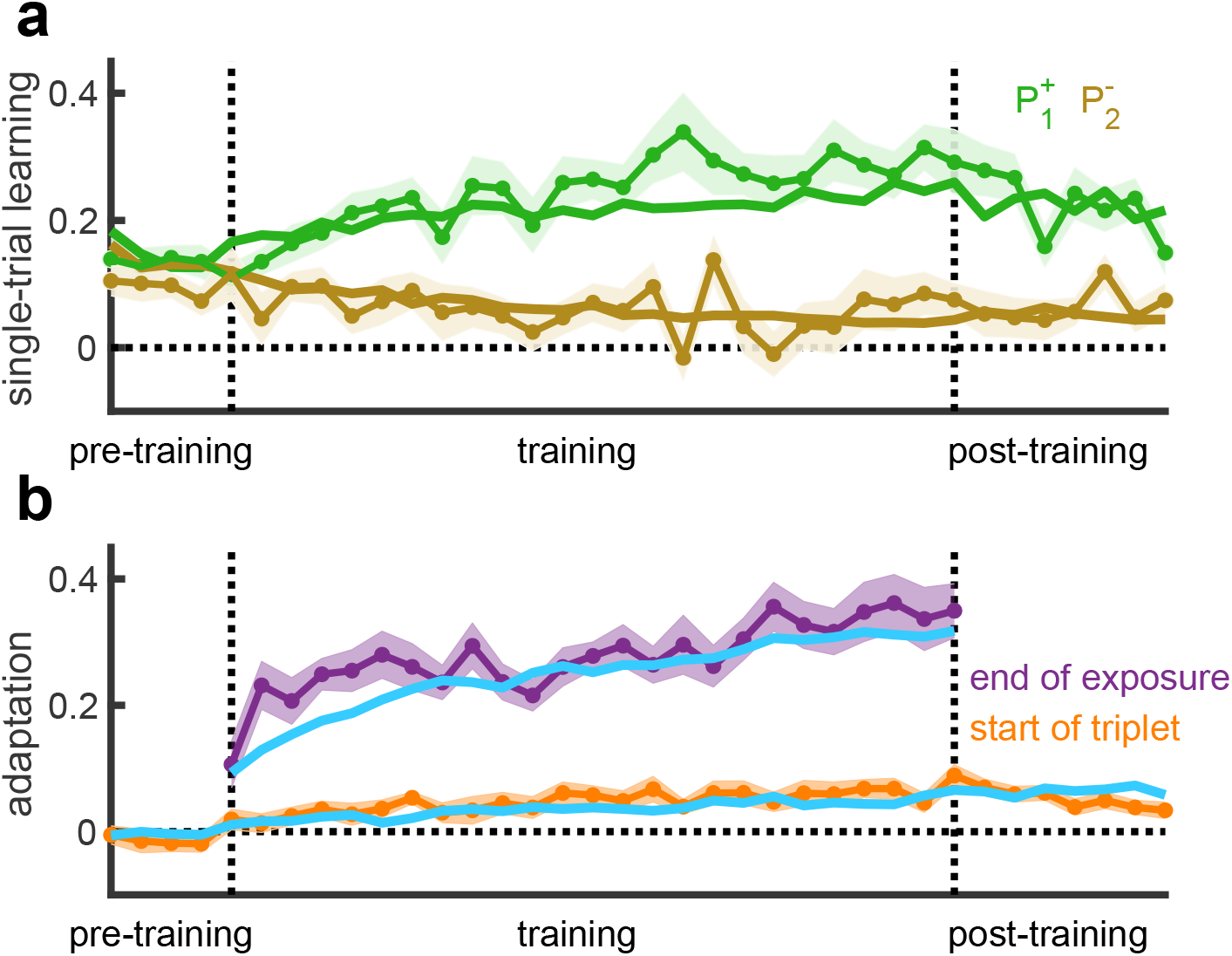
Memory updating experiment time course of learning. **a**, Single-trial learning on triplets that were consistent with the training contingencies. Data (mean ± s.e. across participants) with mean of COIN model fits. Positive learning reflects changes in the direction expected based on the force field of the exposure trial (an increase following *P*^+^, and a decrease following *P^−^*). **b**, Adaptation on channel trials at the end of each block of training (purple) and on the first channel trial of triplet within each block after washout trials (*P*^0^). Data is mean ± s.e. across participants and lines show mean of COIN model fits.

**Figure S13.**
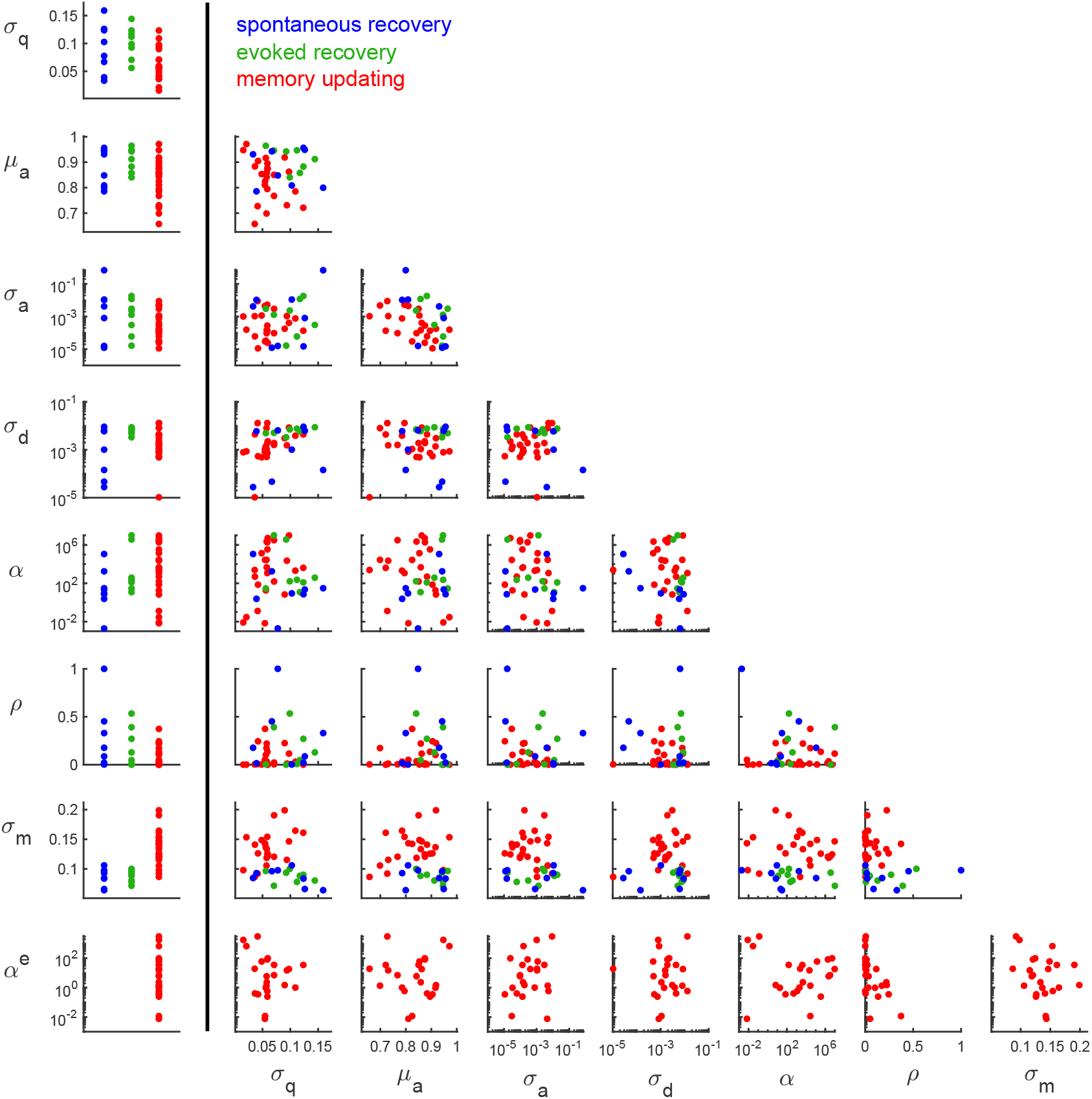
Parameters of the COIN model fit to individual participants. Left column: Individual participant’s parameters in the spontaneous recovery (blue), evoked recovery (green) and memory updating (red) experiments. Right: scatter plots for all pairs of parameters for the three groups. The overlapping clouds suggest parameters are similar across experiments.

**Figure S14.**
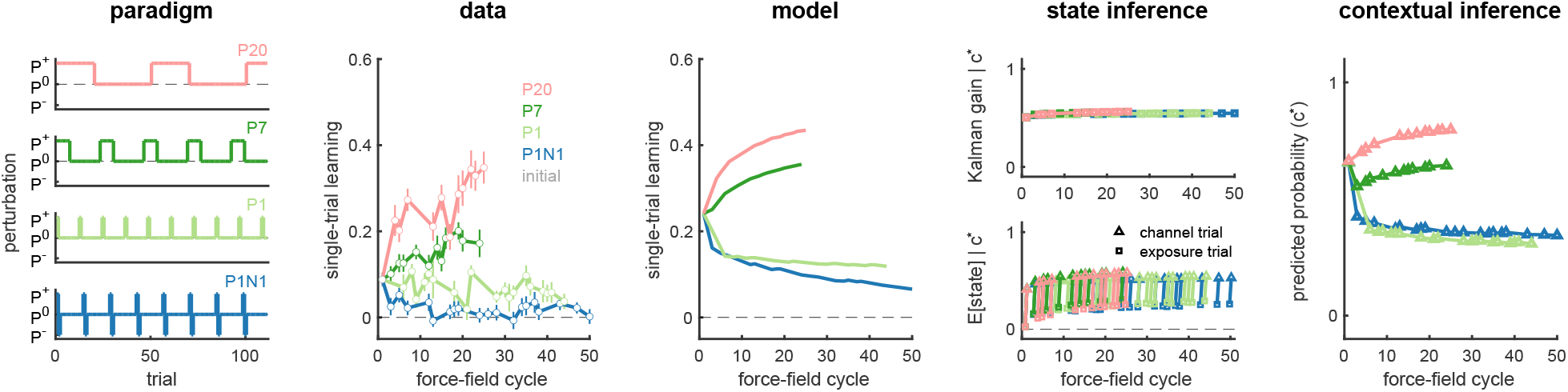
Effect of environmental consistency on single-trial learning. Columns 1 & 2: experimental paradigm and data replotted from Gonzalez Castro et al. ^12^. Columns 3 to 5 show the output and internal inferences of the COIN model in the same format as Fig. 4c.

**Figure S15.**
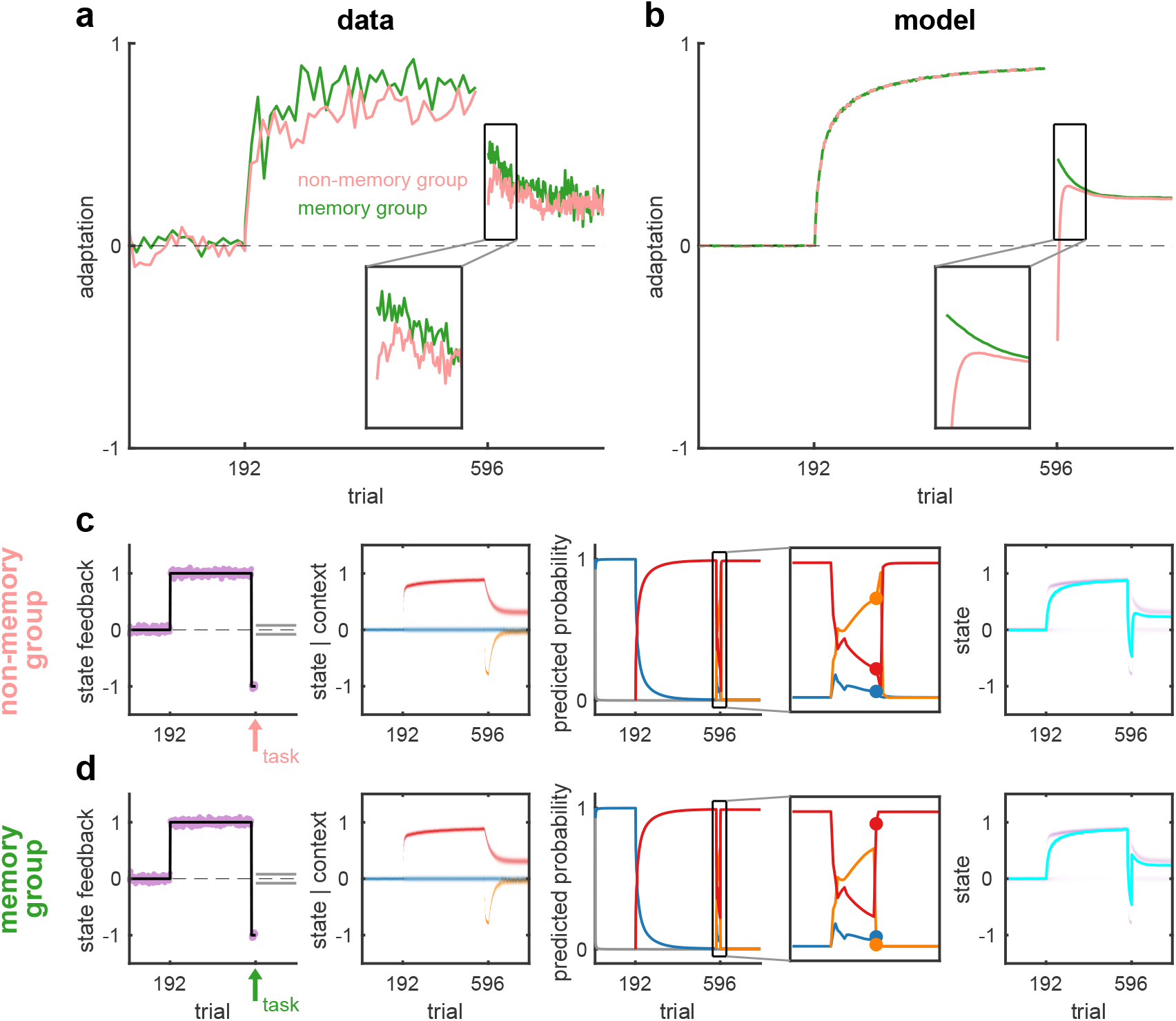
Maintenance of context probabilities may require working memory. **a**, Adaptation in a spontaneous recovery paradigm in which a non-memory (pink) or working memory task (green) is performed before starting the channel trial phase (data reproduced from Keisler and Shadmehr ^19^). Initial adaptation in the channel trial phase (inset) shows the working memory task abolishes spontaneous recovery and leads to adaptation akin to evoked recovery (cf. Fig. S5). **b**, COIN model simulation in which the working memory task abolishes the memory of the context at the end of the *P^−^* phase. **c**, COIN model state feedback, state of each context, predicted probabilities and state output for the non-memory task. The circles on the predicted probability (zoomed view) show the values on the first trial in the channel phase. **d**, as (**c**) for the working memory task. The predicted probabilities on the first trial in the channel phase are the values under the stationary distribution.

**Figure S16.**
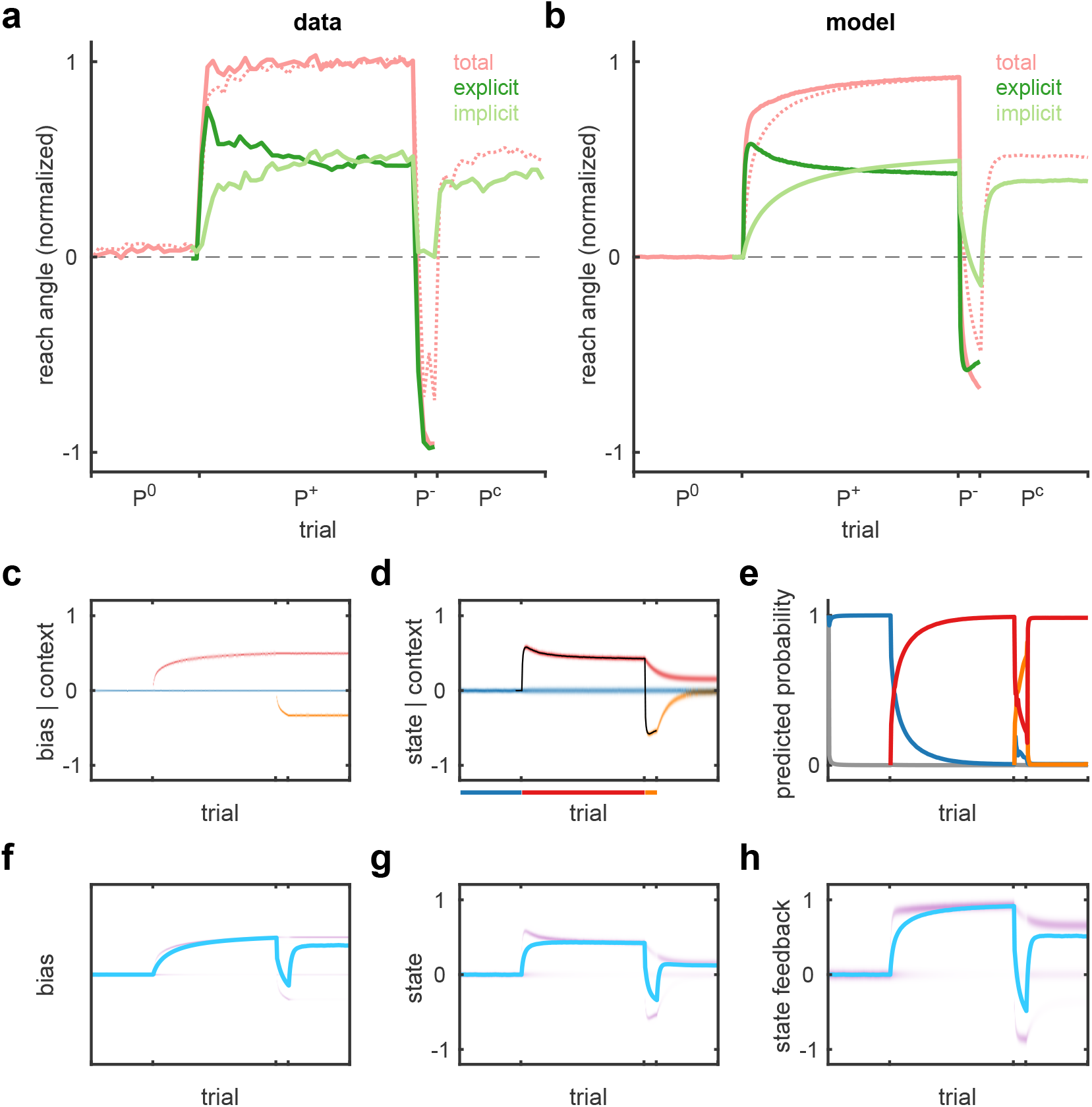
Explicit versus implicit learning in the COIN model. **a,** Results of a spontaneous recovery paradigm (as in Fig. 2b) for visuomotor learning. Explicit learning (dark green) is measured by participants indicating their intended reach direction. Implicit learning (light green) is obtained as the difference between total adaptation (solid pink) and explicit learning. In the visual error-clamp phase (*P*^c^), participants were told to stop using any aiming strategy so that the direction they moved was taken as the implicit component of learning. A control experiment (dashed pink) was also performed in which there was no reporting of intended reach direction. Reproduced from McDougle et al. ^21^. **b,** Simulation of the COIN model with total adaptation (solid pink) reflecting the sum of most probable state from the previous trial (dark green) and estimate of bias (light green). Dashed pink line shows total adaptation in the COIN model for a non-reporting condition. **c**-**h** show simulation of the COIN model. **c**-**e,** Inferred bias (**c**), state, with black line showing the state of the most probable context (colored line below axis) (**d**) and predicted probability (**e**) of each context. **f**-**h,** Predicted (across-context average, as in Fig. 1f) distributions (purple) of bias (**f**), state (**g**), and state feedback (**h**, sum of bias and state), and their means (cyan lines).

**Table S1.** COIN model parameters. Parameters for simulation shown in Fig. 1 (F), model validation (V) and fits for the spontaneous (S) and evoked (E) recovery participants, to the average of both groups (A), and the memory-updating participants (M). Participants in the S and E groups are ordered by decreasing BIC difference between the dual-rate and COIN model (i.e. S1’s and E1’s data most favor the COIN model). Participants in the M group are ordered by decreasing probability of their adaptation data under the COIN model (likelihood).

| participant | $\sigma_q$ | $\mu_a$ | $\sigma_a$ | $\sigma_d$ | $\alpha$ | $\rho$ | $\sigma_m$ | $\alpha^e$ |
| --- | --- | --- | --- | --- | --- | --- | --- | --- |
| F | 0.0500 | 0.9000 | 1.0e-02 | 1.0e-02 | 5.0e+01 | 0.1000 | 0.0500 | 1.0e-01 |
| V | 0.1000 | 0.9000 | 1.0e-01 | 1.0e-01 | 1.0e+01 | 0.9000 | – | 1.0e+01 |
| S1 | 0.1591 | 0.7996 | 6.9e-01 | 1.4e-04 | 3.1e+01 | 0.3297 | 0.0641 | – |
| S2 | 0.1260 | 0.9488 | 8.1e-04 | 6.2e-03 | 2.2e+01 | 0.0869 | 0.0661 | – |
| S3 | 0.0331 | 0.9308 | 4.2e-03 | 2.8e-05 | 1.1e+05 | 0.1766 | 0.0846 | – |
| S4 | 0.1237 | 0.9561 | 1.5e-05 | 9.1e-03 | 7.1e+00 | 0.0208 | 0.0836 | – |
| S5 | 0.0668 | 0.9427 | 1.2e-05 | 4.7e-05 | 1.8e+03 | 0.4522 | 0.0961 | – |
| S6 | 0.0775 | 0.8480 | 1.6e-05 | 6.4e-03 | 1.9e-03 | 0.9999 | 0.0978 | – |
| S7 | 0.1026 | 0.8088 | 1.1e-02 | 1.0e-03 | 8.9e+00 | 0.0016 | 0.1060 | – |
| S8 | 0.0392 | 0.7857 | 1.0e-02 | 5.9e-03 | 2.4e+00 | 0.0152 | 0.0929 | – |
| E1 | 0.1234 | 0.8827 | 1.8e-02 | 7.6e-03 | 1.2e+02 | 0.2699 | 0.0901 | – |
| E2 | 0.1441 | 0.9121 | 3.0e-04 | 8.7e-03 | 3.8e+02 | 0.1285 | 0.0804 | – |
| E3 | 0.1119 | 0.9466 | 6.0e-05 | 7.5e-03 | 2.3e+02 | 0.0070 | 0.0781 | – |
| E4 | 0.1172 | 0.8578 | 1.2e-02 | 5.9e-03 | 1.2e+01 | 0.0477 | 0.0883 | – |
| E5 | 0.0705 | 0.9460 | 1.3e-03 | 5.1e-03 | 1.0e+07 | 0.3911 | 0.0715 | – |
| E6 | 0.0559 | 0.9643 | 3.0e-03 | 4.9e-03 | 2.9e+01 | 0.0044 | 0.0966 | – |
| E7 | 0.0926 | 0.9422 | 1.6e-05 | 3.3e-03 | 3.7e+06 | 0.0003 | 0.0936 | – |
| E8 | 0.0990 | 0.8405 | 2.3e-03 | 7.1e-03 | 1.6e+02 | 0.5336 | 0.1000 | – |
| A | 0.0089 | 0.9425 | 1.2e-03 | 8.2e-04 | 9.0e+00 | 0.2501 | 0.0182 | – |
| M1 | 0.0360 | 0.6575 | 1.1e-03 | 1.0e-05 | 2.4e+03 | 0.0029 | 0.0865 | 1.9e+01 |
| M2 | 0.0412 | 0.7274 | 8.9e-03 | 1.3e-02 | 1.3e-01 | 0.0062 | 0.0921 | 3.0e+03 |
| M3 | 0.0155 | 0.9473 | 1.0e-03 | 7.9e-04 | 8.3e-03 | 0.0003 | 0.0979 | 1.8e+03 |
| M4 | 0.0572 | 0.6986 | 4.8e-03 | 8.0e-03 | 2.8e+04 | 0.1732 | 0.1056 | 1.3e+00 |
| M5 | 0.0574 | 0.8420 | 9.5e-05 | 5.0e-04 | 2.2e+06 | 0.0352 | 0.1187 | 5.9e+00 |
| M6 | 0.0938 | 0.7309 | 1.1e-03 | 1.5e-03 | 2.2e+04 | 0.0118 | 0.1153 | 1.5e+01 |
| M7 | 0.0594 | 0.8744 | 2.4e-05 | 2.3e-03 | 5.2e+06 | 0.0007 | 0.1245 | 9.9e+01 |
| M8 | 0.0708 | 0.7676 | 9.6e-05 | 1.6e-03 | 3.1e+06 | 0.0040 | 0.1206 | 7.0e+00 |
| M9 | 0.0586 | 0.8751 | 2.8e-04 | 1.9e-03 | 1.9e+06 | 0.0139 | 0.1206 | 8.2e+01 |
| M10 | 0.0589 | 0.8949 | 1.3e-04 | 7.3e-04 | 3.4e+05 | 0.1347 | 0.1258 | 2.4e-01 |
| M11 | 0.0419 | 0.9047 | 1.1e-05 | 5.4e-04 | 7.1e+01 | 0.2443 | 0.1262 | 3.5e-01 |
| M12 | 0.0576 | 0.8587 | 1.5e-04 | 1.1e-03 | 4.6e+03 | 0.2205 | 0.1333 | 2.3e+00 |
| M13 | 0.0560 | 0.9164 | 3.3e-05 | 1.4e-03 | 1.8e+01 | 0.1020 | 0.1366 | 9.7e-01 |
| M14 | 0.0364 | 0.8833 | 6.1e-05 | 4.4e-03 | 4.9e+02 | 0.0041 | 0.1410 | 4.0e-01 |
| M15 | 0.0543 | 0.8100 | 4.4e-03 | 8.3e-04 | 6.8e-03 | 0.0508 | 0.1433 | 7.5e-03 |
| M16 | 0.0546 | 0.8249 | 3.0e-05 | 1.1e-03 | 2.7e+04 | 0.3736 | 0.1423 | 1.2e-02 |
| M17 | 0.0485 | 0.8534 | 1.2e-03 | 4.9e-04 | 1.4e+05 | 0.0019 | 0.1487 | 6.0e+01 |
| M18 | 0.0586 | 0.7947 | 5.3e-03 | 1.3e-02 | 1.2e+03 | 0.0002 | 0.1542 | 5.8e-01 |
| M19 | 0.1233 | 0.7211 | 1.3e-04 | 4.4e-03 | 4.1e+03 | 0.0002 | 0.1610 | 3.5e+01 |
| M20 | 0.0210 | 0.9711 | 1.6e-04 | 8.6e-04 | 2.9e-02 | 0.0004 | 0.1534 | 6.6e+02 |
| M21 | 0.0976 | 0.8624 | 4.1e-04 | 8.4e-03 | 9.6e+06 | 0.1154 | 0.1464 | 1.8e+01 |
| M22 | 0.1091 | 0.7853 | 7.6e-04 | 3.9e-03 | 2.1e+03 | 0.0314 | 0.1644 | 1.0e+00 |
| M23 | 0.0708 | 0.8501 | 3.0e-03 | 1.9e-03 | 1.5e+02 | 0.0220 | 0.1905 | 3.5e+01 |
| M24 | 0.0898 | 0.9185 | 1.8e-04 | 3.0e-03 | 6.8e+00 | 0.2249 | 0.1989 | 1.5e+00 |

**Table S2.** Dual-rate model parameter fits for spontaneous and evoked recovery group. Same format as Table S1.

| participant | $a_f$ | $a_s$ | $b_f$ | $b_s$ | $\sigma_m$ |
| --- | --- | --- | --- | --- | --- |
| S1 | 0.5284 | 0.9956 | 0.5108 | 0.0333 | 0.1043 |
| S2 | 0.4194 | 0.9973 | 0.4014 | 0.0427 | 0.0784 |
| S3 | 0.1517 | 0.9879 | 0.1476 | 0.0263 | 0.0984 |
| S4 | 0.7078 | 0.9990 | 0.3869 | 0.0388 | 0.0987 |
| S5 | 0.2565 | 0.9936 | 0.2436 | 0.0497 | 0.1029 |
| S6 | 0.8818 | 1.0000 | 0.3705 | 0.0029 | 0.1043 |
| S7 | 0.5435 | 0.9890 | 0.3407 | 0.0492 | 0.1055 |
| S8 | 0.9296 | 0.9996 | 0.0426 | 0.0306 | 0.0872 |
| E1 | 0.6311 | 0.9984 | 0.4835 | 0.0307 | 0.1448 |
| E2 | 0.4933 | 0.9972 | 0.4524 | 0.0336 | 0.1257 |
| E3 | 0.6069 | 0.9967 | 0.2859 | 0.0331 | 0.0994 |
| E4 | 0.7525 | 0.9947 | 0.5218 | 0.0482 | 0.1081 |
| E5 | 0.6474 | 0.9972 | 0.2409 | 0.0341 | 0.0918 |
| E6 | 0.7642 | 0.9969 | 0.2859 | 0.0366 | 0.1059 |
| E7 | 0.7819 | 0.9901 | 0.1251 | 0.0398 | 0.0895 |
| E8 | 0.7984 | 0.9959 | 0.2233 | 0.0299 | 0.0829 |

**Table S3.** Comparison of COIN to other models. Table shows which experimental phenomena (rows) can be explained by different single and multiple-context models (columns). Alphabetical superscripts index the key feature(s) missing from each model which are primarily responsible for their inability to explain a particular phenomenon. Note that for many models it is either not possible or not clear how the feature(s) could be added. Orange cross-ticks are for models that can partially explain a phenomemon. **Spontaneous recovery**, the gradual, non-monotonic re-expression of *P*^+^ in the channel trial phase (Fig. 2c), requires a single-context model to have multiple states that decay on different timescales or a multiple-context model that can change the expression of memories in a gradual manner based on the amount of experience with each context. Therefore, single-context models that have a single state^**a**^, or multiple-context models that do not learn context transition probabilities^**b**^ or do not have state dynamics^**d**^ do not show spontaneous recovery. Models that learn transition probabilities but that do not represent uncertainty about the previous context^**c**^ (the “local” approximation in DP-KF) can either include a self-transition bias or not. With a self-transition bias, the expression of memories changes in an abrupt manner when, in the channel trial phase, the belief about the previous context changes (e.g. from *P^−^* to *P*^+^), and thus such models fail to explain the gradual nature of spontaneous recovery. Without a self-transition bias, the change in expression of memories is gradual based on updated context counts, but this occurs too slowly relative to the timescale on which the rise of spontaneous recovery occurs. **Evoked recovery**, the rapid re-expression of the memory of *P*^+^ in the channel trial phase (Fig. 2e) that does not simply decay exponentially to baseline (Fig. S6), requires a model to be able to switch between different memories based on state feedback. Therefore, single-context models^**e**^ that cannot switch between memories are unable to show the evoked recovery pattern seen in the data. Multiple-context models with memories that decay exponentially to zero^**f**^ can only partially explain evoked recovery, showing the initial evocation but not the subsequent change in adaptation over the channel trial phase. Models with no state decay^**d**^ cannot explain evoked recovery. **Memory updating** requires a model to update memories in a graded fashion and to use sensory cues to compute these graded updates. Therefore, models that either have no concept of sensory cues^**g**^ or multiple-context models that only update the state of the most probable context in an all-or-none manner^**h**^ do not show graded memory updating. **Savings**, faster learning during re-exposure compared to initial exposure, after full washout requires a single-context model to increase its learning rate or a multiple-context model to protect its memories from washout and/or learn context transition probabilities. Therefore, single-context models with fixed learning rates^**i**^ do not show savings. **Anterograde interference**, increasing exposure to *P*^+^ leads to slower subsequent adaptation to *P^−^*, requires a single-context model to learn on multiple timescales or a multiple-context model to learn transition probabilities that generalize across contexts. Therefore, single-context models with a single state^**a**^, or multiple-context models that either do not learn transition probabilities^**b**^ or that learn transition probabilities independently for each row of the context transition matrix^**j**^ do not show anterograde interference. **Environmental consistency**, the increase/decrease in single-trial learning for slowly/rapidly switching environments, requires a model to either adapt its learning rate or learn context transition probabilities based on context transition counts. Therefore, single-context models with fixed learning rates^**i**^ or multiple-context models that either do not learn transition probabilities^**b**^ or that learn transition probabilities based only on context counts^**k**^ do not show the effects of environmental consistency on single-trial learning. **Explicit and implicit learning**, the decomposition of visuomotor learning into explicit and implicit components, requires a model to have elements that can be mapped onto these components. For most models, there is no clear way to map model elements onto these components^**l**^. It has been suggested that the fast and slow processes of the dual-rate model correspond to the explicit and implicit components of learning, respectively. However, in a spontaneous recovery paradigm, this mapping only holds during initial exposure and fails to account for the time course of the implicit component during the counter-exposure and channel trial phases^**m**^ (see Suppl. Text).

|  | single-context models |  | multiple-context models |  |  |  |  |
| --- | --- | --- | --- | --- | --- | --- | --- |
|  | dual-rate<br>Smith et al. <sup>3</sup> | memory<br>of errors<br>Herzfeld et al. <sup>9</sup> | source of<br>errors<br>Berniker and<br>Kording <sup>8</sup> | winner-<br>take-all<br>Oh and<br>Schweighofer <sup>2</sup> | DP-KF<br>Gershman et al. <sup>6</sup> | MOSAIC<br>Haruno et al. <sup>23</sup> | COIN |
| spontaneous recovery | ✓ | ✗<sup>a</sup> | ✗<sup>b</sup> | ✗<sup>b</sup> | ✗<sup>c</sup> | ✗<sup>d</sup> | ✓ |
| evoked recovery | ✗<sup>e</sup> | ✗<sup>e</sup> | ✗<sup>f</sup> | ✗<sup>f</sup> | ✗<sup>f</sup> | ✗<sup>d</sup> | ✓ |
| memory updating | ✗<sup>g</sup> | ✗<sup>g</sup> | ✗<sup>g</sup> | ✗<sup>h</sup> | ✗<sup>g,h</sup> | ✓ | ✓ |
| savings after full washout | ✗<sup>i</sup> | ✓ | ✓ | ✓ | ✓ | ✓ | ✓ |
| anterograde interference | ✓ | ✗<sup>a</sup> | ✗<sup>b</sup> | ✗<sup>b</sup> | ✓ | ✗<sup>j</sup> | ✓ |
| environmental consistency | ✗<sup>i</sup> | ✓ | ✗<sup>b</sup> | ✗<sup>b</sup> | ✗<sup>k</sup> | ✓ | ✓ |
| explicit/implicit learning | ✗<sup>m</sup> | ✗<sup>l</sup> | ✗<sup>l</sup> | ✗<sup>l</sup> | ✗<sup>l</sup> | ✗<sup>l</sup> | ✓ |

